# Patterns and Drivers of Diatom Diversity and Biogeography in the North Pacific

**DOI:** 10.64898/2026.08.30.746603

**Authors:** Amaury Barral, Koji Suzuki, Yukie Kikuchi, Shin-ichiro Nakaoka, Shintaro Takao, Shinji Nakaoka

## Abstract

Marine diatoms contribute to about 20% of global primary production. We present the first basin-scale, multiyear assessment of diatom communities in the North Pacific, combining taxonomically high-resolution RuBisCO large subunit gene (*rbcL*) metabarcoding with concurrent environmental measurements. Using a nine-year time series of daily samples resolved at the species level via ∼500 bp *rbcL* fragments, we performed multivariate analyses across biogeographic provinces, identifying significant correlations between community structure and environmental drivers such as temperature and macronutrient availability.

We report the prevalence of a previously overlooked centric diatom species in the North Pacific, *Eunotogramma lunatum*, which appears to be near-dominant even in subarctic high-nitrate, low-chlorophyll waters where pennate diatoms are typically favored.

These results demonstrate the power of *rbcL* for large-scale ocean monitoring and provide a critical baseline for future studies of diatom population dynamics, climate change impacts, and ecosystem resilience in a key marine region.

**Significance:** Diatoms are among the ocean’s most important primary producers, yet large-scale, long-term assessments of their diversity have been limited by the resolution of traditional monitoring methods. This study provides the first basin-scale, multiyear baseline of diatom community structure across the North Pacific, demonstrating that genetic metabarcoding can resolve species-level diversity at a scale and resolution previously unattainable. By linking nine years of daily samples to concurrent environmental data, we show that temperature and nutrient availability are among the key predictors of community composition across distinct biogeographic regions. These findings establish a critical reference point for tracking how diatom communities – and the ecological services they underpin, including carbon export and marine food web support – may shift in response to ongoing climate change.

---

Diatoms represent the most abundant and diverse group of eukaryotic phytoplankton, comprising tens of thousands of species that span a broad size spectrum (1, 2). They are fundamental to marine ecosystem functioning, believed to be responsible for approximately 20% of the net primary production on Earth and acting as the base of aquatic food webs (3). Through the formation of rigid silica frustules, diatoms are the world’s largest contributors to biosilicification and play a critical role in the export of carbon from the surface ocean to depth (4, 5) through the ocean biological pump. Consequently, understanding the environmental drivers of diatom diversity and spatial distribution is essential for predicting the resilience of marine ecosystems to natural and human-induced perturbations (6).

The North Pacific Ocean features significant oceanographic transitions, creating distinct regimes that differentially favor diatom populations. North of the Subarctic Front, deep winter mixing resupplies macronutrients, yet chronic iron limitation sustains high-nutrient (nitrate), low chlorophyll (HNLC) dynamics (7, 8). Massive blooms of large, chain-forming diatoms can be triggered when this limitation is relieved by iron supply from atmospheric or lateral sources from marginal seas (9–11). Southward, the North Pacific Subtropical Gyre (NPSG) maintains year-round water-column stratification and ultra-oligotrophic surface waters where picoplankton account for most standing biomass (6), yet diatoms contribute disproportionately to episodic carbon export events linked to mesoscale eddies, atmospheric deposition, or transient mixing (12–14). Between these contrasting regimes, the Transition Zone and the Kuroshio-Oyashio Extension host dynamic frontal systems where complex interactions between basin-scale circulation and mesoscale processes generate considerable spatial and temporal heterogeneity in nutrient supply and plankton communities (15, 16). Despite decades of observations at time-series stations such as ALOHA and K2 (17, 18), systematic assessments of diatom taxonomic diversity across the entire North Pacific remain limited.

This knowledge gap is particularly acute outside of established subarctic and marginal hotspots, leaving the seasonal and interannual community dynamics of the vast open-ocean regions poorly understood (19, 20).

Efforts to resolve these biogeographic patterns have historically been hindered by methodological limitations. Traditional routine monitoring relies on microscopy, which, while valuable, is observer-dependent and often fails to discriminate cryptic species, small-celled taxa or variable cellular morphologies (21).

In recent years, metabarcoding has emerged as a scalable alternative, allowing for the detection of rare and cryptic taxa that microscopy may overlook (22). Largescale surveys, such as the Tara Oceans expedition, have utilized the V9 region of the 18S rRNA gene to map global plankton diversity (23, 24). However, the 18S marker frequently lacks sufficient variability for species-level resolution in diatoms, even when longer fragments are sequenced (25, 26) - a significant limitation given that ecologically distinct populations often diverge below the genus level (27).

To overcome these limitations, the plastid-encoded RuBisCO large subunit (*rbcL*) gene has been proposed as a more suitable marker for diatom metabarcoding (28–30). Although *rbcL* reference databases are less comprehensive than those for 18S, recent curation efforts have substantially improved taxonomic coverage for marine diatoms, making basin-scale surveys increasingly feasible (31). Studies comparing *rbcL* to 18S have shown that *rbcL* offers superior barcoding suitability, particularly for resolving populations of ecologically important genera such as *Pseudo-nitzschia* and *Chaetoceros* (32). Although *rbcL* has been used in freshwater and local marine studies, its application to basin-scale marine monitoring remains limited.

In this study, we present a multiyear assessment of diatom diversity and biogeography across the North Pacific. While conventional *rbcL* metabarcoding studies typically target a 260 bp region, we utilized an extended 500 bp fragment to achieve substantially higher taxonomic resolution and paired this molecular dataset with concurrent environmental measurements. While correlative analyses cannot establish underlying mechanisms, this work provides an informed baseline for understanding diatom biogeography in a critically important but understudied ocean basin.

## 1. Results

We analyzed diatom communities in the North Pacific accross eight biogeochemical Longhurst provinces ^∗^ (33, 34): Eastern/Western Pacific subarctic gyres (PSAE/PSAW), Kuroshio current (KURO), North Pacific polar front (NPPF), Northwest Pacific subtropical (NPSW), North Pacific Tropical gyre (NTG), California current (CCAL), North Pacific equatorial counter current (PNEC). Figure 1 shows the boundary of each province as well as the spatiotemporal distribution of our 1392 samples spanning nine years (2014/09-2023/04). The number of samples per location is displayed in Supplementary S3.

**Fig. 1.**
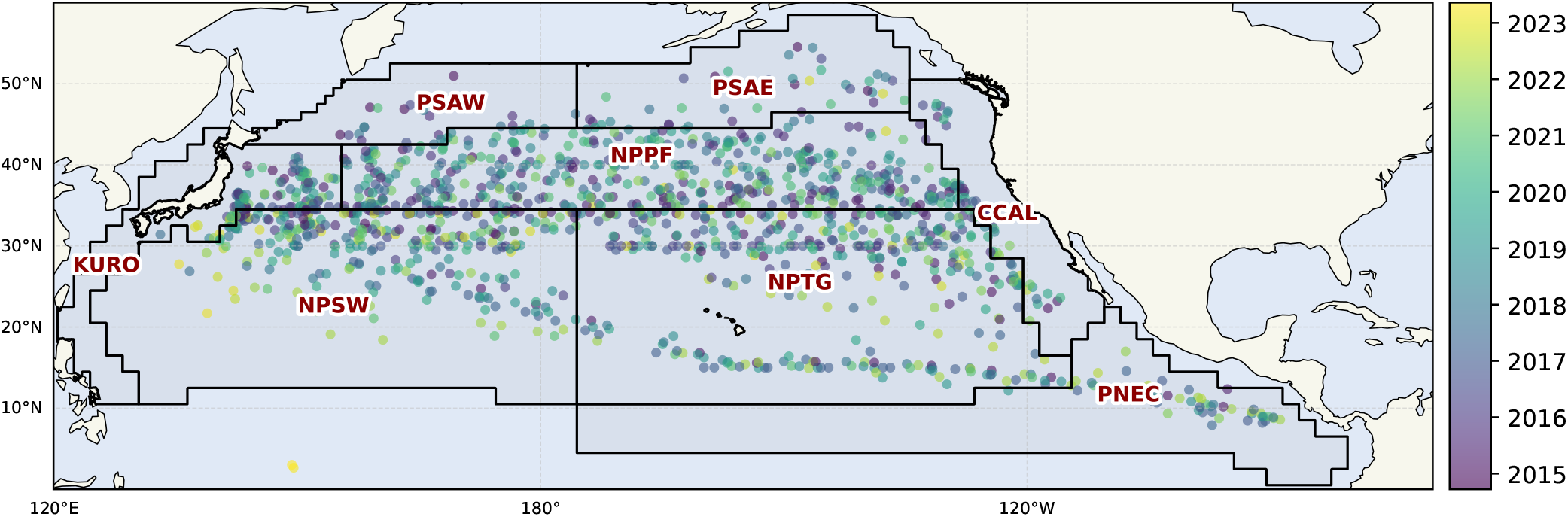
Spatiotemporal distribution of the 1392 sampling sites. Samples are color-coded by their sampling time. Longhurst provinces of interest are overlaid (black outlines, red names).

### 1.1. Diatom Community Composition

Our dataset comprises 73 million^†^ sequences grouped into 40 050 distinct Amplicon Sequence Variants (ASVs) representing 491 species in 178 genera. *Chaetoceros* was the most abundant genus (21% of total assigned sequences), followed by *Eunotogramma* (17%), *Toxarium* (9%), *Fragilariopsis* (6%), *Bacillaria* (6%) and *Pseudo-nitzschia* (5%) (Supplementary S4).

While some genera (e.g., *Asterionella, Cymbella*) were represented by a single ASV, *Chaetoceros* comprised 11 294 ASVs – accounting for 28% of all ASVs recovered, highlighting pronounced disparity in genetic richness at the genus level (Figure 2).

**Fig. 2.**
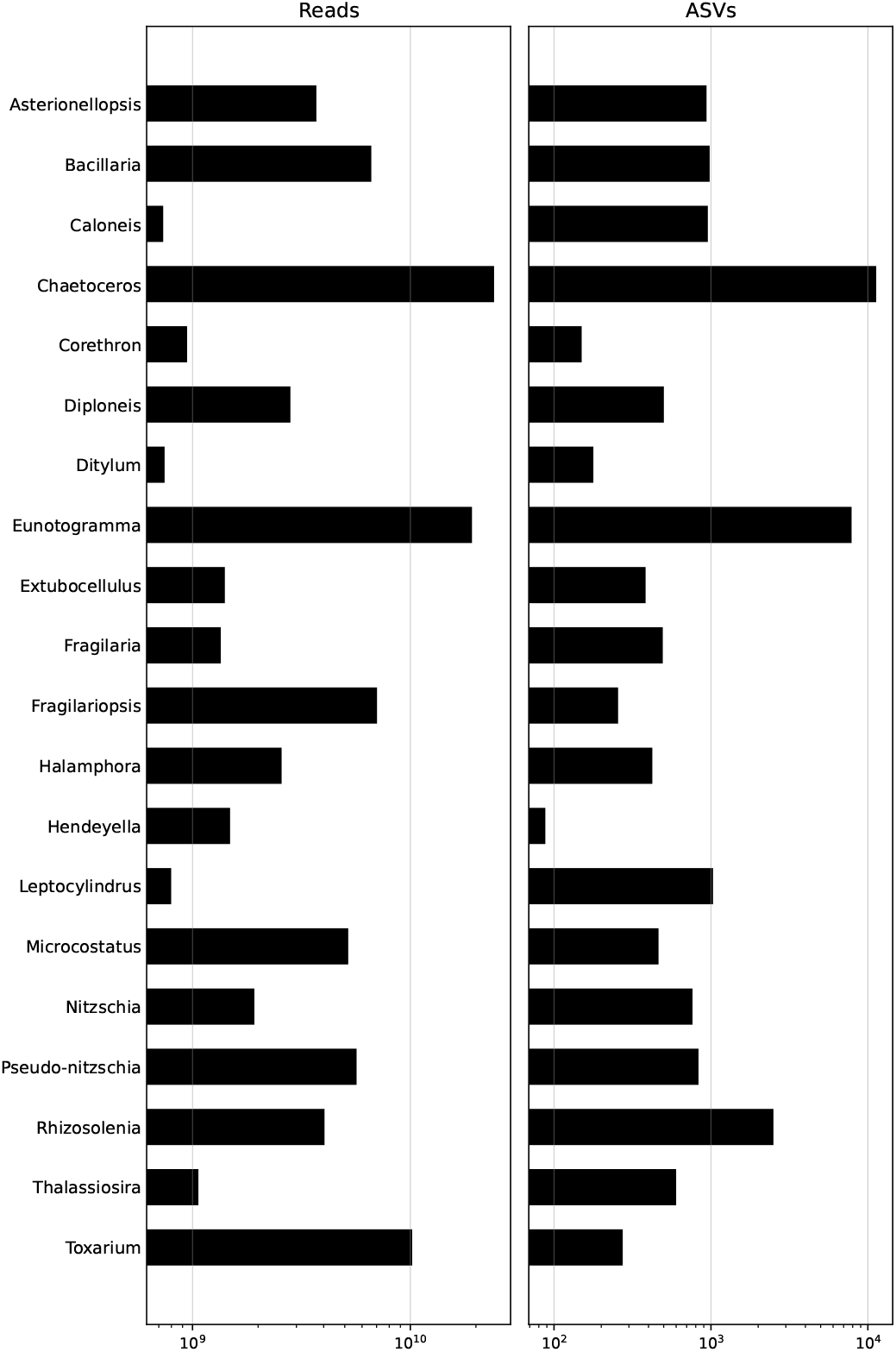
Taxonomic distribution of this study’s dataset: comparison of read abundance and ASV richness by genus for the 20 most abundant genera. The left panel displays the total sequencing reads, while the right panel shows the count of unique Amplicon Sequence Variants (ASVs) assigned to each genus. The full figure with all genera is presented in Supplementary S5.

Surprisingly, the overall community composition is remarkably stable over the seasons (fig. 3), despite some species like *Asterionellopsis glacialis* exhibiting significant fluctuations.

**Fig. 3.**
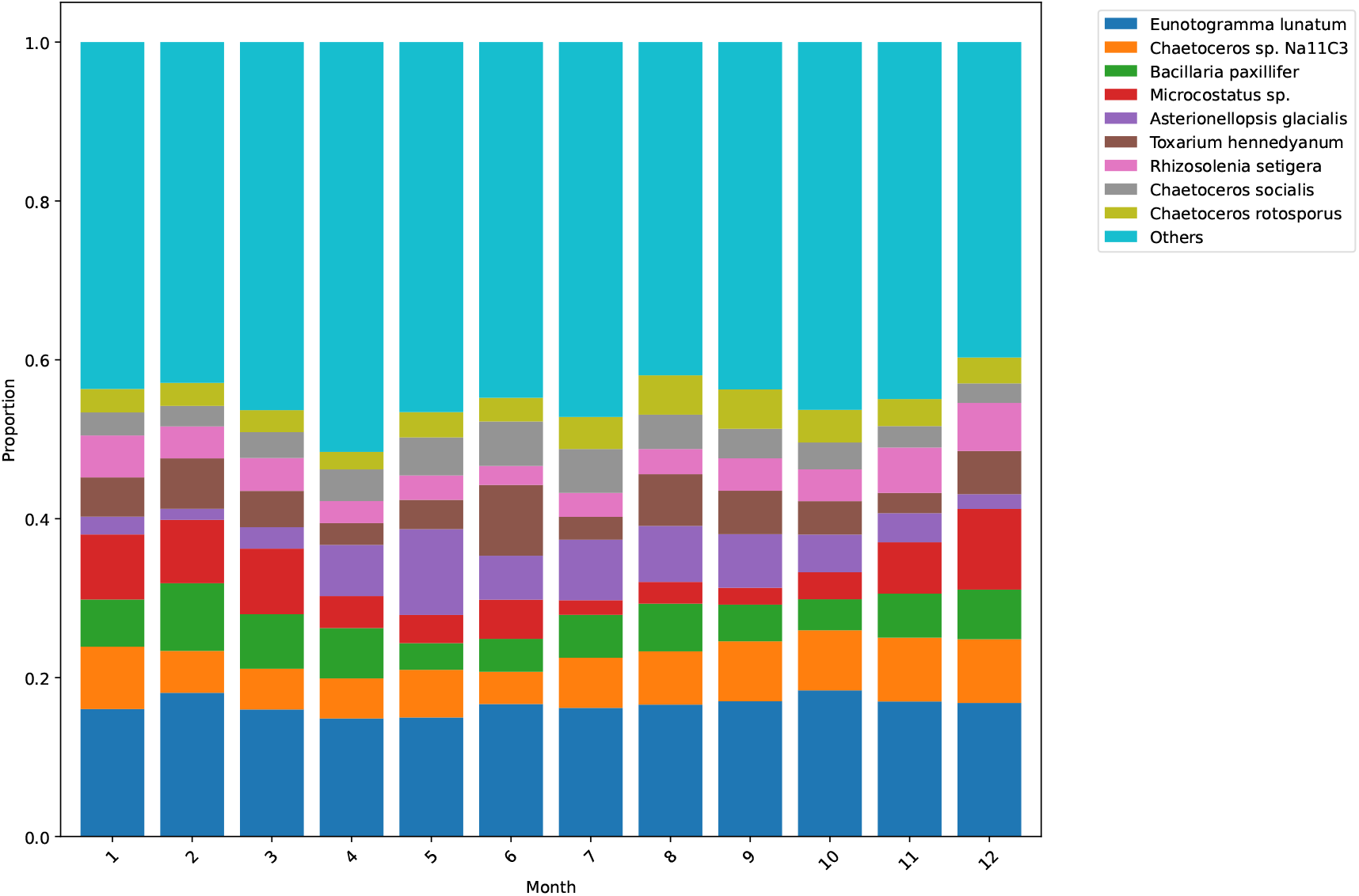
Monthly community composition at the species level. The top nine dominant species are highlighted. The overall composition is remarkably stable.

### 1.2. Diversity patterns

The existence of a latitudinal gradient of diatom diversity is still an active field of research (24, 35). Although our data show such a gradient at mid-latitudes (Figures 4b and 4c), we advise against interpreting this result at face value, as it is evidently biased by sampling unevenness.

**Fig. 4.**
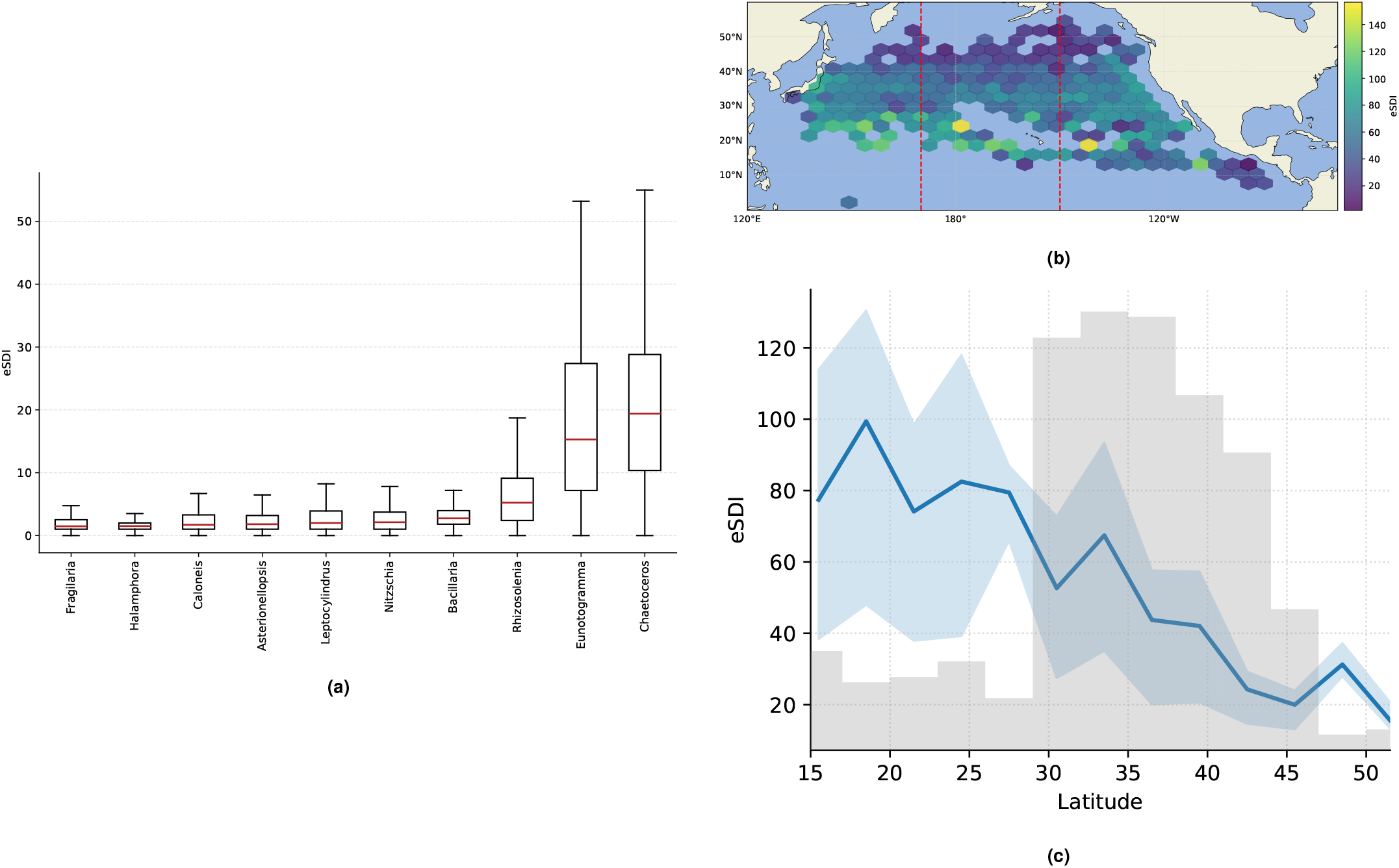
Global patterns of Exponential Shannon Diversity Index (eSDI). (a) Distribution of eSDI (proxy for alpha diversity) for the ten genera with the highest median diversity. (b) Spatial distribution of eSDI; vertical dashed lines demarcate the longitudinal transect (170°–210°) (to reduce coastal effects). (c) Latitudinal gradient of eSDI averaged across the selected transect. The solid line represents the mean, shading indicates the 25-75 interquartile range, and the histogram displays sample density.

The most alpha-diverse^‡^ genera were *Chaetoceros, Eunotogramma, Rhizosolenia* and *Bacillaria* (Figure 4a).

### 1.3. Environmental drivers

Diatom diversity and abundance exhibit clear seasonal patterns (Figure 5a), characterized by distinct species assemblages that are differentially abundant across seasons (Supplementary S8).

**Fig. 5.**
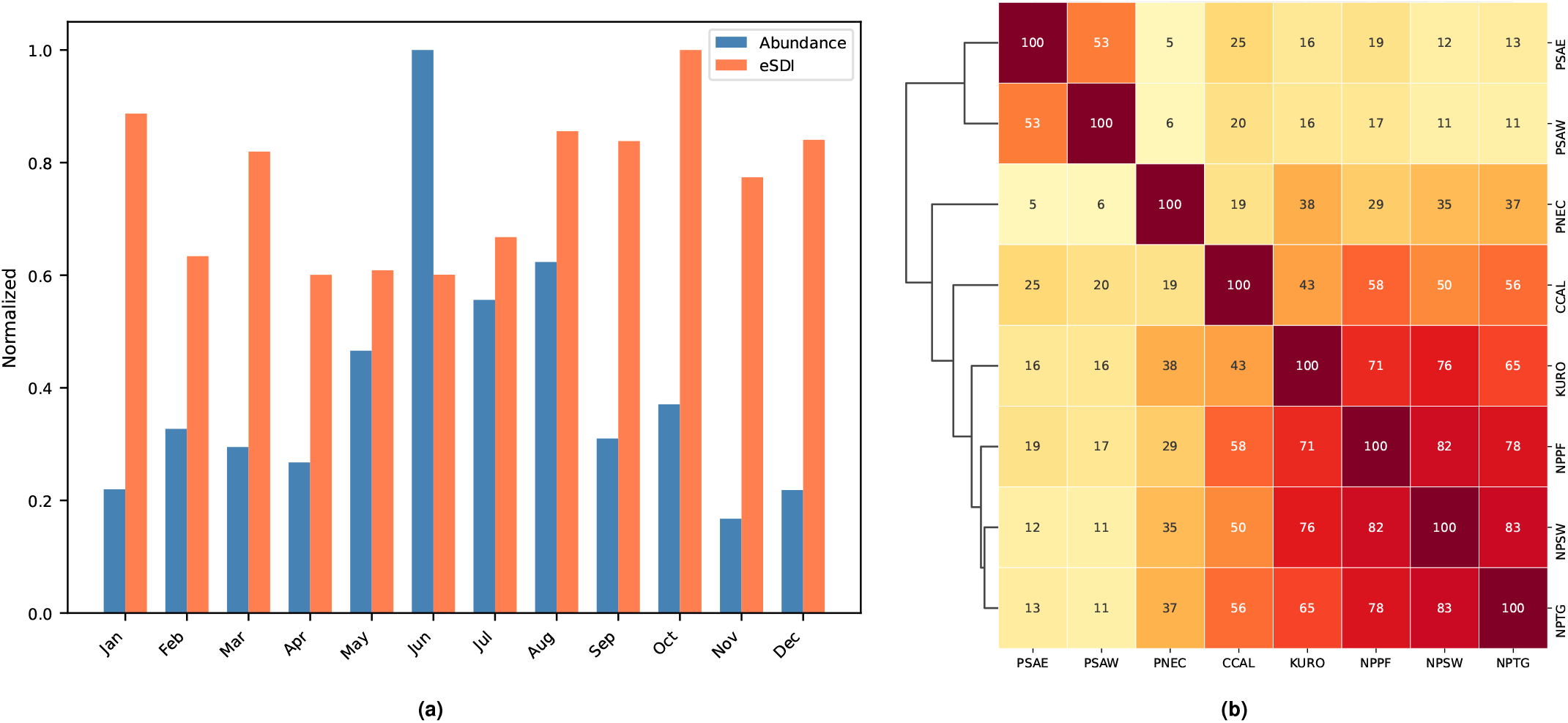
Seasonal and regional distribution of alpha diversity (a) Seasonal patterns of abundance (blue) and exponential Shannon Index (eSDI, orange, proxy for alpha diversity). (b) Pairwise similarity of ASV composition across oceanographic provinces. The heatmap displays the Jaccard similarity index (%) calculated from ASV presence/absence data. The dendrogram indicates complete linkage of regions based on community composition.

The environmental drivers considered include physical properties (distance to coast, sea surface salinity [SSS] and temperature [SST]), carbonate system variables (pH, dissolved inorganic carbon [DIC], and partial pressure of CO_2_ in the atmosphere [ApCO2], sea surface [SpCO2], and the air-sea difference [DpCO2]), nutrient concentrations (ammonium [NH4N], silicate [SiO2Si], nitrite + nitrate [NO2NO2N], phosphate [PO4P], and nitrite [NO2N]), and phytoplankton standing stock (Chlorophyll a).

The relationships and correlations among these drivers are illustrated in Figure 6a. Figure 6b illustrates correlations between environmental drivers and diatom abundance or alpha diversity (eSDI). These results largely align with (35) when considering overlapping drivers but demonstrate enhanced statistical significance, likely due to our larger dataset and reduced spatial scale. Significant negative correlations were observed between silicate and eSDI, and between ammonium (NH_4_^+^) with abundance. In contrast, (35) reported a non-significant NH_4_^+^-abundance correlation and a positive NH_4_^+^-eSDI correlation, whereas our study detected positive NH_4_^+^- abundance and negative NH_4_^+^-eSDI correlations.

**Fig. 6.**
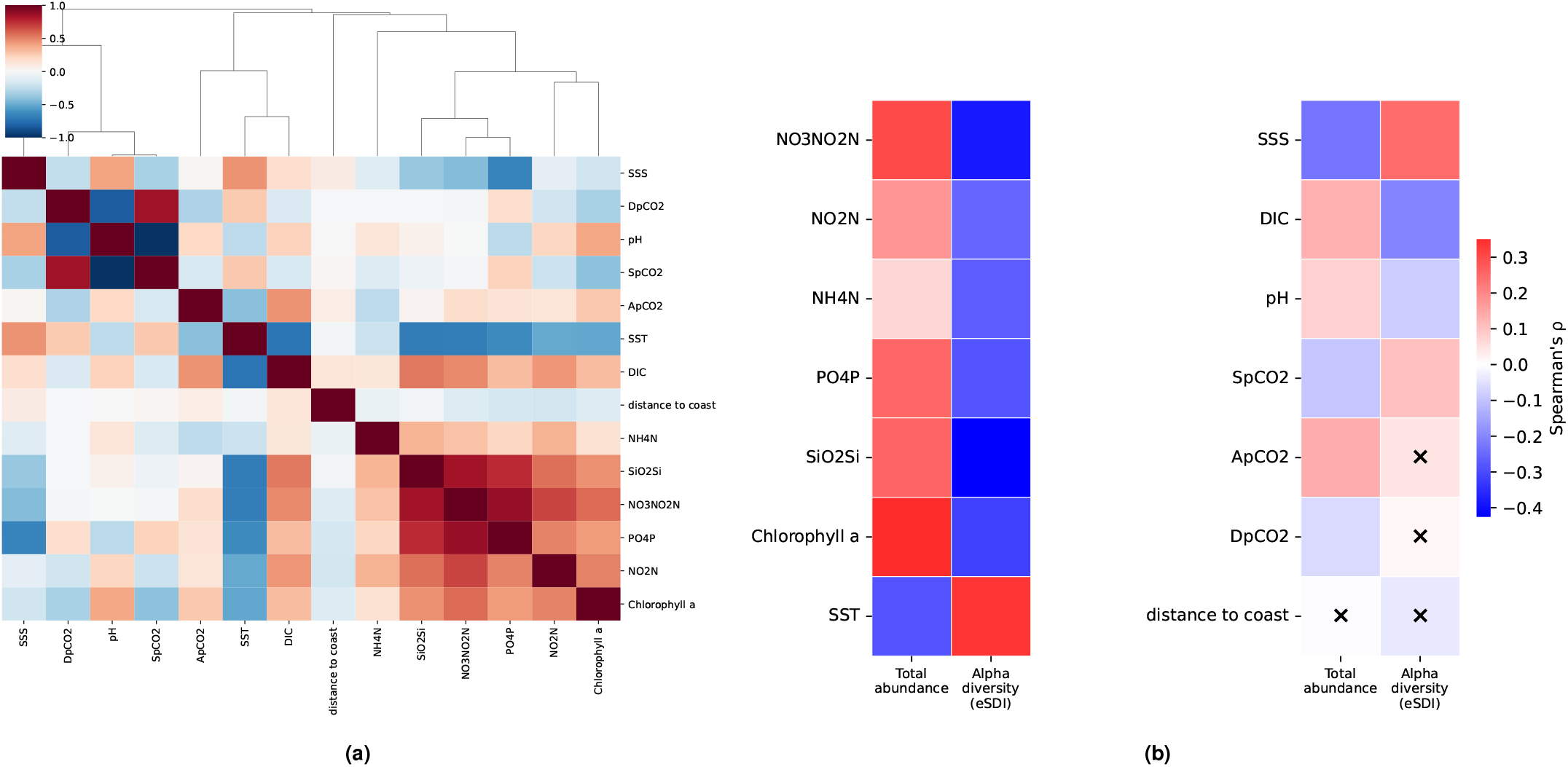
Relationships among environmental drivers and biological metrics. Prior to analysis, skewed variables were log-transformed and all variables were standardized (mean 0, unit variance). (a) Correlation matrix and hierarchical clustering of environmental drivers. Colors represent Spearman correlation coefficients (*ρ*). The dendrogram shows complete linkage clustering based on similarity (1 − |*ρ*^2^|). (b) Spearman’s rho correlations between environmental drivers and abundance / alpha diversity (exponential Shannon Diversity Index). Crosses indicate non-significant correlations (*p >* 0.05).

The redundancy analysis (RDA) of species-level community composition in relation to normalized environmental drivers is presented in Figure 7. Species abundances were Hellinger-transformed prior to analysis. RDA was performed using forward-selected environmental drivers, which were pre-clustered to mitigate multicollinearity and reduce the Variance Inflation Factor (VIF; see Supplementary S6). Forward selection identified that variations in the diatom community are primarily driven by the [SST, DIC] cluster, followed by the [PO4P, SiO2Si, NO3NO2N] cluster. Collectively, the selected environmental drivers account for 76% of the total observed variance.

**Fig. 7.**
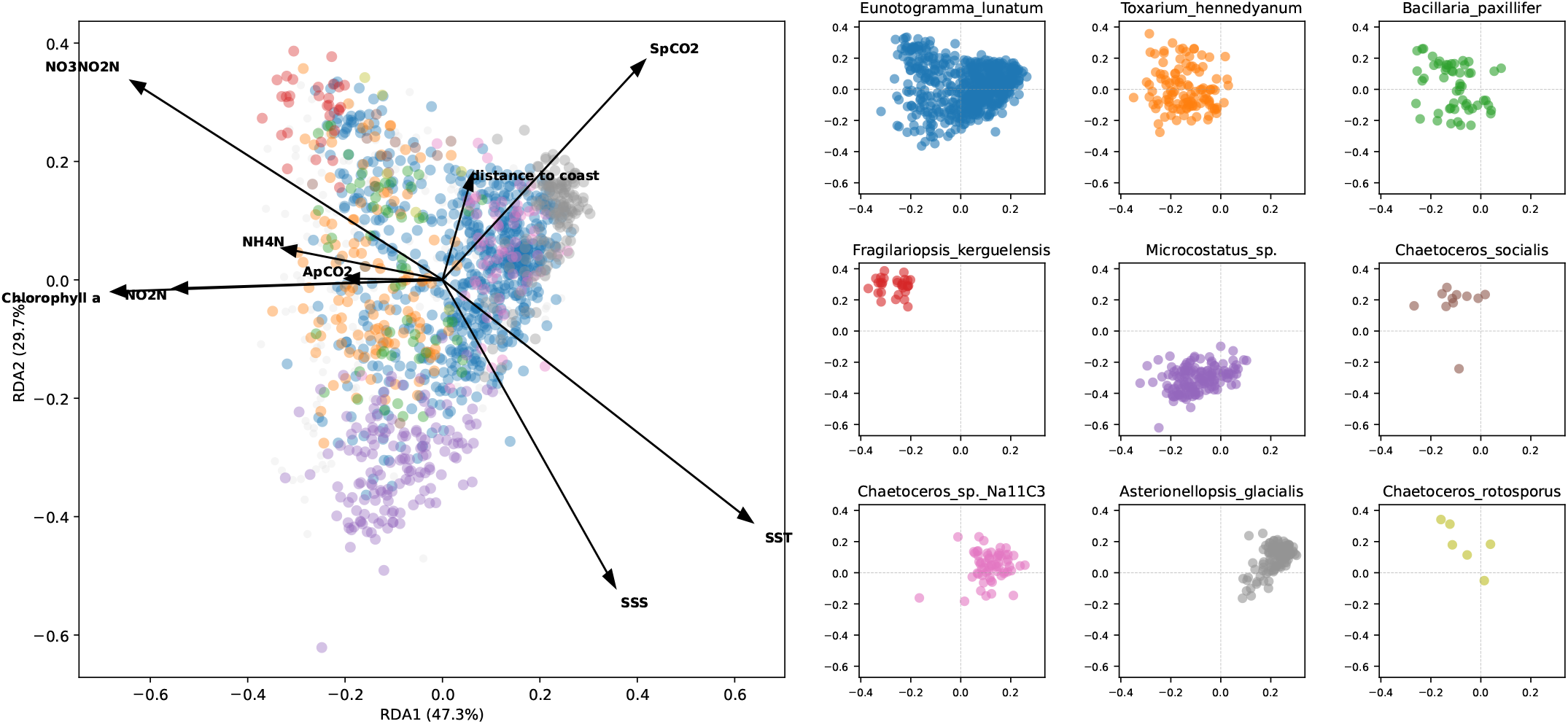
Redundancy Analysis (RDA) of species-level community composition. Abundance counts are Hellinger-transformed. (Left) Biplot of all samples colored by the dominant species. Black arrows indicate significant environmental drivers (*p*_*adj*_ *<* 0.05); samples dominated by species outside the top nine are shown in light grey. Axes indicate the percentage of variance explained. (Right) Individual panels displaying the distribution of the nine most abundant species projected onto the same RDA coordinates.

### 1.4. Discussion

The present study provides a species-level perspective on diatom assemblages across the North Pacific, a region that has been historically underrepresented in high-resolution phytoplankton surveys.

Using metabarcoding data, we identified dominant taxa in each Longhurst province, and linked their distribution patterns to a suite of environmental drivers through RDA. By focusing on the *rbcL* marker, we achieve species-level resolution that exceeds the taxonomic precision of previous studies. As suggested by (36), the most detectable signal of climate-induced change is not total biomass, but the turnover of species composition. By providing a species-resolved baseline using the *rbcL* marker, this study establishes a high-resolution reference point necessary for detecting the community turnover predicted by global ecosystem models.

Overall, the community composition remained surprisingly stable. Although certain species, such as *Asterionellopsis glacialis*, exhibited significant monthly fluctuations, further analysis reveals that these shifts were compensated for by competitors occupying similar ecological niches, such as *Chaetoceros sp. Na11C3*.

RDA identified SST^§^ as a key variable associated with diatom community variation in the North Pacific. Temperature is known to modulate diatom-related biogeochemical fluxes (37, 38). As the North Pacific transition region has experienced accelerated warming (39), such temperature-sensitive processes may increasingly influence diatom-driven carbon and silicate export. Interestingly, we did not observe a clear alpha diversity hotspot in the Kuroshio transition region, contrary to the expectations often associated with such highly dynamic frontal systems.

Unexpectedly, our survey revealed a pronounced dominance of *Eunotogramma lunatum* (EL), a species that has never been identified in prior large-scale diatom studies. EL was only recently described (40) through detailed electron-microscopy analysis and has since been incorporated into the Diat.barcode database following the work of de Oliveira et al. (41). Since its initial discovery in the North Atlantic, this represents, to our knowledge, the first record of EL in the North Pacific. RDA revealed a positive association between SST and the relative abundance of EL, suggesting that warming scenarios may favor this taxon within North Pacific diatom communities. Interestingly, EL persists as a dominant centric diatom in iron-limited subarctic waters, which challenges the prevailing expectation that such conditions favor pennate taxa (42).

Our data set consists of nearly 1 500 samples collected daily over a decade, providing the most extensive and temporally resolved diatom survey in the North Pacific. While most models project a decrease in biomass and alpha diversity under global warming scenarios (43, 44), our ten-year dataset remains insufficient to make statistically significant claims regarding such long-term trends. This high-resolution, long-term archive will be a valuable resource for the broader oceanography and microbial ecology communities. Future analyses could focus on incorporating diatom cell size – a critical trait for understanding nutrient uptake and carbon export (45) – as well as dissolved metal concentrations (especially iron), examine interannual population dynamics, combine in-situ measurements with satellite remote sensing, and investigate in more depth the environmental drivers that shape community shifts across Longhurst provinces.

## Materials and Methods

### 1.5. Dataset generation

#### 1.5.1. Sampling and hydrographic analysis

Seawater samples were collected from a seawater intake at a depth of about 5 m on the bottom of the M/V *New Century 2* (Ocean Link Ltd., Japan) around 2:00 PM (local time) from September 2014 to spring 2023 as part of ship-of-opportunity observations in the North Pacific between Japan and North America, conducted by the National Institute for Environmental Studies (NIES), Japan. Each seawater sample (1.2 L) was filtered onto Millipore Express® PLUS polyether sulfone membrane filters (25 mm in diameter, 0.45 µm pore size) using a gentle vacuum (0.013 MPa). After filtration, the filter samples were stored at -70°C or in liquid nitrogen until analysis on land.

The sea surface temperature (SST) and salinity (SSS) were continuously measured using an SBE 45 MicroTSG Thermosalinograph (Sea-Bird Scientific) in the seawater intake system. Nutrient concentrations were analyzed on land with a Bran + Luebbe auto-analyzer, following Yasunaka et al. (46). The detection limits for nitrate + nitrite, nitrite, ammonium, phosphate, and silicate were 0.01 to 0.13, 0.01 to 0.06, 0.06 to 0.40, 0.02 to 0.05, and 0.04 to 0.70 µM, respectively. Nutrient concentrations below the detection limits were considered zero. Dissolved inorganic nitrogen (DIN) concentration was calculated as the sum of nitrate, nitrite, and ammonium. Chlorophyll a (Chl a) concentration was determined on land by fluorometry (Welshmeyer (47)). Seawater pCO2 and atmospheric pCO2 were measured continuously using the NIES shipboard CO2 measurement system (48). Dissolved inorganic carbon (DIC) was calculated with a seawater carbon calculator (CO2SYS; Lewis et al. (49)), utilizing seawater pCO2 and total alkalinity (TA), which was calculated based on the empirical equations of (50).

#### 1.5.2. DNA extraction and rbcL sequencing

DNA was extracted following the protocol of Endo et al. (51) and purified with a NucleoSpin® gDNA Clean-up kit (Macherey-Nagel) according to the manufacturer’s instructions. Amplicon sequencing was performed by a commercial service provider (Seibutsu Giken Inc., Kanagawa, Japan). DNA concentrations were measured using a Synergy LX microplate reader (Agilent Technologies) with the QuantiFluor dsDNA System (Promega). The *rbcL* gene, which encodes the large subunit of RuBisCO in chromophytic algae, including diatoms, was amplified from the DNA template using the *rbcL* primer set, producing a 554-bp fragment (Wawrik et al. (52)), with KOD FX Neo polymerase (Toyobo Co., Ltd.). The PCR thermal cycling conditions were as follows: an initial denaturation at 94°C for 2 min, followed by 30 cycles of denaturation at 98°C for 10 s, annealing at 52°C for 30 s, and extension at 68°C for 30 s, ending with a final extension at 68°C for 7 min. The PCR products were purified with VAHTS DNA Clean Beads (Vazyme). Sequencing was carried out on an Illumina MiSeq or NextSeq 1000 platform using paired-end 2 × 300 bp reads.

#### 1.5.3. Quantitative PCR for diatoms’ rbcL gene

Real-time PCR amplification of each template was performed on a Bio-Rad CFX Duet using TB Green® Premix Ex Taq™ II (Tli RNaseH Plus), with 0.4 µM of each primer for the diatoms’ rbcL gene, as described by (53). The thermal cycling conditions were 95 °C for 60 s, followed by 40 cycles of 95 °C for 5 s and 52 °C for 60 s. The copy number for each sample was determined using standard curves generated from serial dilutions of the standards, prepared as described by (54).

#### 1.5.4. Bioinformatic processing and dataset curation

Raw paired-end reads were processed using *QIIME 2* (v2025.4) (55). Read sequences were filtered via *Cutadapt* ; only sequences initiating with a primary match to the target primers were extracted, while those shorter than 40 bp or falling below a quality threshold of *Q* = 20 were discarded. The selected reads were then denoised using *DADA2* to resolve Amplicon Sequence Variants (ASVs). Read merging was conducted requiring a minimum overlap of 10 bp. Taxonomic classification was performed using *MMseqs2* against the subset of the *Diat.barcode* (56) database (a curated reference database for diatoms) (`flavor=`“`original`” (57)) targeting the *rbcL* region (32). ASVs were assigned based on the lowest matching E-value, with a threshold of *E* ≤ 10^−5^, after which only members of the *Bacillariophyta* phylum were kept to focus on diatom communities.

The resulting dataset is compositional, which introduces well-documented analytical biases (58, 59). To avoid those, we transformed relative abundances to estimated absolute abundances (up to a global prefactor) using total diatom concentration (determined via qPCR) as a scaling factor.

Concerning the environmental dataset, after basic data curation, we removed all points within 1000 km of the coastline to reduce coastal effects. To address missing values resulting from instrumental or logistical constraints, we imputed the gaps by training a machine-learning model (AutoGluon 1.4 (60), sequential `MultilabelPredictor`) on the corresponding COPERNICUS (61, 62) reanalysis data (variables `thetao_mean,so_mean` from `mems_mod_glo_phy-mnstd_my_0.25deg_P1D-m`; `spco2,fe,ph,phyc` from `cmems_mod_glo_bgc_my_0.25deg_P1M-m`; `chl,no3,nppv,o2,po4,si` from `cmems mod glo bgc my 0.25deg P1D-m`) (Supplementary S1).

This workflow is summarized in Supplementary S2.

## Data, Materials, and Software Availability

The dataset collected in this study, as well as the code used to process and analyze it, will be made Open Source upon publication of the paper. We will update this section with relevant links on publication.

## ACKNOWLEDGMENTS

We gratefully acknowledge the captain, officers, and crew of the *M/V New Century 2*, together with Toyofuji Shipping Co., Ltd. and Ocean Link Ltd., for their generous support and cooperation throughout the ship-of-opportunity observation program. We also wish to express our gratitude to Mr. Tomoyasu Yamada (Global Environmental Forum) for his assistance with the logistics of sampling, and to Ms. Akiko Kamimura, Ms. Ryoko Kawashima, and Ms. Kayo Sakiyama for their analytical technical support. Special thanks are extended to Mr. Takuya Arai (National Institute for Environmental Studies) for his valuable contribution in integrating the sampling and oceanographic datasets used in this study. This work was supported by the Japan Agency for Medical Research and Development, Grant/Award Number 22gm1710004h0001; Japan Science and Technology Agency (JST) Grant Number JPMJCR23J4 (to K.S., Shinichiro N., and Shinji N.), Japan; JSPS Grants-in-Aid for Scientific Research (23H03516); and JST PRESTO (JPMJPR23G7).

## Footnotes

∗ As clearly shown in Figure 1, the KURO region was only sampled in its eastern half; NPSW mostly in its northern half; and PNEC in its eastern half.

† after DADA2 filtering

‡ Exponential Shannon Diversity Index (eSDI) was used as a proxy for alpha diversity

§ part of the [SST, DIC] cluster

