## Supplementary for "Patterns and Drivers of Diatom Diversity and Biogeography in the North Pacific"

#### S1 Imputation of missing environmental values

Some of our samples contain missing environmental values – either from actual lack of said measurement, or if the measurement is below the detection threshold of our instrumentation. Many statistical methods don't handle those well. We therefore inputed the missing values using a machine-learning model.

The model (AutoGluon 1.4's MultilabelPredictor) was trained to infer out original dataset's environmental values from Copernicus' Global Ocean Ensemble Physics Reanalysis and Global Ocean Biogeochemistry Hindcast datasets (variables `thetao_mean,so_mean` from `mems_mod_glo_phy-mnstd_my_0.25deg_P1D-m`; `spco2,fe,ph,phyc` from `cmems_mod_glo_bgc_my_0.25deg_P1M-m`; `chl,no3,nppv,o2,po4,si` from `cmems_mod_glo_bgc_my_0.25deg_P1D-m`).

Calibration curves are plotted in fig. 1.

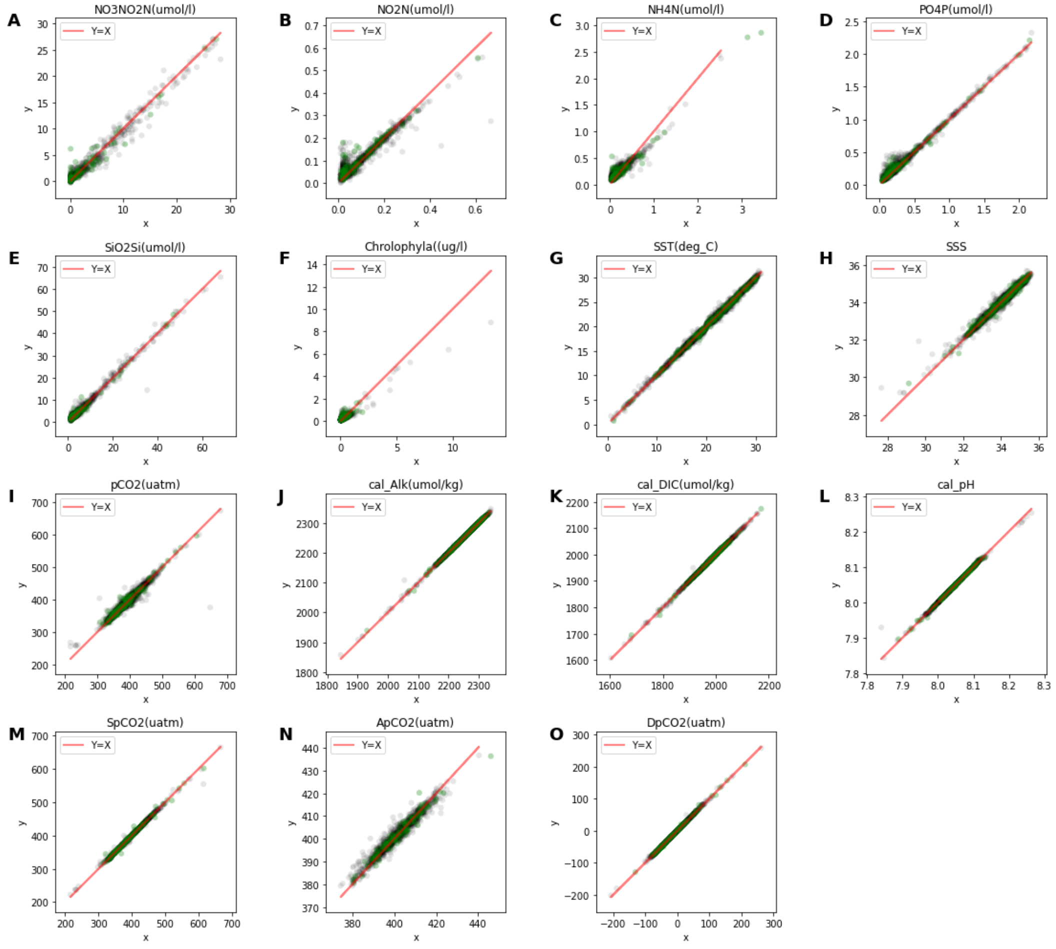


Figure 1: True ($x$) vs predicted ($y$) environmental variables. Grey (resp. green) points correspond to the training (resp. validation) set.

### S2 Dataset generation workflow from .fastq files

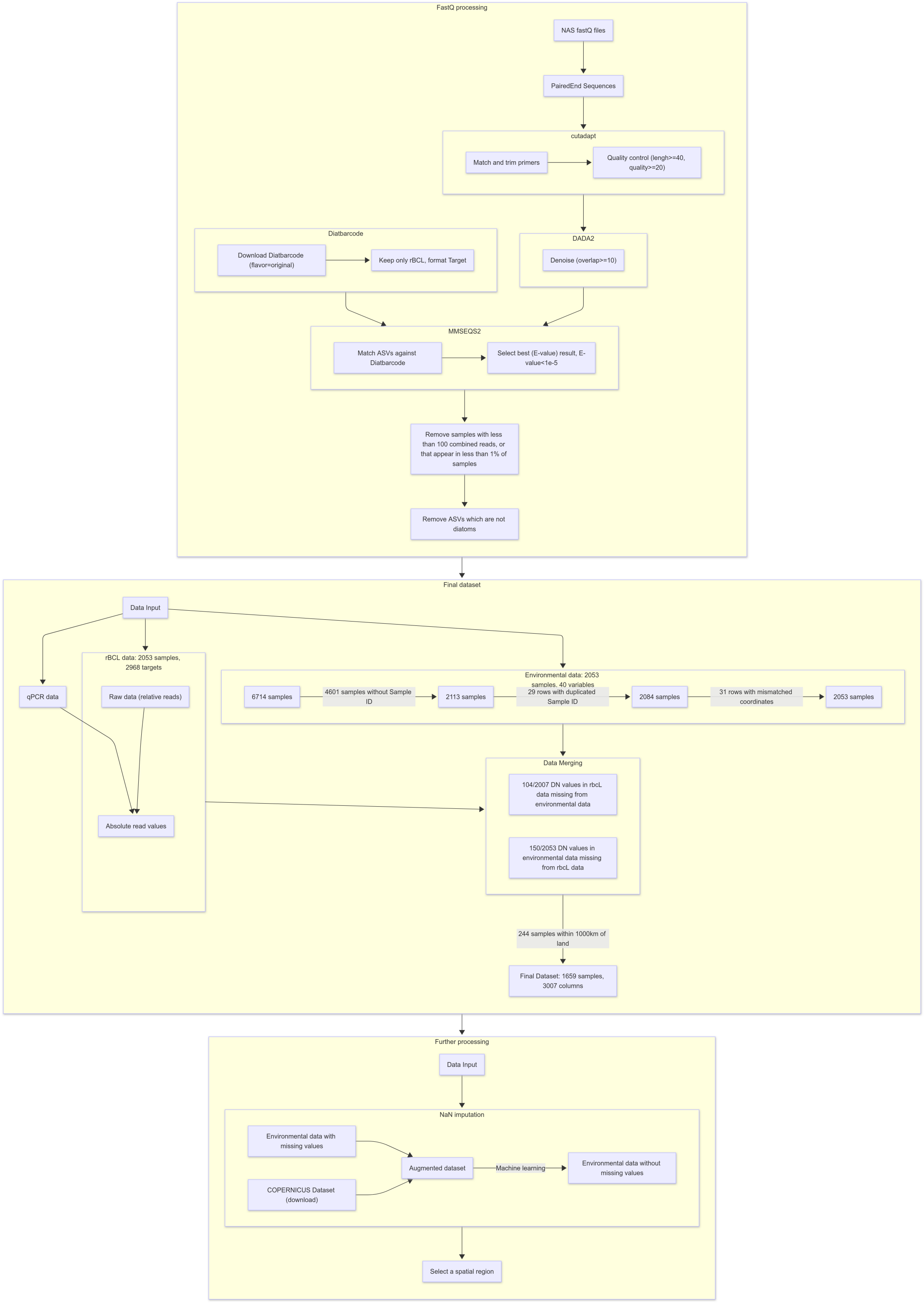


Figure 2: Dataset generation workflow: (a) from raw samples to .fastq / raw environmental values (b) Further bioinformatic processing and dataset curation

#### S3 Samples location hexmap

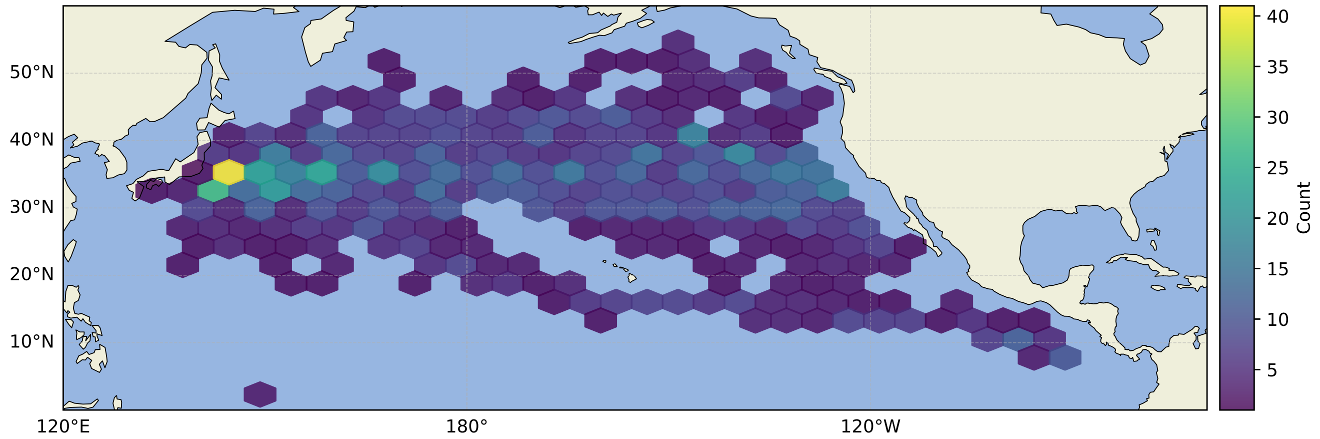


Figure 3: Number of sample per location

#### S4 Global taxonomic distribution

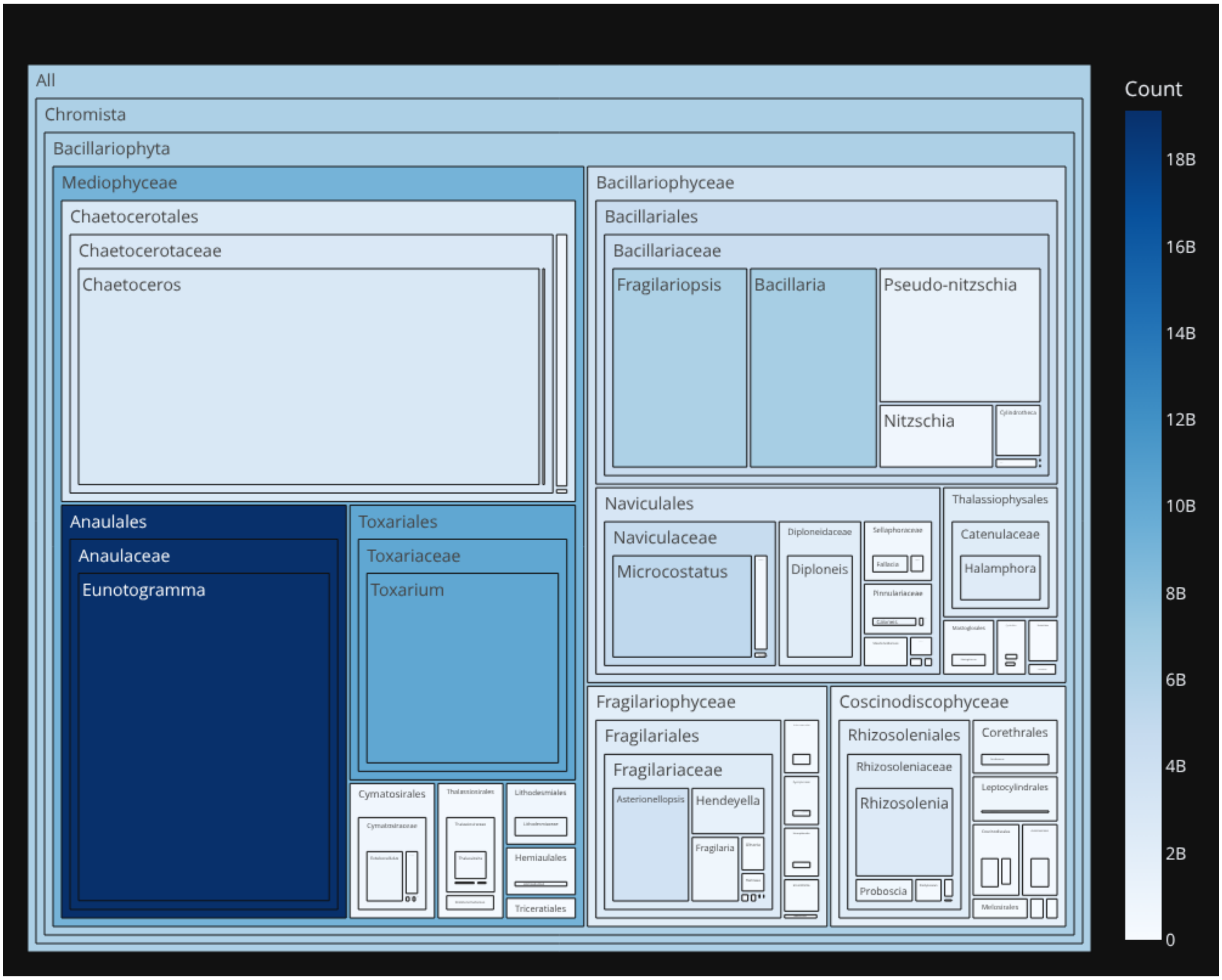


Figure 4: Treemap of the taxonomic distribution of this study's dataset. Area represents total reads; color represents average reads across species.

#### S5 Total reads and number of ASVs for each genus

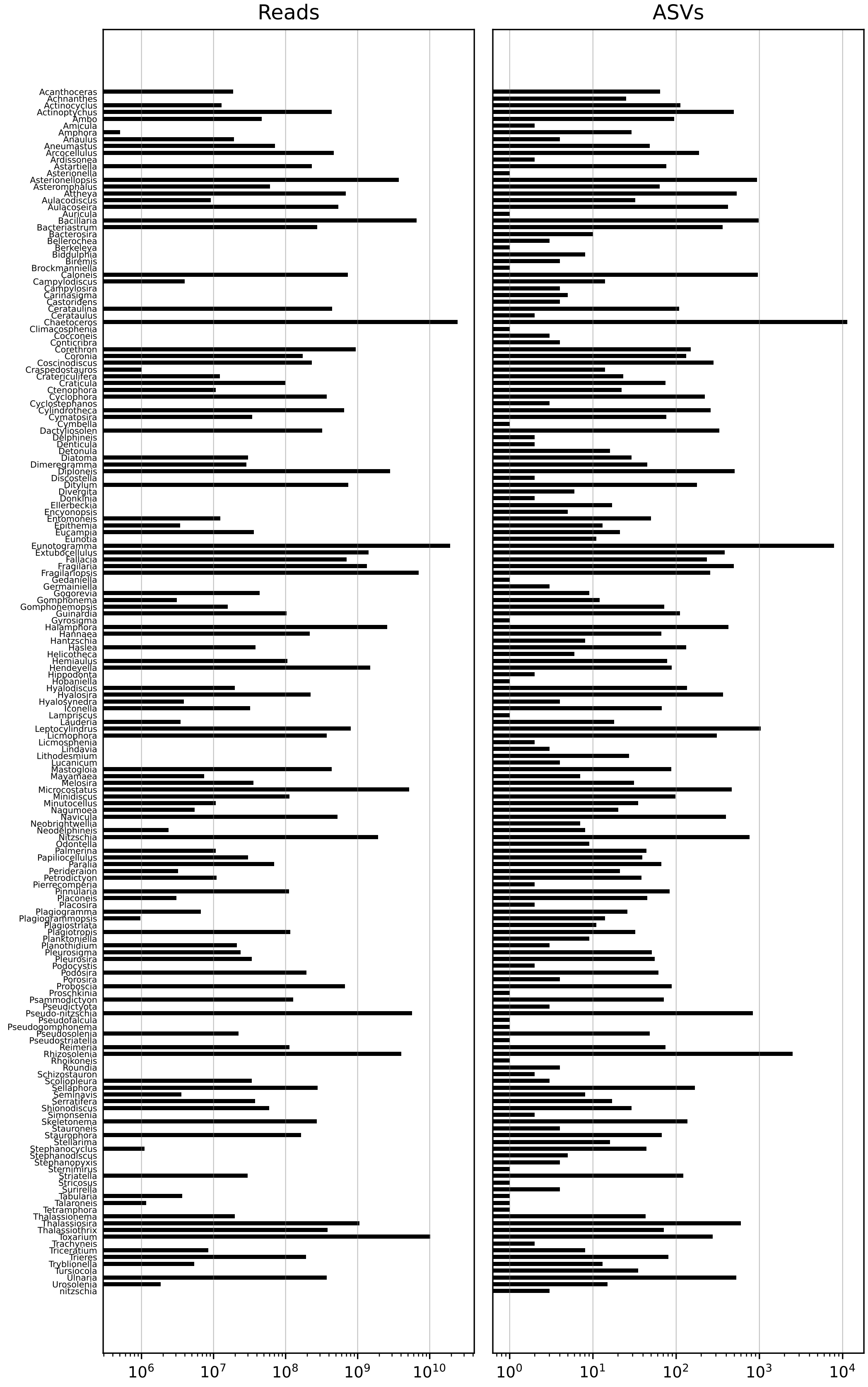


Figure 5: Comparison of read abundance and ASV richness by genus. The left panel displays the total sequencing reads, while the right panel shows the count of unique Amplicon Sequence Variants (ASVs) assigned to each genus.

#### S6 VIF clustering

The quality and interpretability of RDA decreases as the colinearity between predictors increases. To address this issue, given the significant number of predictors we consider, we clustered predictors to reduce the inter-cluster Variance Inflation Factor (VIF) among the retained variables.

More precisely, we first cluster the predictors using complete linkage on the rescaled[1] raw data. We then iteratively evaluate an increasing number of clusters. For each iteration, we identify the medoid of each cluster to serve as the cluster representative. The iterative process stops when the maximum VIF among these cluster medoids exceeds 5, at which point we retain the cluster configuration from the preceding valid step.

For all further analyses (e.g., RDA), the medoid of each cluster is used to represent the entire cluster.

VIF clusters are presented per-region in section S7.3.

[1] Highly skewed variables are first log-transformed to approximate a Gaussian distribution, then all variables are standardized to a mean of 0 and a standard deviation of 1.

### S7 Per-region plots

#### S7.1 Sample summary

Each page contains:

- the location of each sample,
- a heatmap of the number of sample for each month of each year,
- the average number of samples per month-of-year,
- the distribution of total abundance and exponential Shannon Diversity Index (eSDI) per sample,
- the evolution of average abundance and eSDI over time,
- the average abundance and eSDI per month-of-year,
- the top 10 most abundant species,
- the location and category of samples occuring during a marine heatwave,
- SST vs time.

### Region Summary Card: All

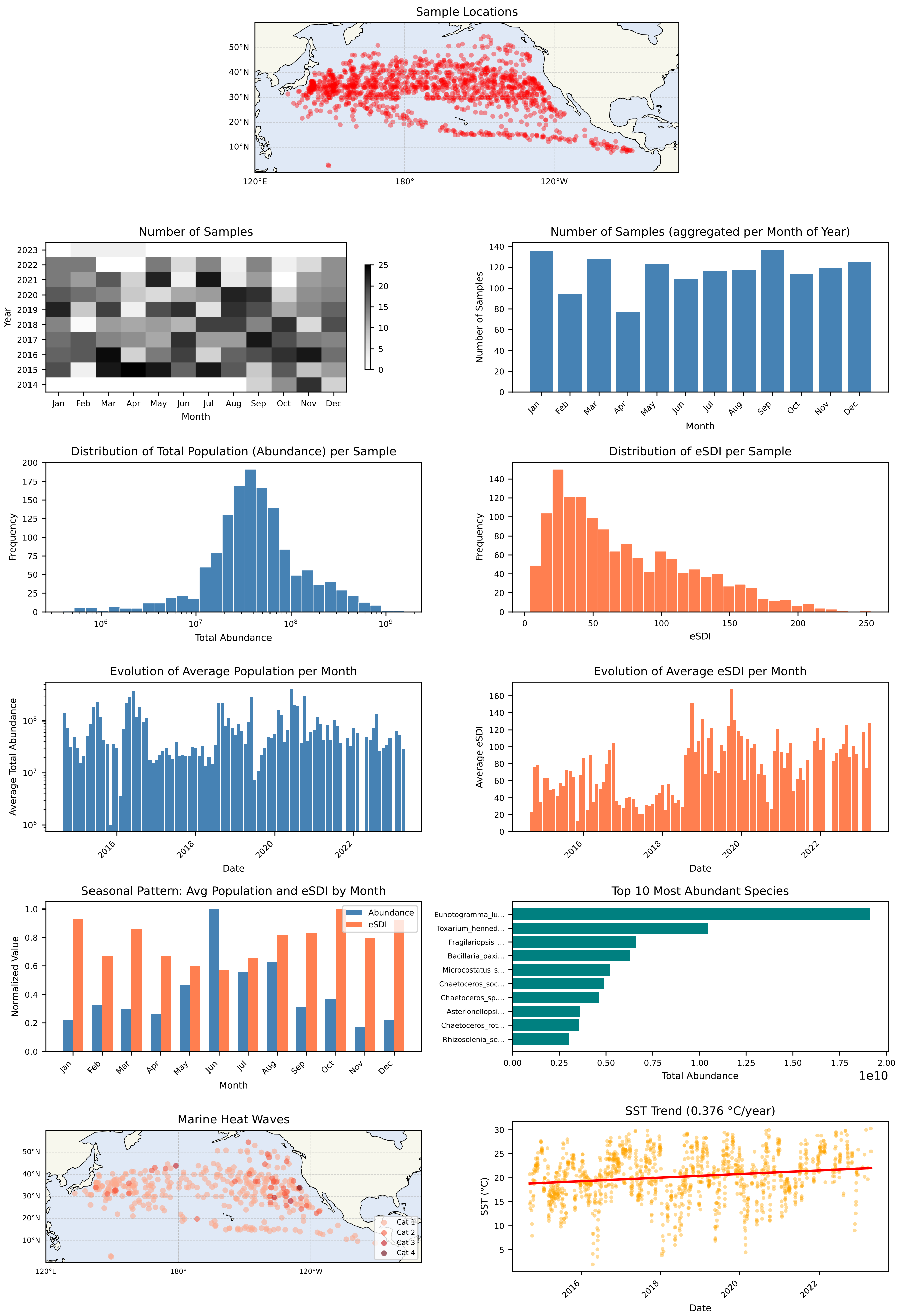

### Region Summary Card: PSAE

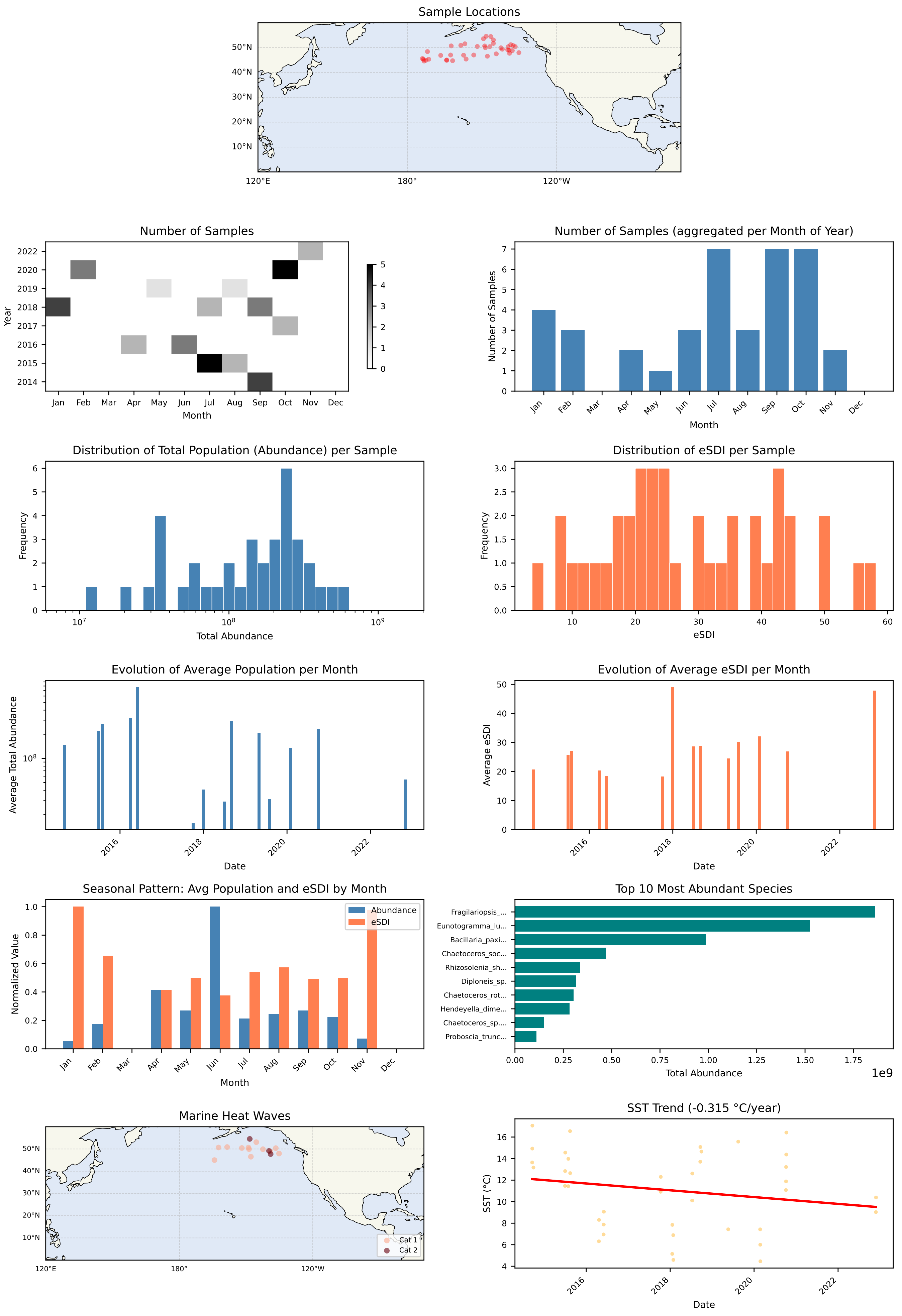

### Region Summary Card: PSAW

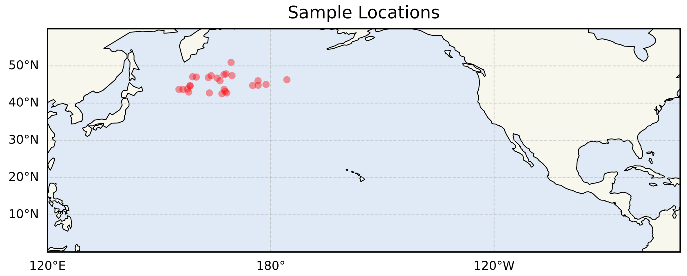


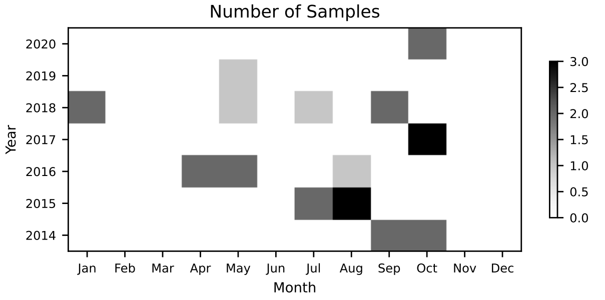


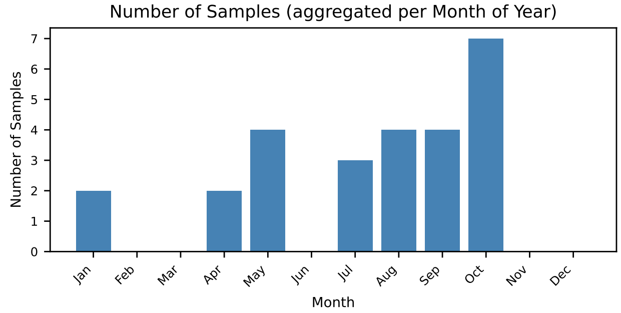


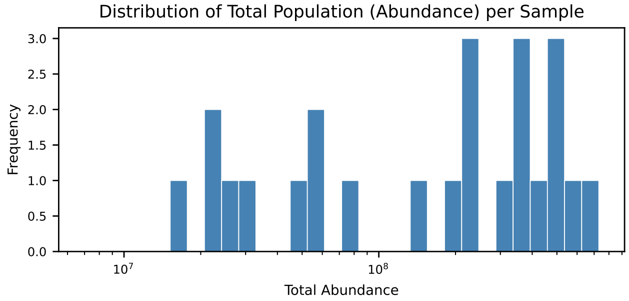


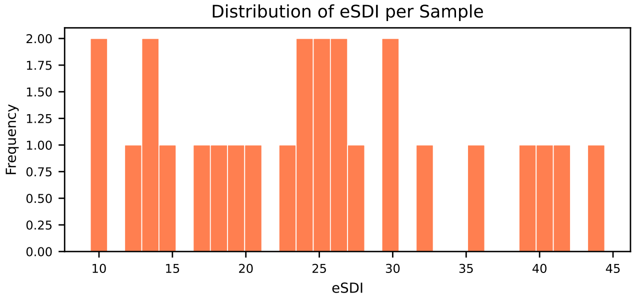


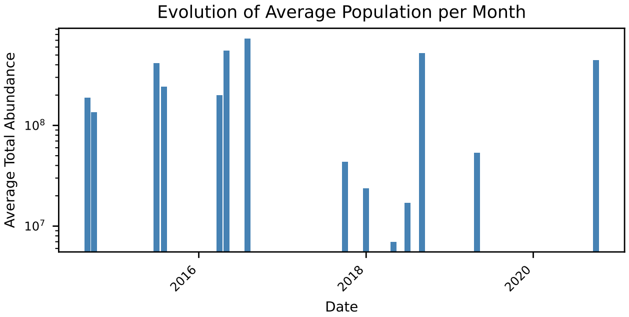


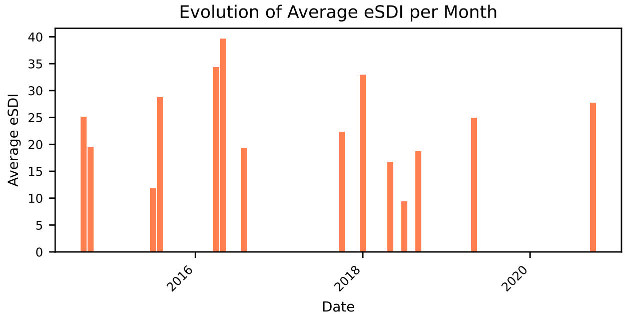


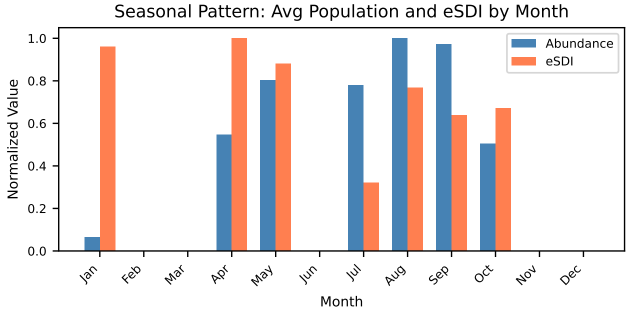


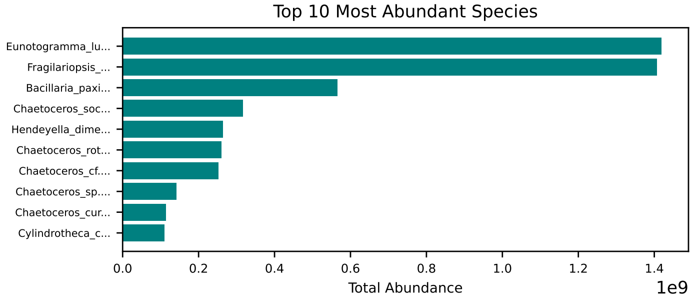


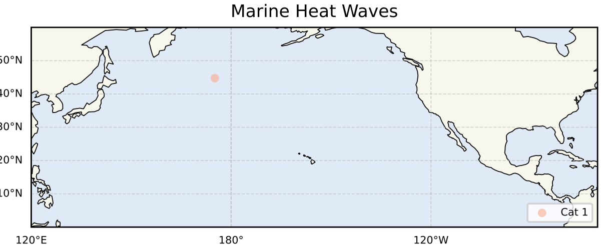


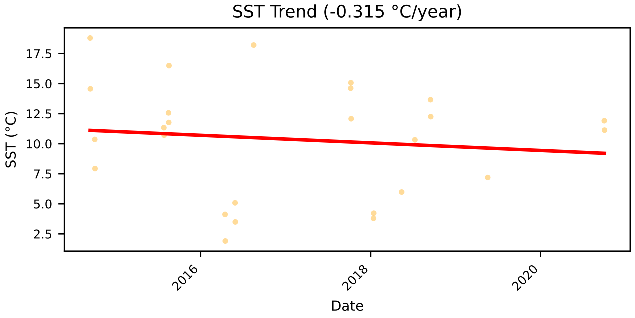

### Region Summary Card: KURO

Sample Locations

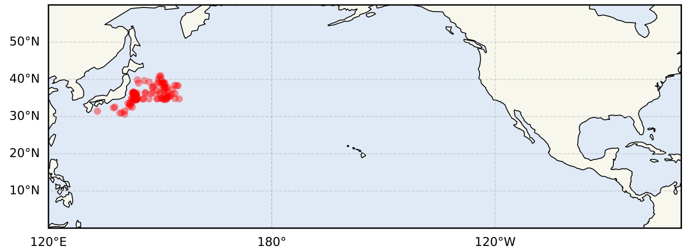


Number of Samples

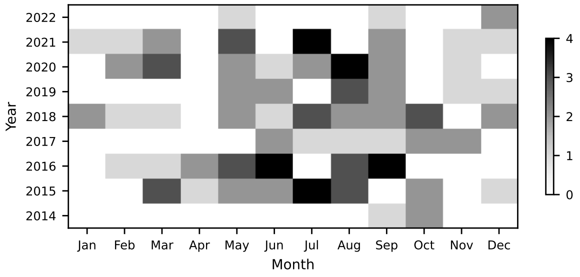


Number of Samples (aggregated per Month of Year)

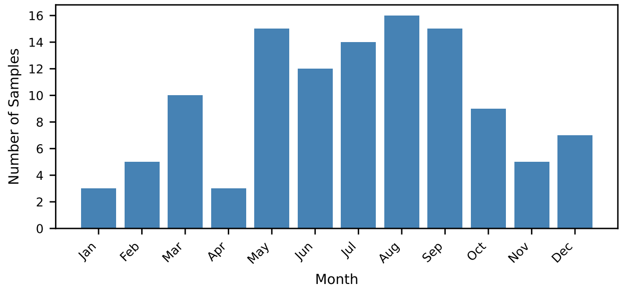


Distribution of Total Population (Abundance) per Sample

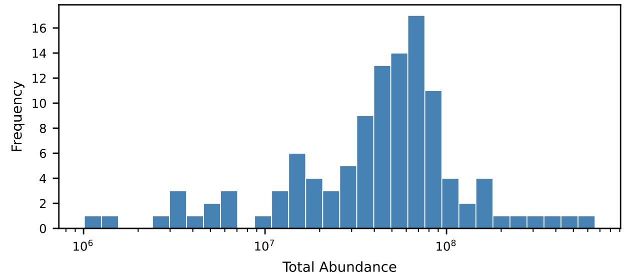


Distribution of eSDI per Sample

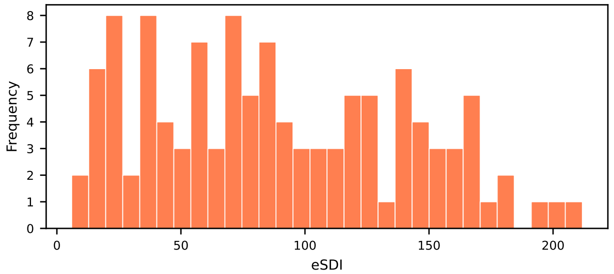


Evolution of Average Population per Month

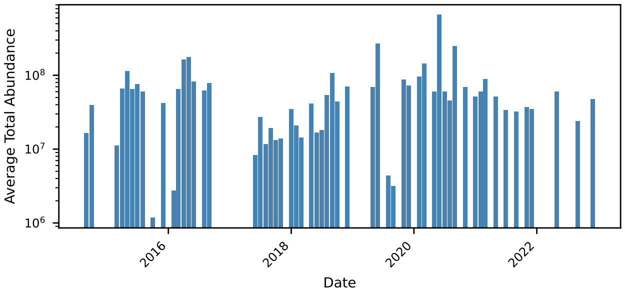


Evolution of Average eSDI per Month

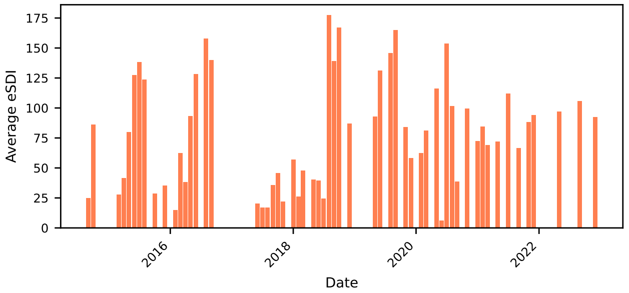


Seasonal Pattern: Avg Population and eSDI by Month

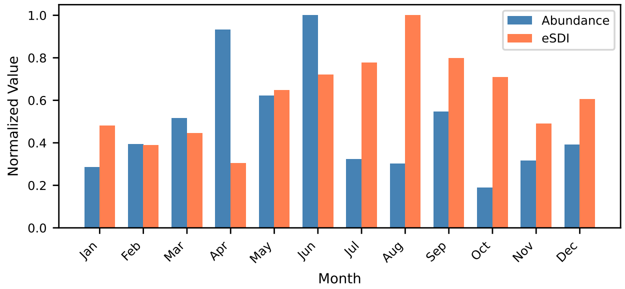


Top 10 Most Abundant Species

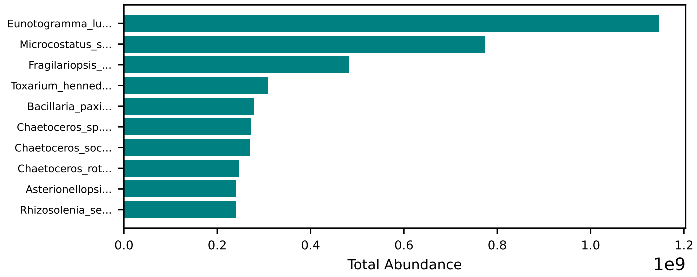


Marine Heat Waves

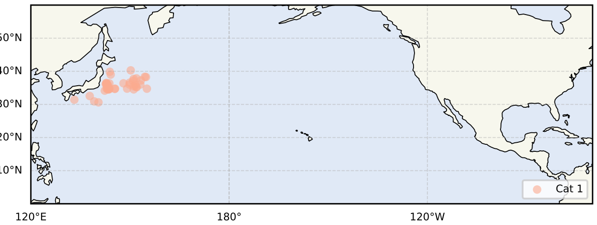


SST Trend (0.133 °C/year)

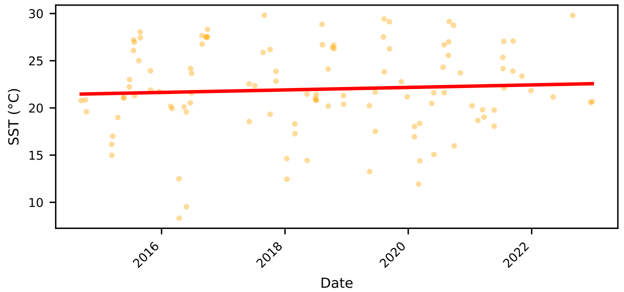

### Region Summary Card: NPPF

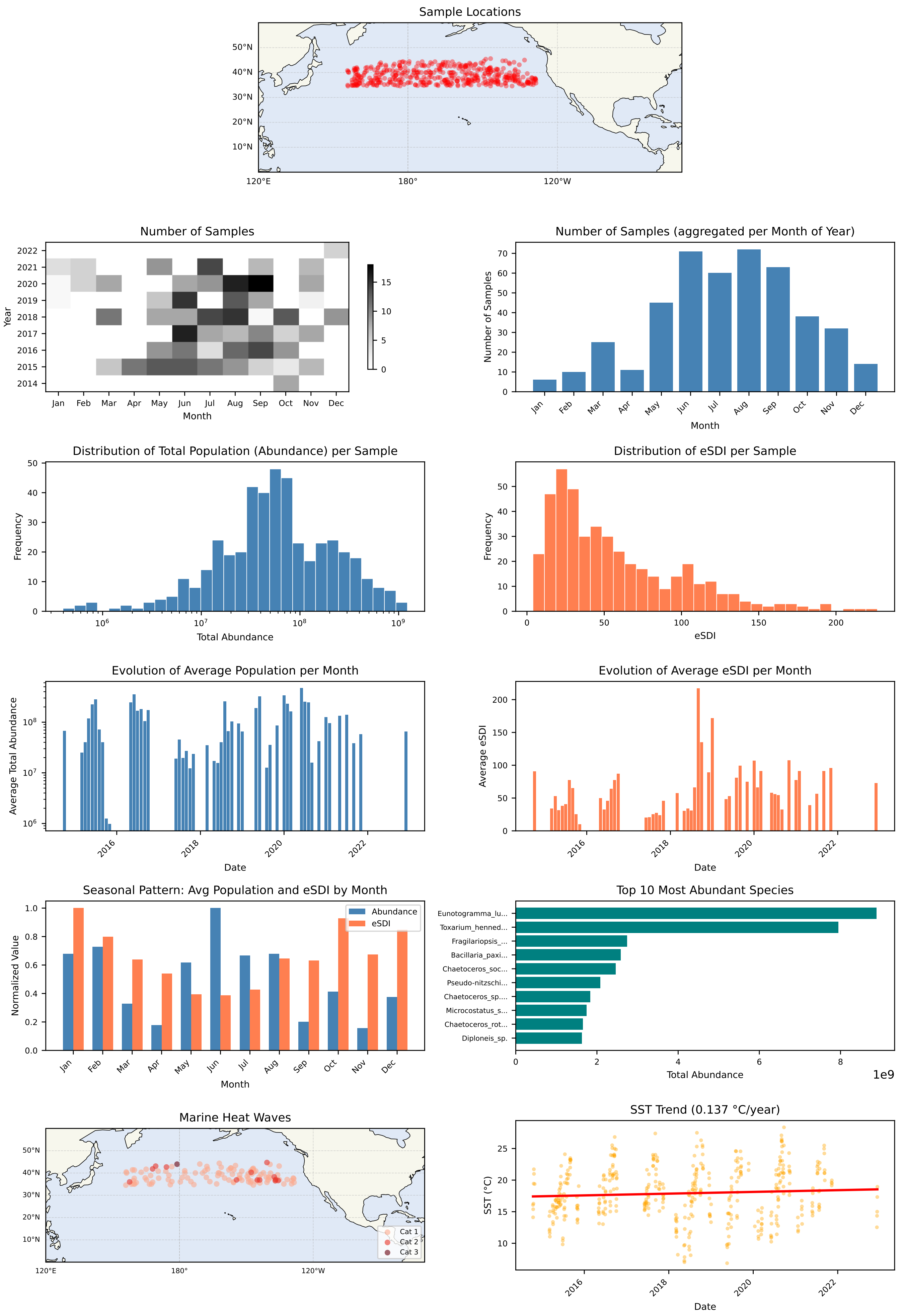

### Region Summary Card: NPSW

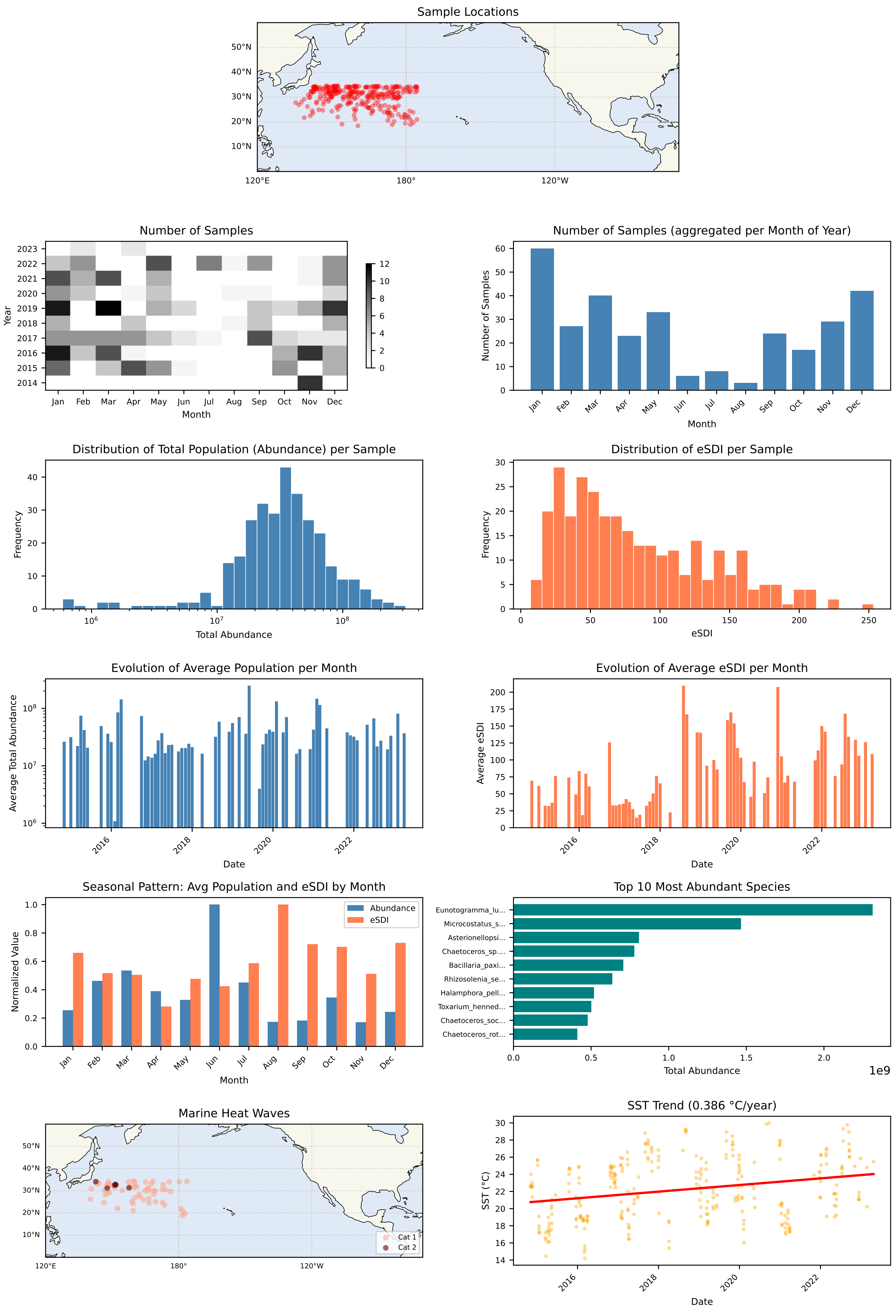

### Region Summary Card: NPTG

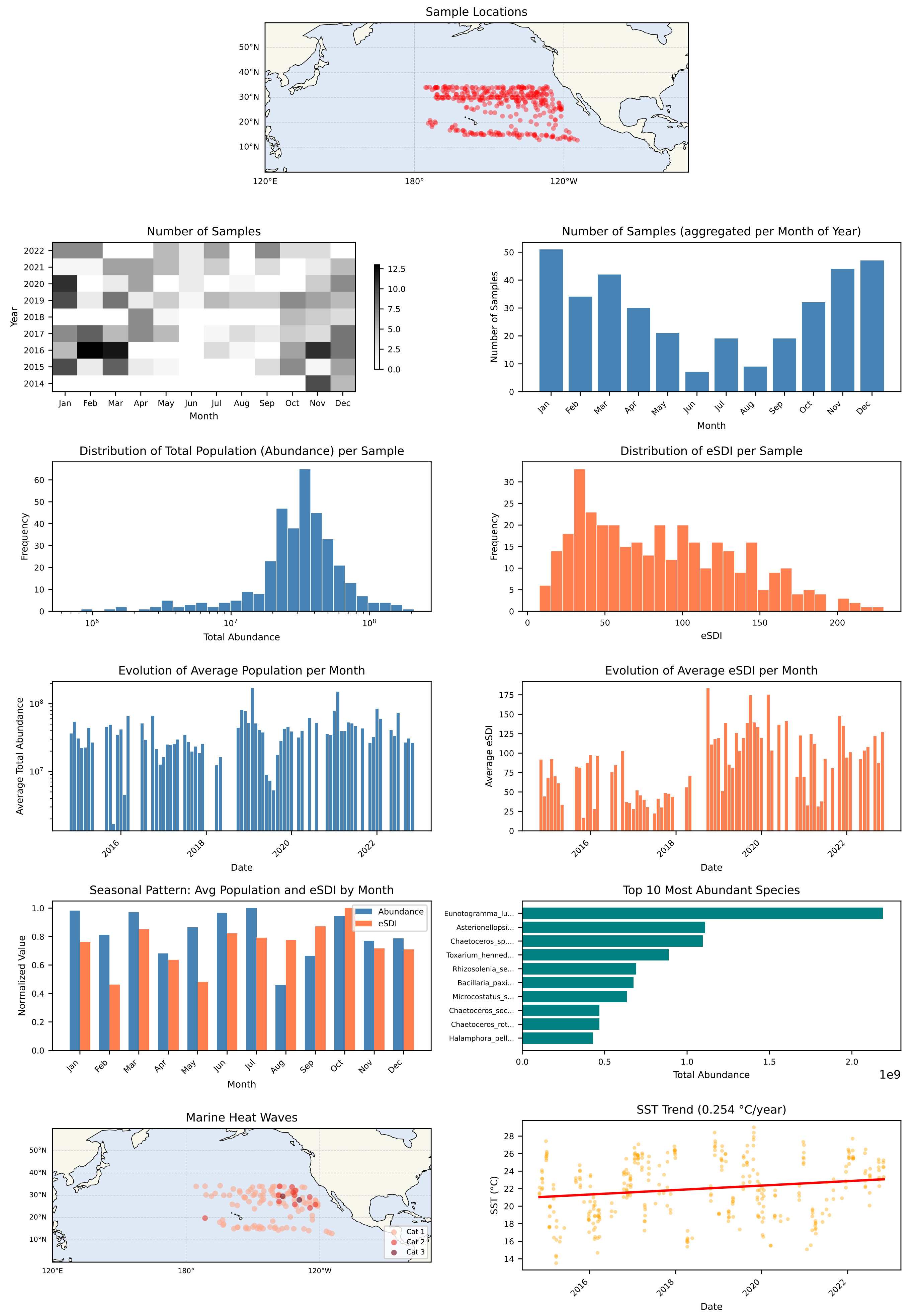

### Region Summary Card: CCAL

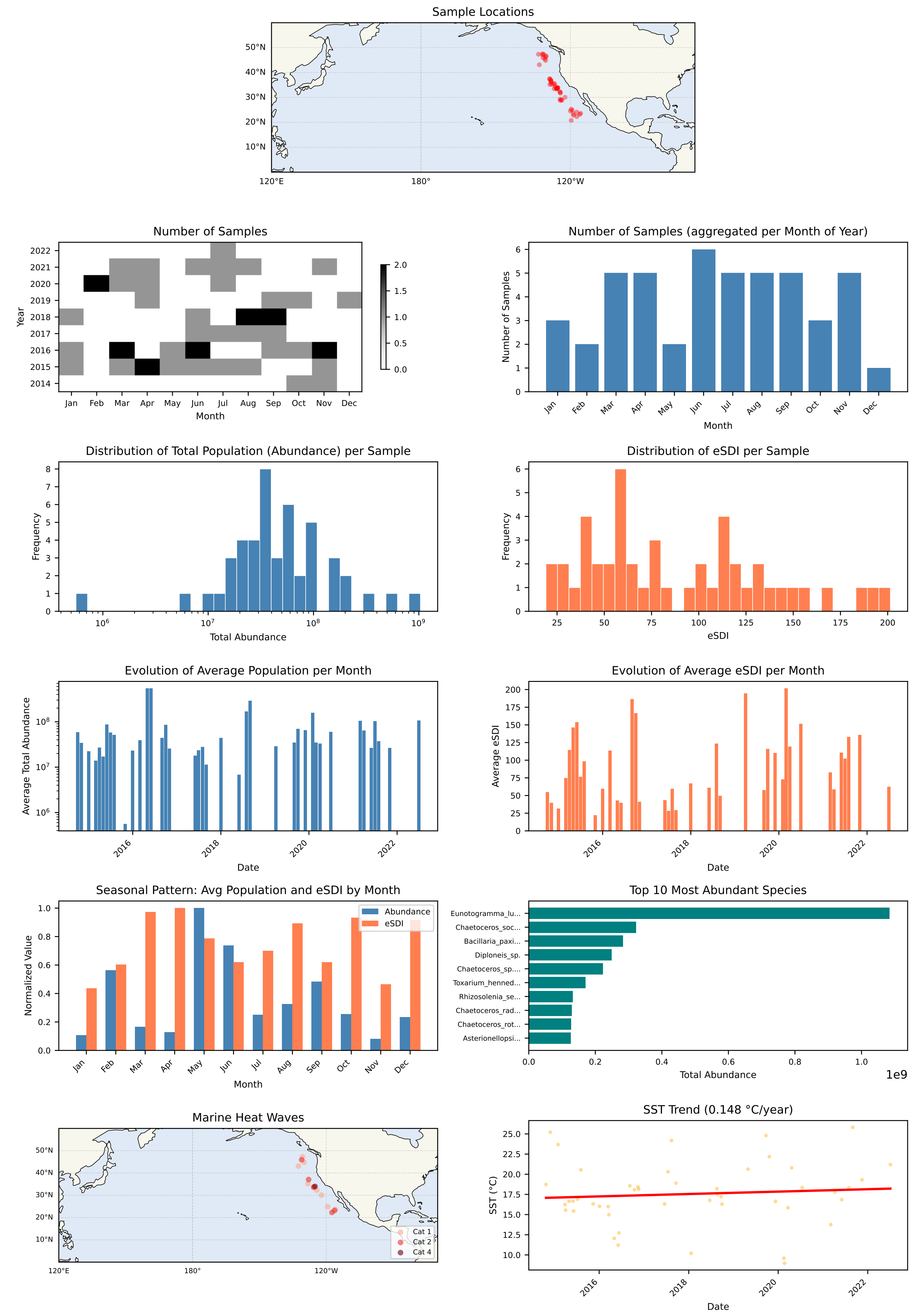

### Region Summary Card: PNEC

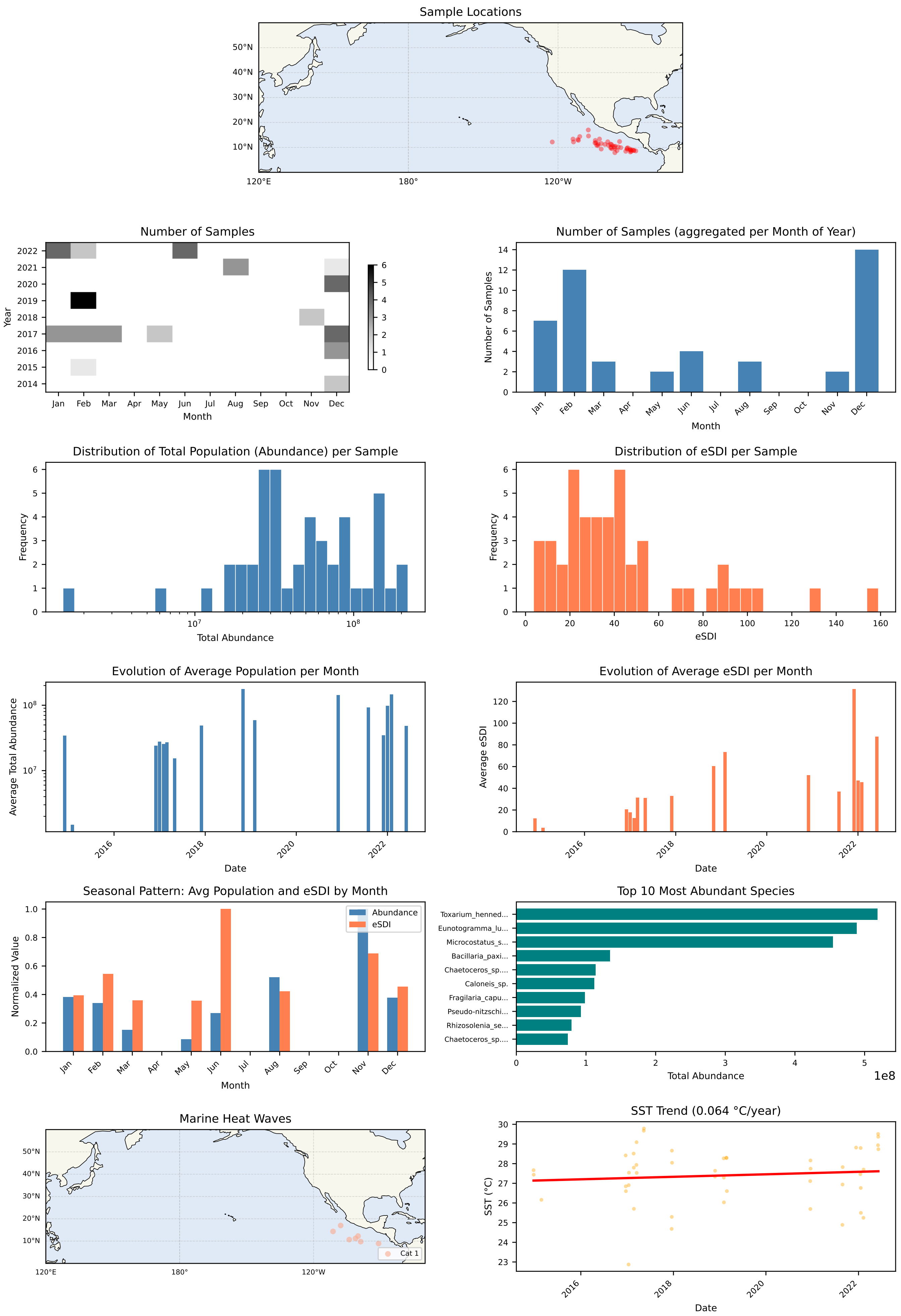

#### S7.2 Taxonomic composition

Each double-page contains:

- normalized sample composition at the Order (resp. Family / Genus / Species) level,
- for the top 9 species, normalized abundance proportion per month-of-year,
- distribution of eSDI for each month-of-year,
- NMDS ordination of sample populations, color-coded by fractional month-of-year,
- species differentially abundant in a given season, per ANCOM. Season 1 is Jan-April, season 2 is May-August, etc.

**TOP N Taxonomic Composition by Month: All**

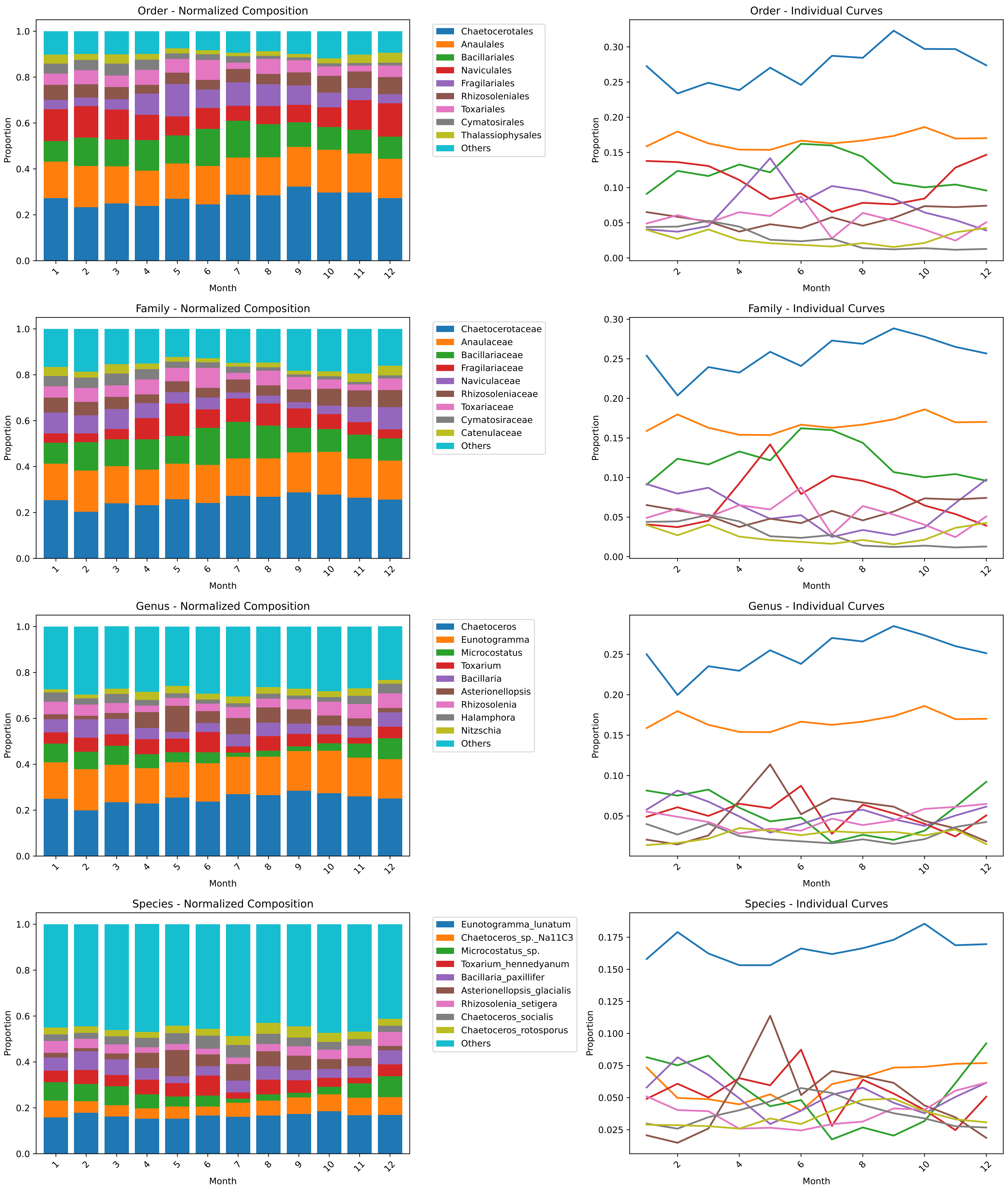

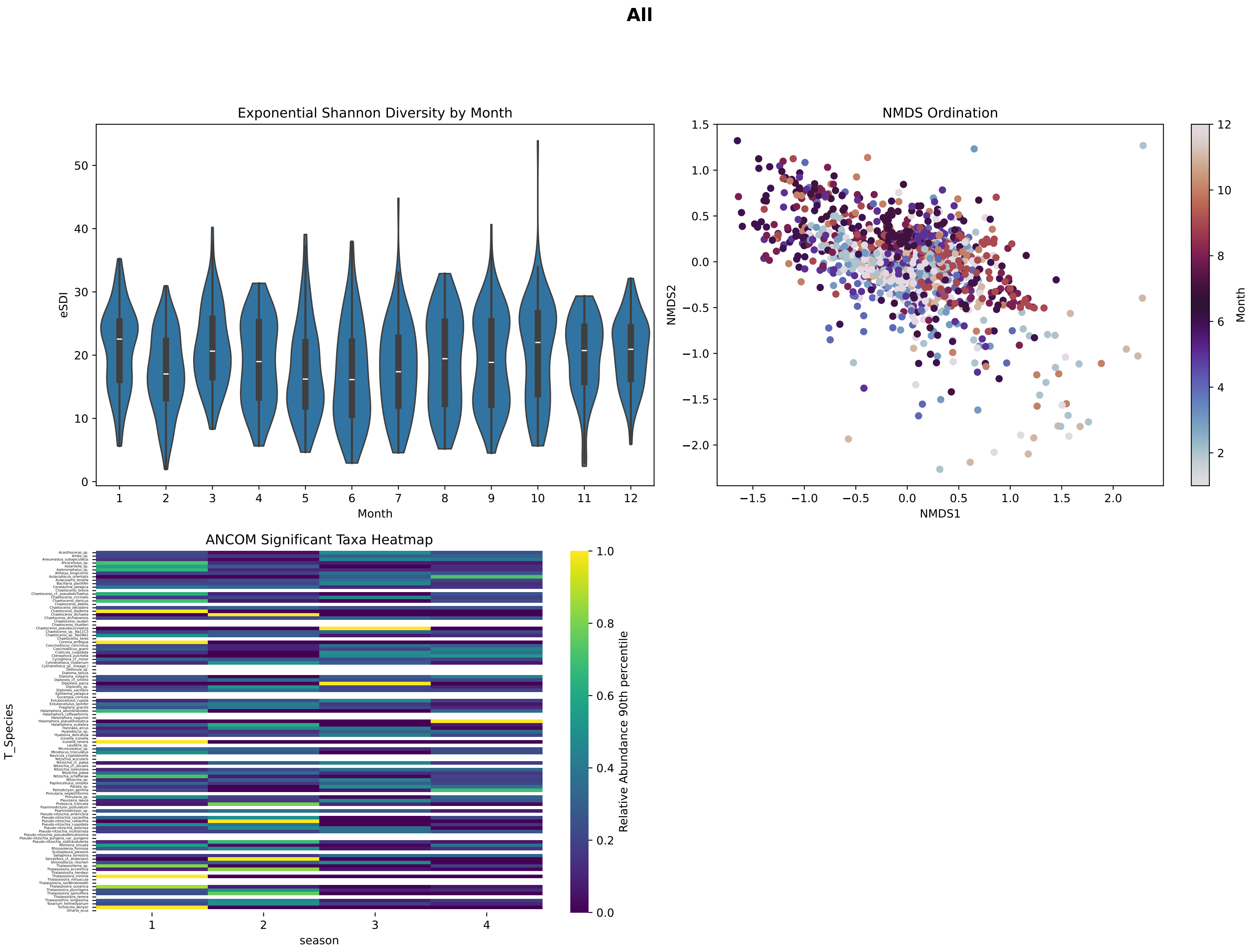
All
Exponential Shannon Diversity by Month
eSDI
Month
NMDS Ordination
NMDS1
NMDS2
Month
ANCOM Significant Taxa Heatmap
T_Species
season
Relative Abundance 90th percentile

**TOP N Taxonomic Composition by Month: PSAE**

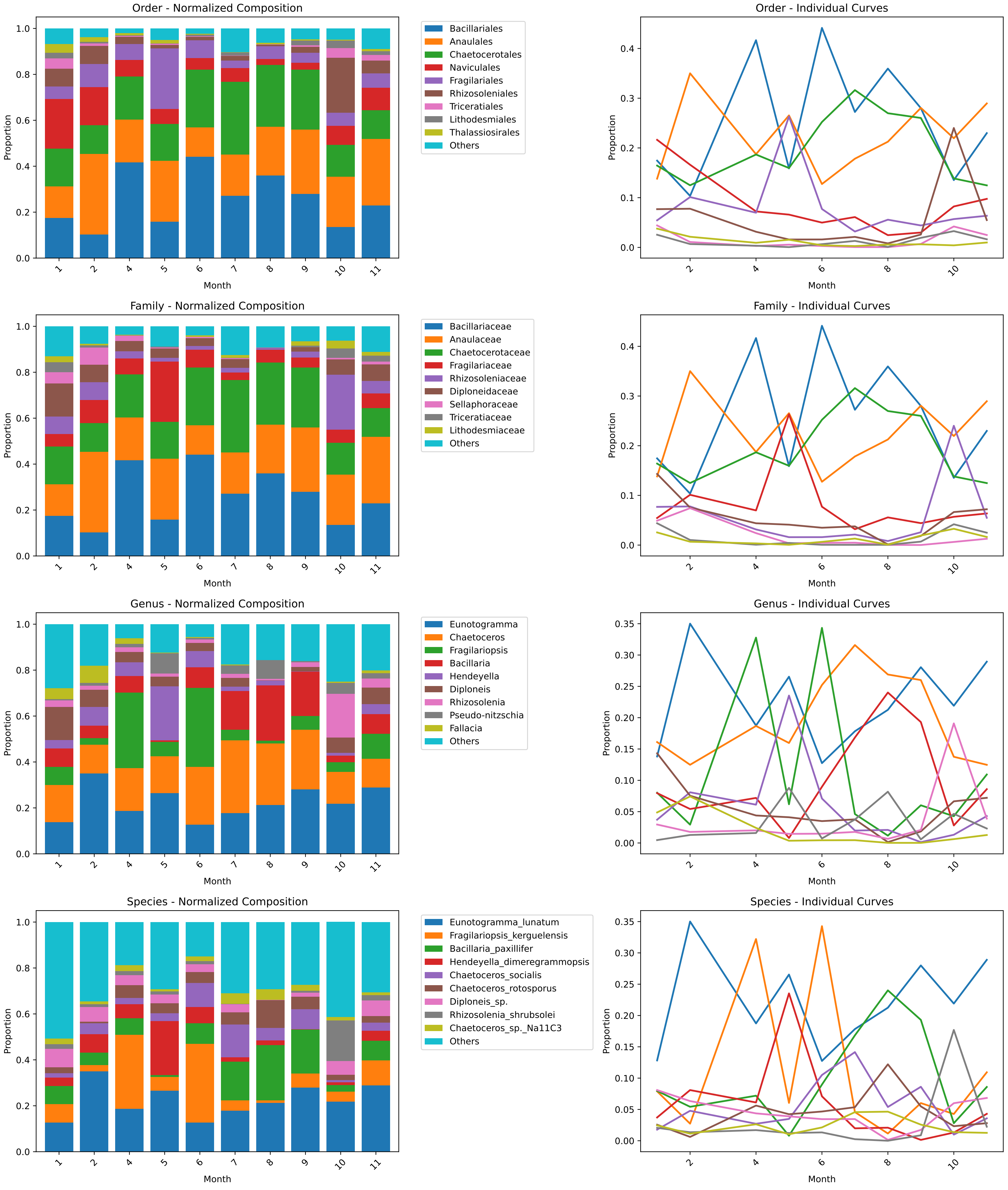

**PSAE**

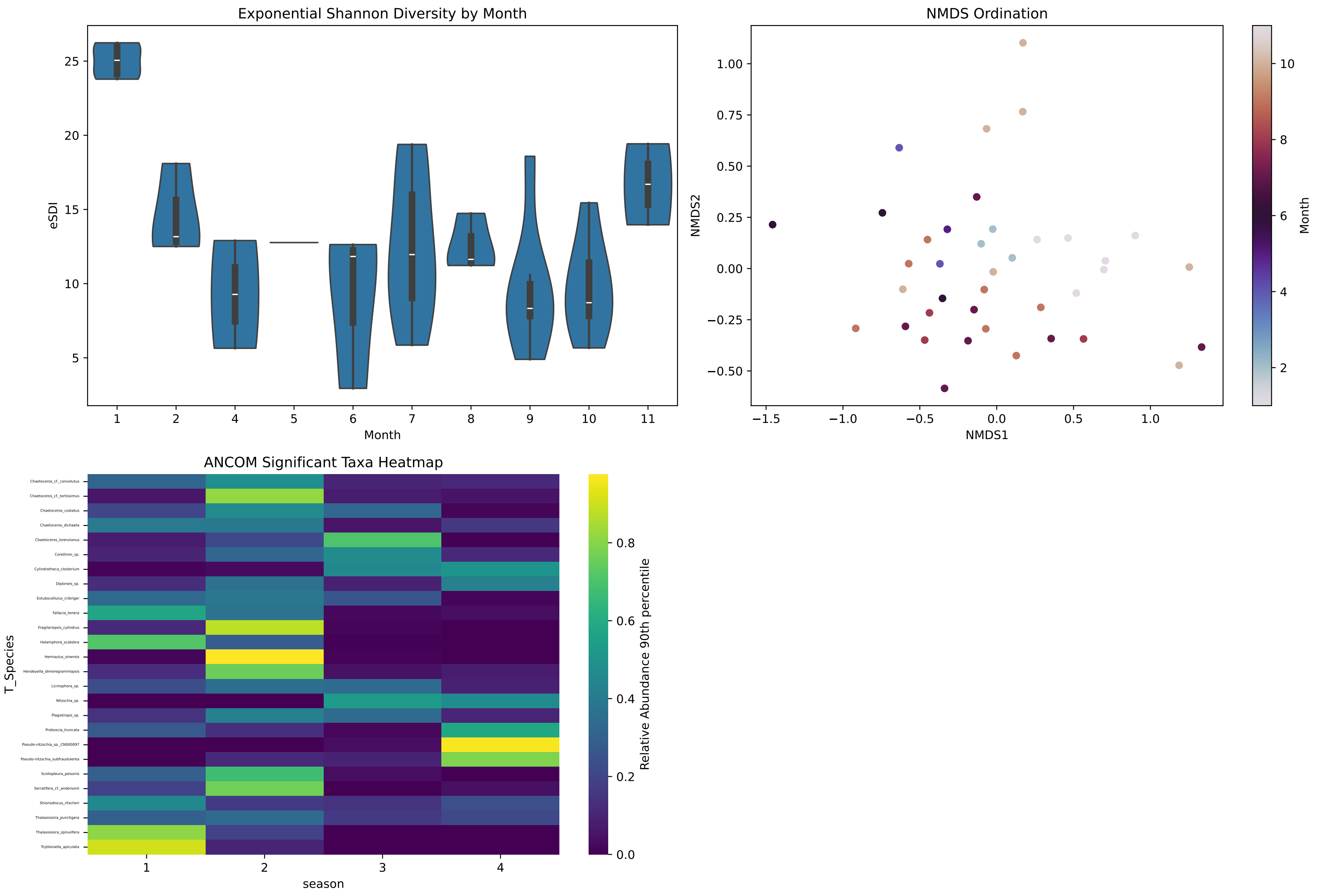

Exponential Shannon Diversity by Month
eSDI
Month
5
10
15
20
25
1
2
4
5
6
7
8
9
10
11
NMDS Ordination
NMDS2
NMDS1
1.00
0.75
0.50
0.25
0.00
−0.25
−0.50
−1.5
−1.0
−0.5
0.0
0.5
1.0
Month
2
4
6
8
10
ANCOM Significant Taxa Heatmap
T_Species
Chaetoceros_cf._convolutus
Chaetoceros_cf._tortissimus
Chaetoceros_costatus
Chaetoceros_dichaeta
Chaetoceros_lorenzianus
Corethron_sp.
Cylindrotheca_closterium
Diploneis_sp.
Extubocellulus_cribriger
Fallacia_tenera
Fragilariopsis_cylindrus
Halamphora_scatebra
Hemiaulus_sinensis
Hendeyella_dimeregrammopsis
Licmophora_sp.
Nitzschia_sp.
Plagiotropis_sp.
Proboscia_truncata
Pseudo-nitzschia_sp._CNS00097
Pseudo-nitzschia_subfraudulenta
Scoliopleura_peisonis
Serratifera_cf._andersonii
Shionodiscus_ritscheri
Thalassiosira_punctigera
Thalassiosira_spinulifera
Tryblionella_apiculata
season
1
2
3
4
Relative Abundance 90th percentile
0.0
0.2
0.4
0.6
0.8

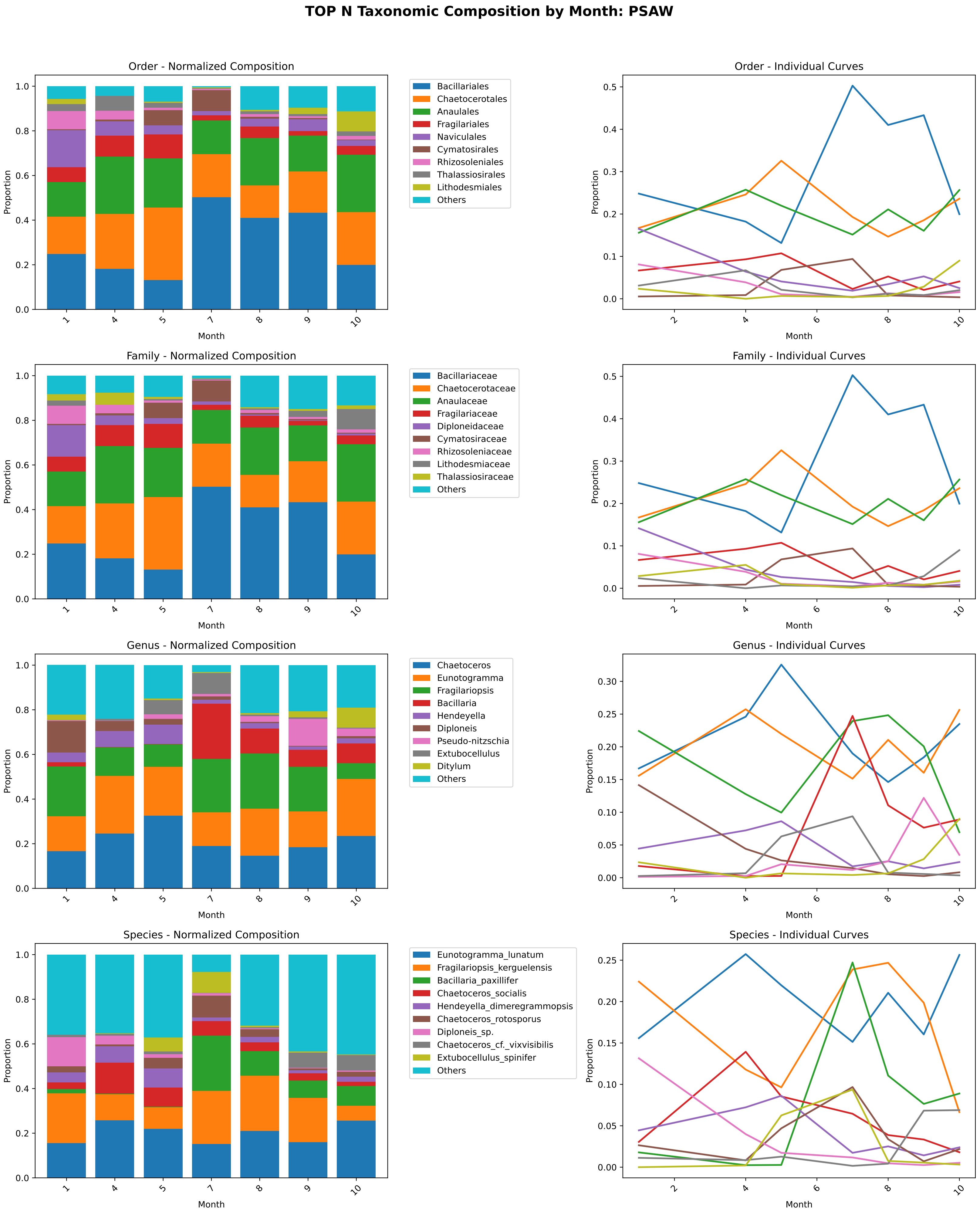
TOP N Taxonomic Composition by Month: PSAW
Order - Normalized Composition
Bacillariales
Chaetocerotales
Anaulales
Fragilariales
Naviculales
Cymatosirales
Rhizosoleniales
Thalassiosirales
Lithodesmiales
Others
Order - Individual Curves
Family - Normalized Composition
Bacillariaceae
Chaetocerotaceae
Anaulaceae
Fragilariaceae
Diploneidaceae
Cymatosiraceae
Rhizosoleniaceae
Lithodesmiaceae
Thalassiosiraceae
Others
Family - Individual Curves
Genus - Normalized Composition
Chaetoceros
Eunotogramma
Fragilariopsis
Bacillaria
Hendeyella
Diploneis
Pseudo-nitzschia
Extubocellulus
Ditylum
Others
Genus - Individual Curves
Species - Normalized Composition
Eunotogramma_lunatum
Fragilariopsis_kerguelensis
Bacillaria_paxillifer
Chaetoceros_socialis
Hendeyella_dimeregrammopsis
Chaetoceros_rotosporus
Diploneis_sp.
Chaetoceros_cf._vixvisibilis
Extubocellulus_spinifer
Others
Species - Individual Curves
Proportion
Month

### PSAW

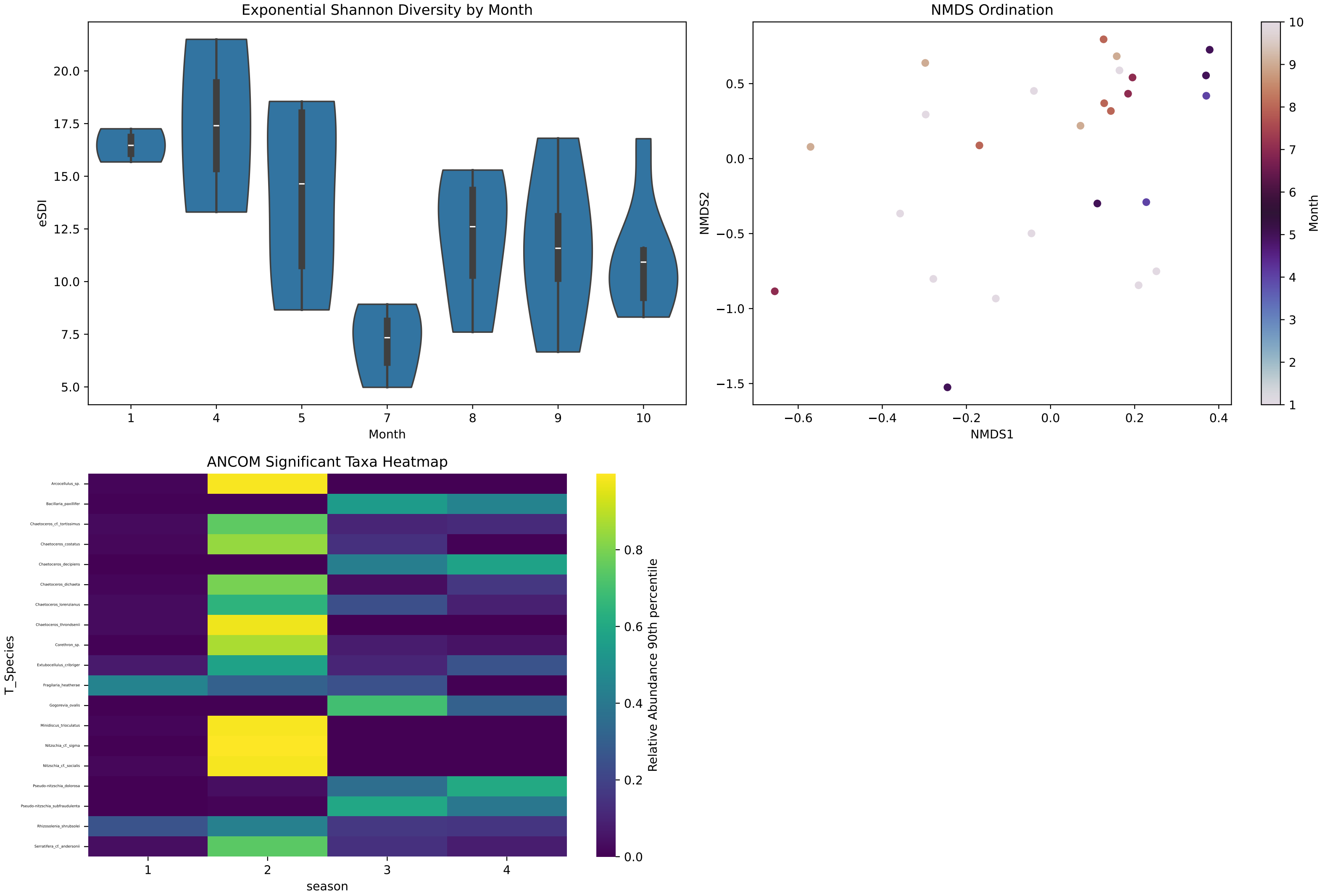

Exponential Shannon Diversity by Month
eSDI
Month
1
4
5
7
8
9
10
5.0
7.5
10.0
12.5
15.0
17.5
20.0
NMDS Ordination
NMDS1
NMDS2
−0.6
−0.4
−0.2
0.0
0.2
0.4
−1.5
−1.0
−0.5
0.0
0.5
Month
1
2
3
4
5
6
7
8
9
10
ANCOM Significant Taxa Heatmap
T_Species
season
Arcocellulus_sp.
Bacillaria_paxillifer
Chaetoceros_cf._tortissimus
Chaetoceros_costatus
Chaetoceros_decipiens
Chaetoceros_dichaeta
Chaetoceros_lorenzianus
Chaetoceros_throndsenii
Corethron_sp.
Extubocellulus_cribriger
Fragilaria_heatherae
Gogorevia_ovalis
Minidiscus_trioculatus
Nitzschia_cf._sigma
Nitzschia_cf._socialis
Pseudo-nitzschia_dolorosa
Pseudo-nitzschia_subfraudulenta
Rhizosolenia_shrubsolei
Serratifera_cf._andersonii
1
2
3
4
Relative Abundance 90th percentile
0.0
0.2
0.4
0.6
0.8

### TOP N Taxonomic Composition by Month: KURO

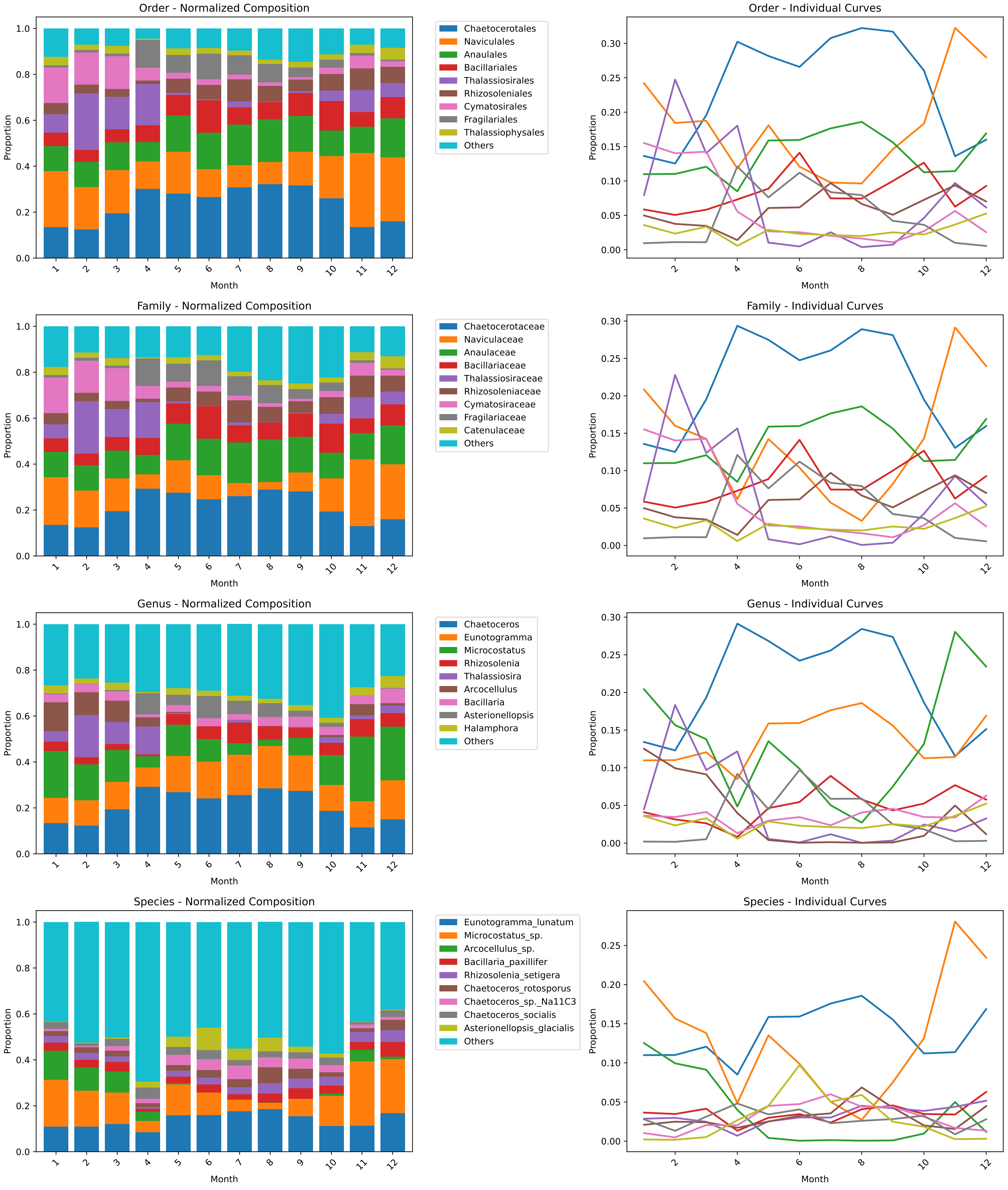

**KURO**

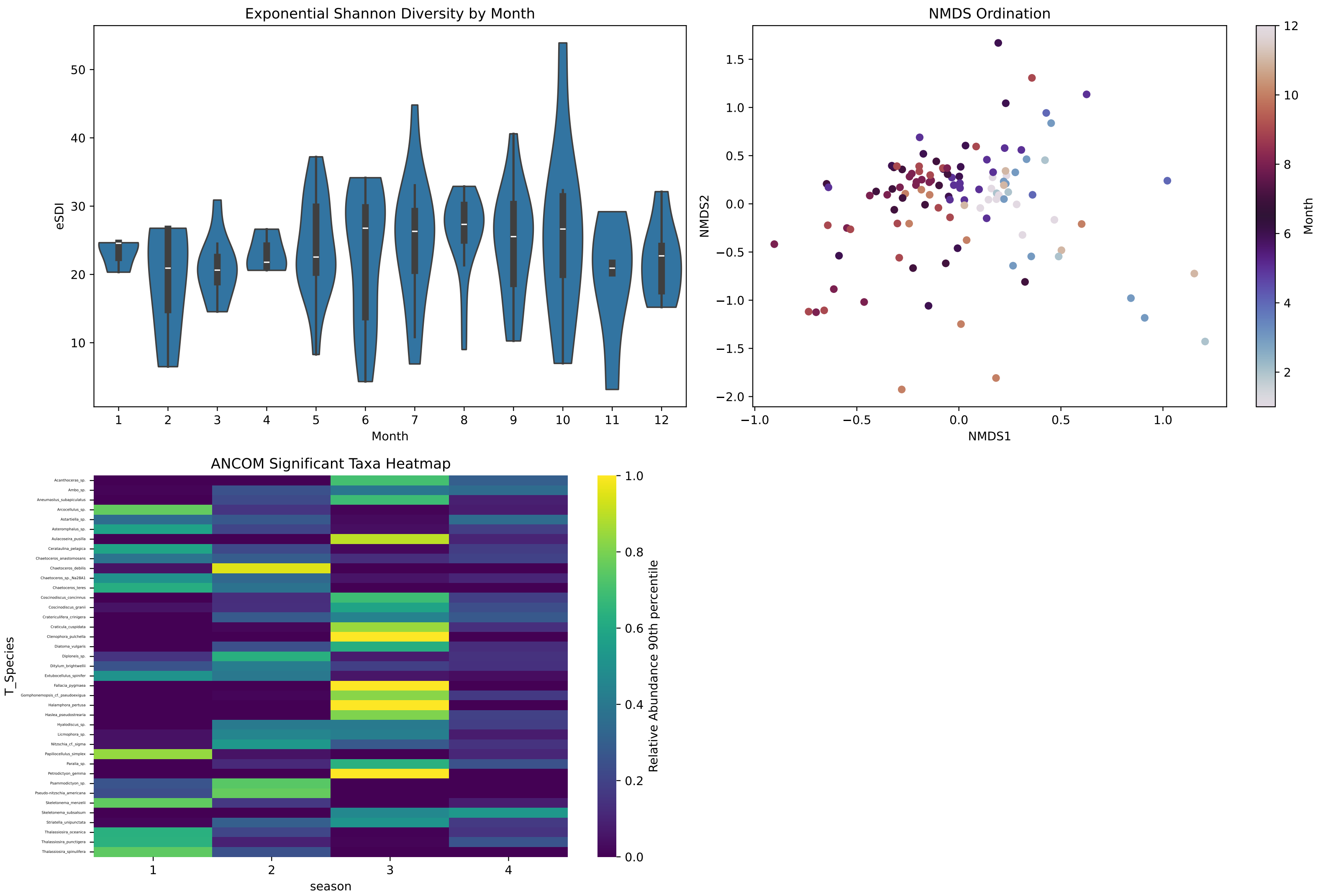
Exponential Shannon Diversity by Month
eSDI
10
20
30
40
50
1
2
3
4
5
6
7
8
9
10
11
12
Month
NMDS Ordination
NMDS2
NMDS1
1.5
1.0
0.5
0.0
−0.5
−1.0
−1.5
−2.0
−1.0
−0.5
0.0
0.5
1.0
Month
2
4
6
8
10
12
ANCOM Significant Taxa Heatmap
T_Species
Acanthoceras_sp.
Ambo_sp.
Aneumastus_subapiculatus
Arcocellulus_sp.
Astartiella_sp.
Asteromphalus_sp.
Aulacoseira_pusilla
Cerataulina_pelagica
Chaetoceros_anastomosans
Chaetoceros_debilis
Chaetoceros_sp._Na28A1
Chaetoceros_teres
Coscinodiscus_concinnus
Coscinodiscus_granii
Cratericulifera_crinigera
Craticula_cuspidata
Ctenophora_pulchella
Diatoma_vulgaris
Diploneis_sp.
Ditylum_brightwellii
Extubocellulus_spinifer
Fallacia_pygmaea
Gomphonemopsis_cf._pseudoexigua
Halamphora_pertusa
Haslea_pseudostrearia
Hyalodiscus_sp.
Licmophora_sp.
Nitzschia_cf._sigma
Papiliocellulus_simplex
Paralia_sp.
Petrodictyon_gemma
Psammodictyon_sp.
Pseudo-nitzschia_americana
Skeletonema_menzelii
Skeletonema_subsalsum
Striatella_unipunctata
Thalassiosira_oceanica
Thalassiosira_punctigera
Thalassiosira_spinulifera
1
2
3
4
season
Relative Abundance 90th percentile
0.0
0.2
0.4
0.6
0.8
1.0

#### TOP N Taxonomic Composition by Month: NPPF

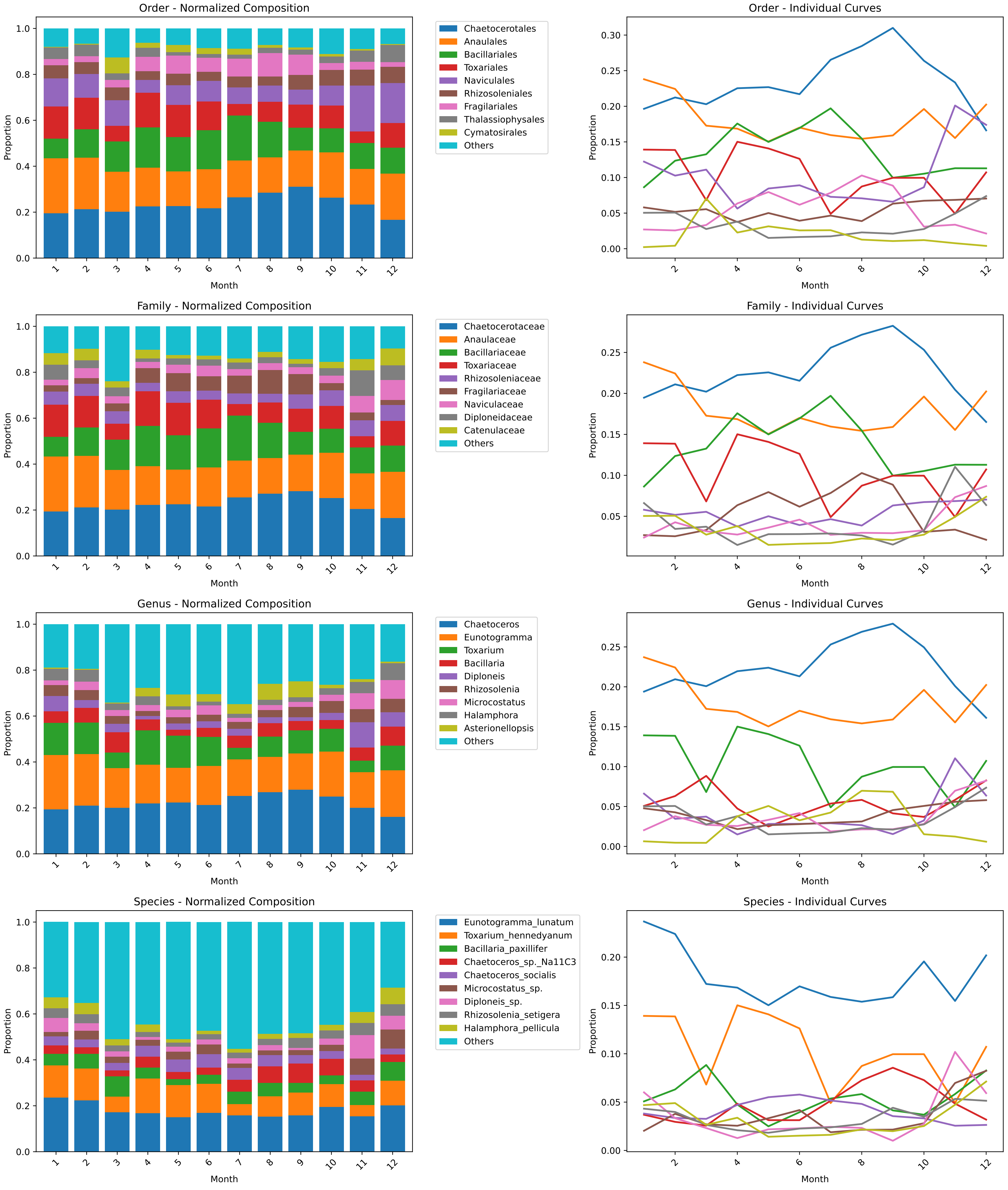

**NPPF**

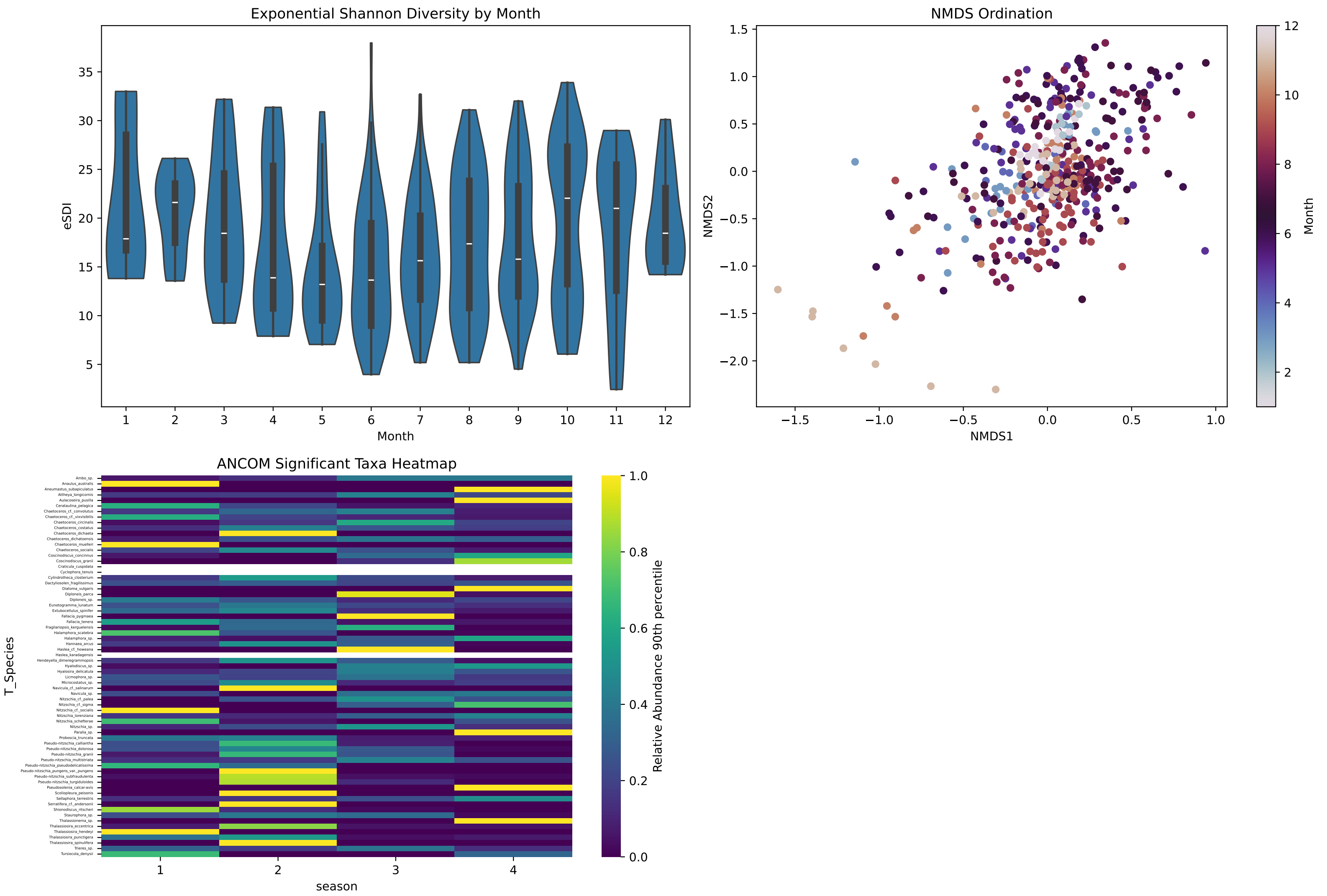
Exponential Shannon Diversity by Month
eSDI
Month
5
10
15
20
25
30
35
1
2
3
4
5
6
7
8
9
10
11
12
NMDS Ordination
NMDS2
NMDS1
−2.0
−1.5
−1.0
−0.5
0.0
0.5
1.0
1.5
−1.5
−1.0
−0.5
0.0
0.5
1.0
Month
2
4
6
8
10
12
ANCOM Significant Taxa Heatmap
T_Species
season
1
2
3
4
Relative Abundance 90th percentile
0.0
0.2
0.4
0.6
0.8
1.0
Ambo_sp.
Anaulus_australis
Aneumastus_subapiculatus
Attheya_longicornis
Aulacoseira_pusilla
Cerataulina_pelagica
Chaetoceros_cf._convolutus
Chaetoceros_cf._vixvisibilis
Chaetoceros_circinalis
Chaetoceros_costatus
Chaetoceros_dichaeta
Chaetoceros_dichatoensis
Chaetoceros_muelleri
Chaetoceros_socialis
Coscinodiscus_concinnus
Coscinodiscus_granii
Craticula_cuspidata
Cyclophora_tenuis
Cylindrotheca_closterium
Dactyliosolen_fragilissimus
Diatoma_vulgaris
Diploneis_parca
Diploneis_sp.
Eunotogramma_lunatum
Extubocellulus_spinifer
Fallacia_pygmaea
Fallacia_tenera
Fragilariopsis_kerguelensis
Halamphora_scatebra
Halamphora_sp.
Hannaea_arcus
Haslea_cf._howeana
Haslea_karadagensis
Hendeyella_dimeregrammopsis
Hyalodiscus_sp.
Hyalosira_delicatula
Licmophora_sp.
Microcostatus_sp.
Navicula_cf._salinarum
Navicula_sp.
Nitzschia_cf._palea
Nitzschia_cf._sigma
Nitzschia_cf._socialis
Nitzschia_lorenziana
Nitzschia_schefterae
Nitzschia_sp.
Paralia_sp.
Proboscia_truncata
Pseudo-nitzschia_calliantha
Pseudo-nitzschia_dolorosa
Pseudo-nitzschia_granii
Pseudo-nitzschia_multistriata
Pseudo-nitzschia_pseudodelicatissima
Pseudo-nitzschia_pungens_var._pungens
Pseudo-nitzschia_subfraudulenta
Pseudo-nitzschia_turgiduloides
Pseudosolenia_calcar-avis
Scoliopleura_peisonis
Sellaphora_terrestris
Serratifera_cf._andersonii
Shionodiscus_ritscheri
Staurophora_sp.
Thalassionema_sp.
Thalassiosira_eccentrica
Thalassiosira_hendeyi
Thalassiosira_punctigera
Thalassiosira_spinulifera
Trieres_sp.
Tursiocola_denysii

### TOP N Taxonomic Composition by Month: NPSW

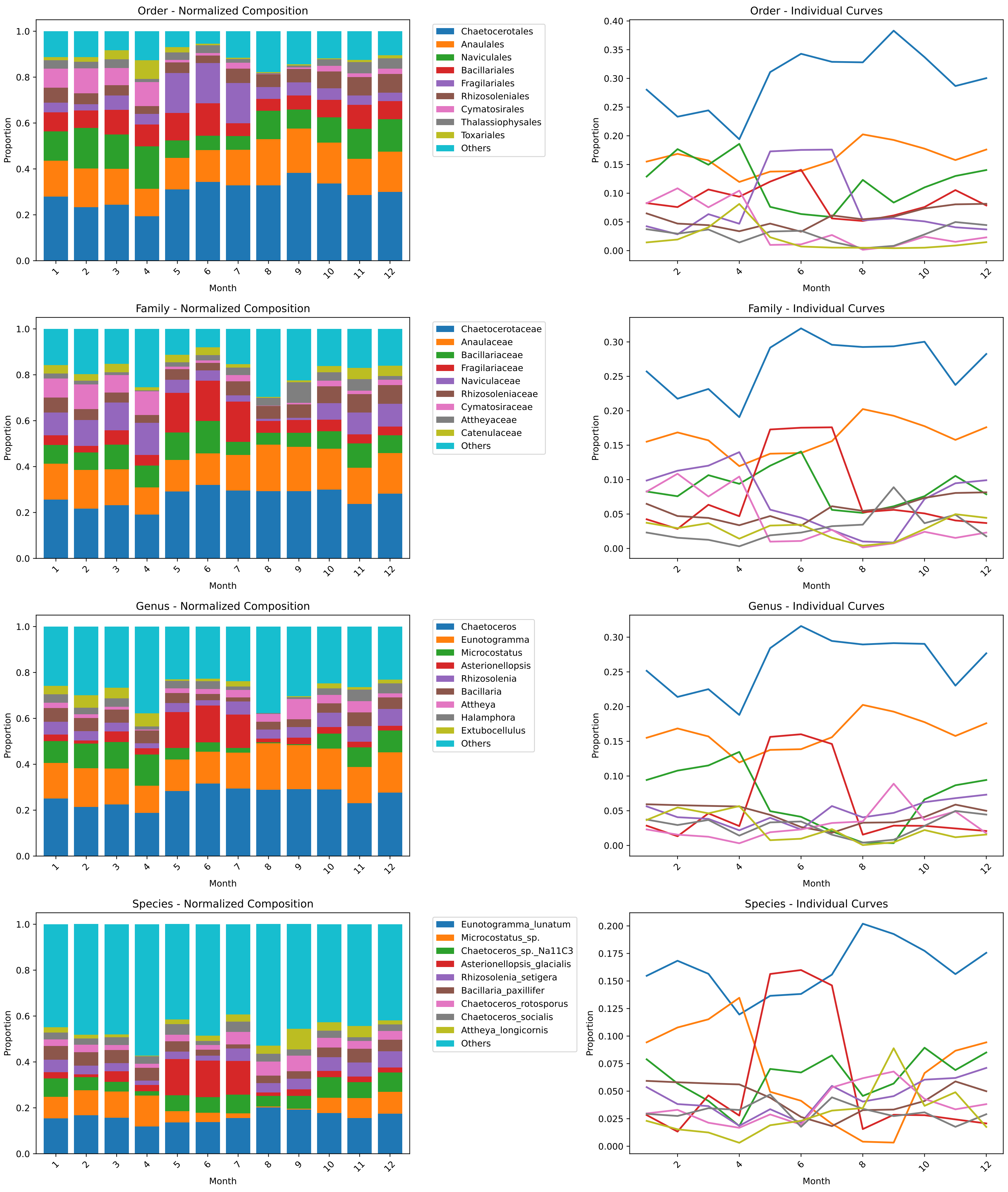

**NPSW**

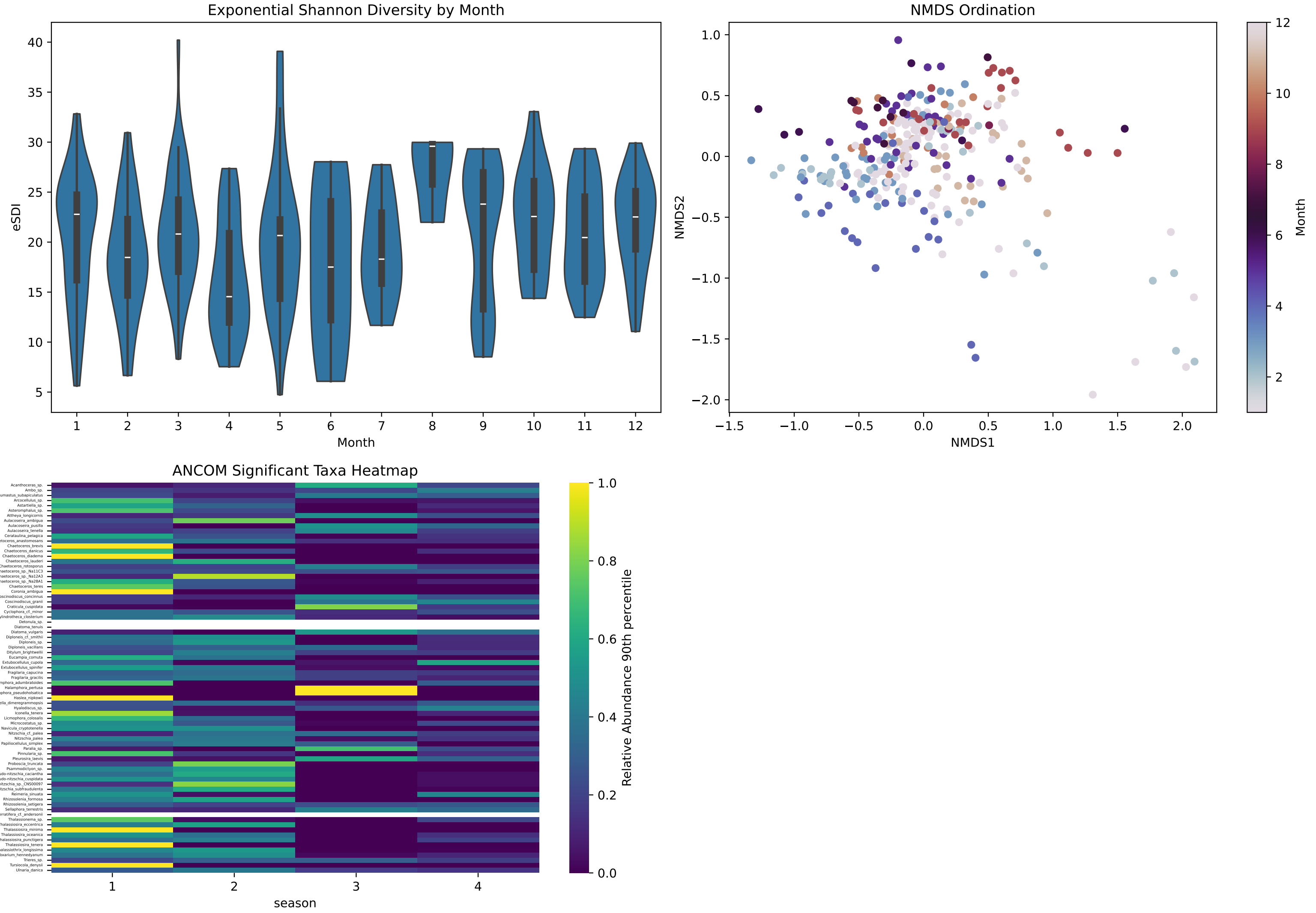
Exponential Shannon Diversity by Month
eSDI
Month
40
35
30
25
20
15
10
5
1
2
3
4
5
6
7
8
9
10
11
12
NMDS Ordination
NMDS2
NMDS1
1.0
0.5
0.0
−0.5
−1.0
−1.5
−2.0
−1.5
−1.0
−0.5
0.0
0.5
1.0
1.5
2.0
Month
12
10
8
6
4
2
ANCOM Significant Taxa Heatmap
T_Species
season
1
2
3
4
Relative Abundance 90th percentile
1.0
0.8
0.6
0.4
0.2
0.0

### TOP N Taxonomic Composition by Month: NPTG

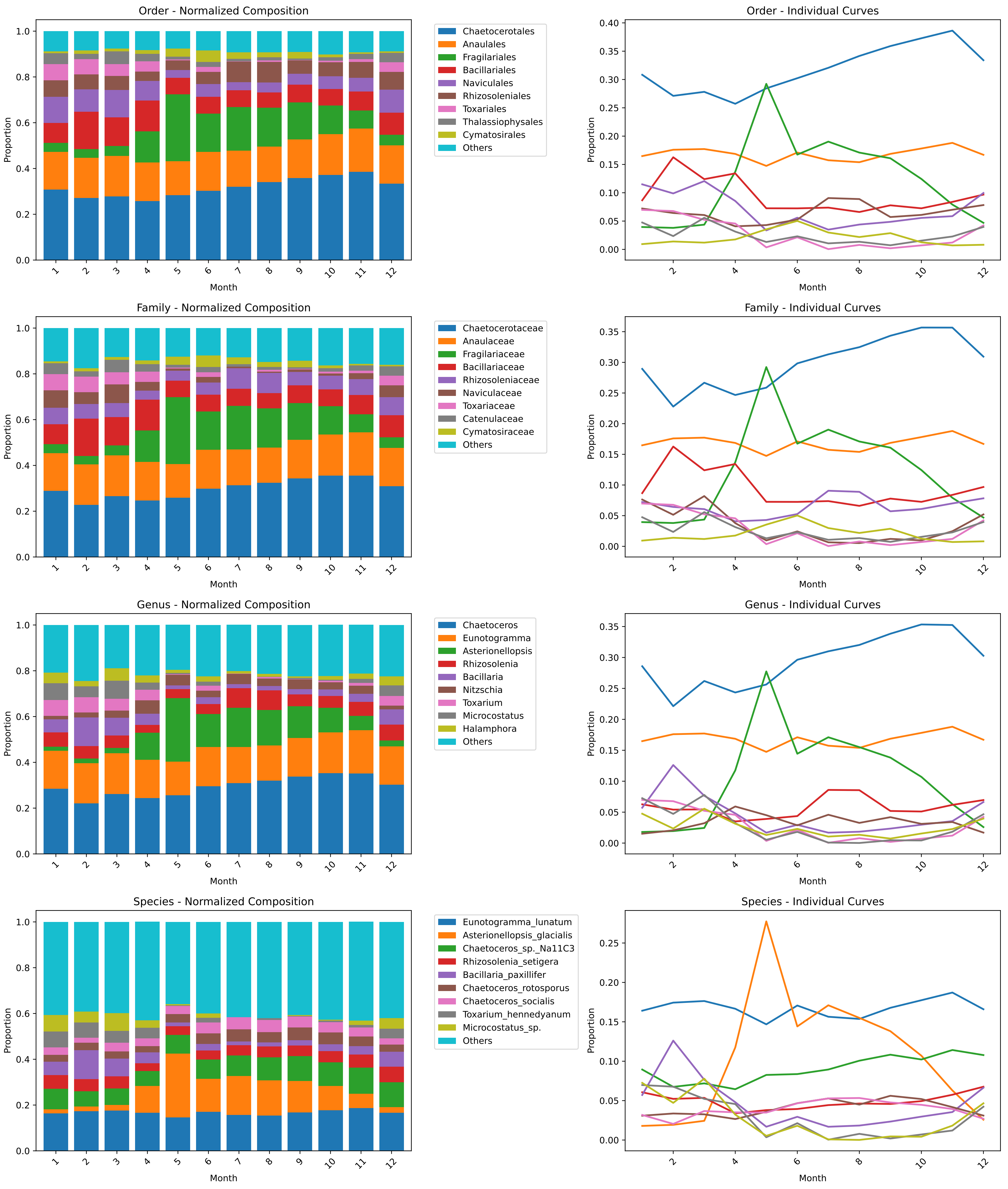

### NPTG

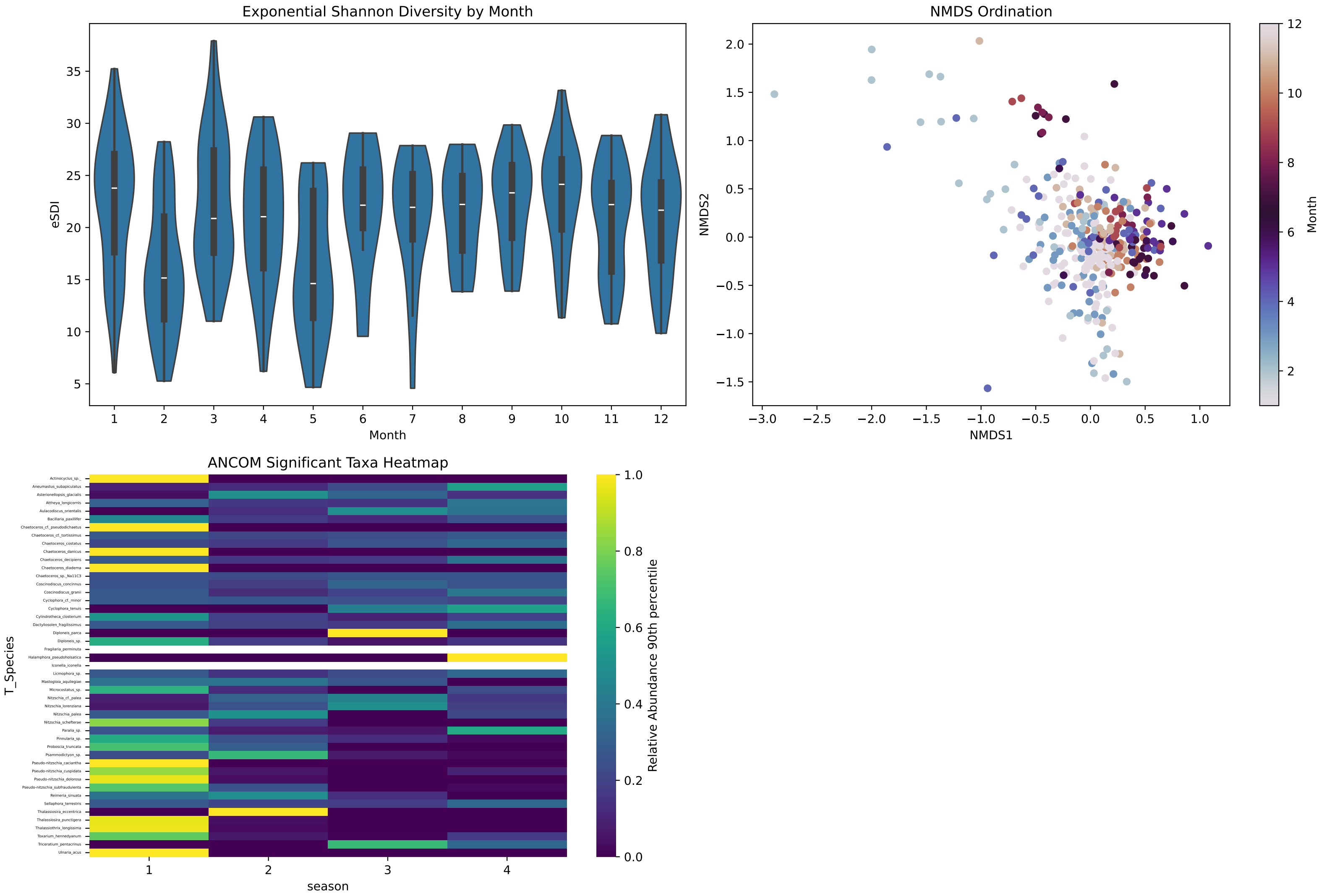

Exponential Shannon Diversity by Month
eSDI
Month
5
10
15
20
25
30
35
1
2
3
4
5
6
7
8
9
10
11
12
NMDS Ordination
NMDS2
NMDS1
−1.5
−1.0
−0.5
0.0
0.5
1.0
1.5
2.0
−3.0
−2.5
−2.0
−1.5
−1.0
−0.5
0.0
0.5
1.0
Month
2
4
6
8
10
12
ANCOM Significant Taxa Heatmap
T_Species
season
Relative Abundance 90th percentile
0.0
0.2
0.4
0.6
0.8
1.0
Actinocyclus_sp._
Aneumastus_subapiculatus
Asterionellopsis_glacialis
Attheya_longicornis
Aulacodiscus_orientalis
Bacillaria_paxillifer
Chaetoceros_cf._pseudodichaetus
Chaetoceros_cf._tortissimus
Chaetoceros_costatus
Chaetoceros_danicus
Chaetoceros_decipiens
Chaetoceros_diadema
Chaetoceros_sp._Na11C3
Coscinodiscus_concinnus
Coscinodiscus_granii
Cyclophora_cf._minor
Cyclophora_tenuis
Cylindrotheca_closterium
Dactyliosolen_fragilissimus
Diploneis_parca
Diploneis_sp.
Fragilaria_perminuta
Halamphora_pseudoholsatica
Iconella_iconella
Licmophora_sp.
Mastogloia_aquilegiae
Microcostatus_sp.
Nitzschia_cf._palea
Nitzschia_lorenziana
Nitzschia_palea
Nitzschia_schefterae
Paralia_sp.
Pinnularia_sp.
Proboscia_truncata
Psammodictyon_sp.
Pseudo-nitzschia_caciantha
Pseudo-nitzschia_cuspidata
Pseudo-nitzschia_dolorosa
Pseudo-nitzschia_subfraudulenta
Reimeria_sinuata
Sellaphora_terrestris
Thalassiosira_eccentrica
Thalassiosira_punctigera
Thalassiothrix_longissima
Toxarium_hennedyanum
Triceratium_pentacrinus
Ulnaria_acus
1
2
3
4

**TOP N Taxonomic Composition by Month: CCAL**

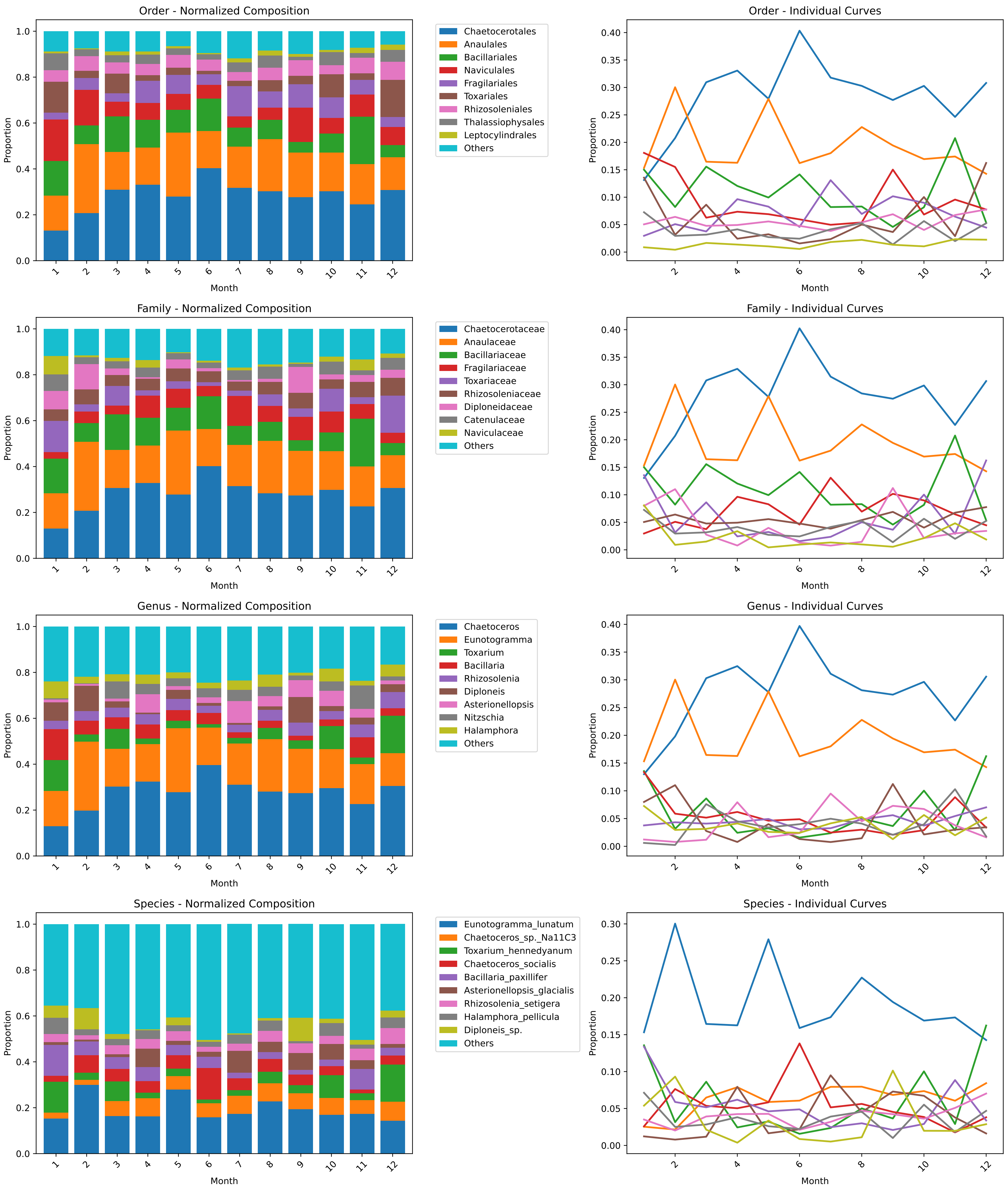

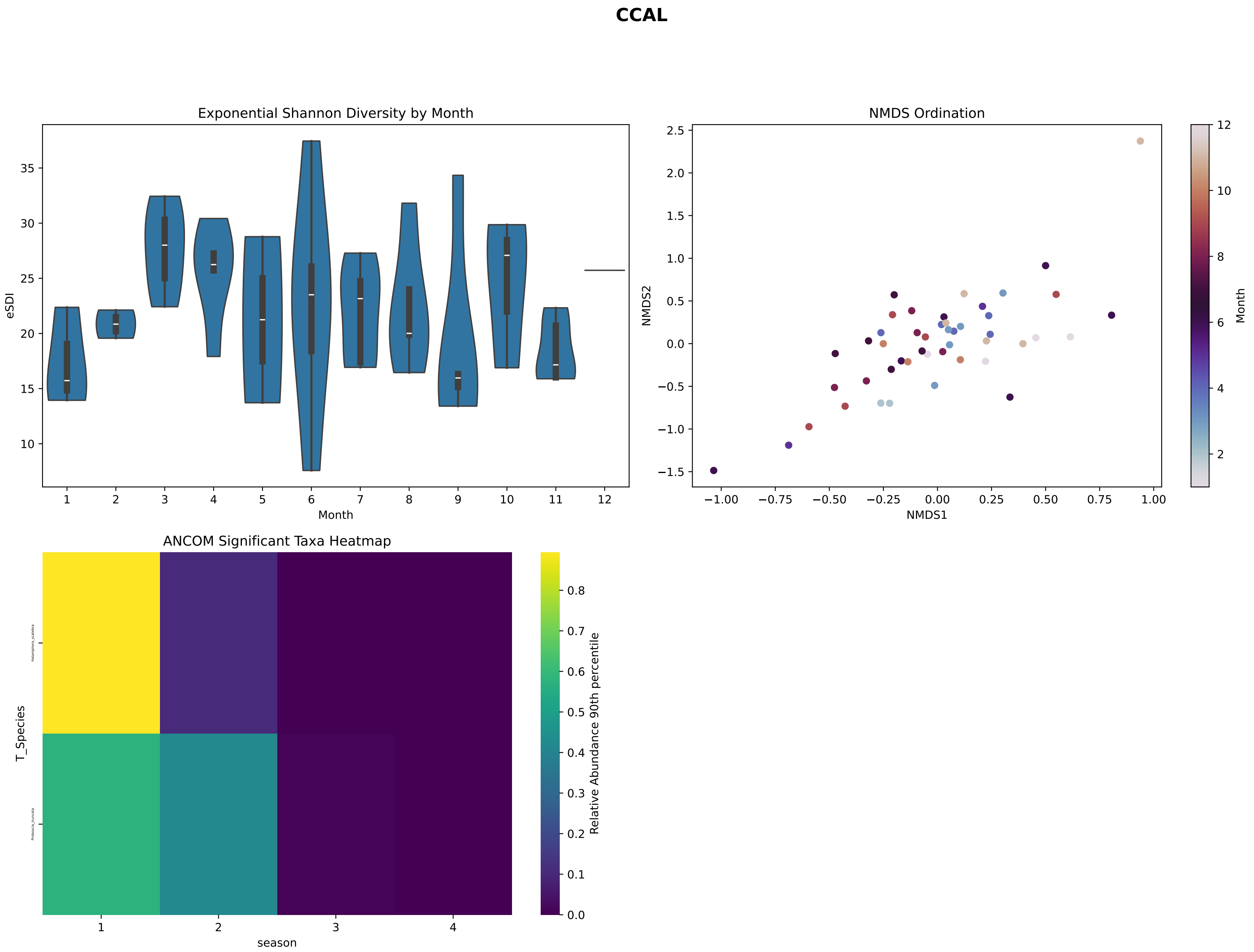
CCAL
Exponential Shannon Diversity by Month
eSDI
Month
1 2 3 4 5 6 7 8 9 10 11 12
NMDS Ordination
NMDS1
NMDS2
Month
ANCOM Significant Taxa Heatmap
T_Species
season
1 2 3 4
Relative Abundance 90th percentile

### TOP N Taxonomic Composition by Month: PNEC

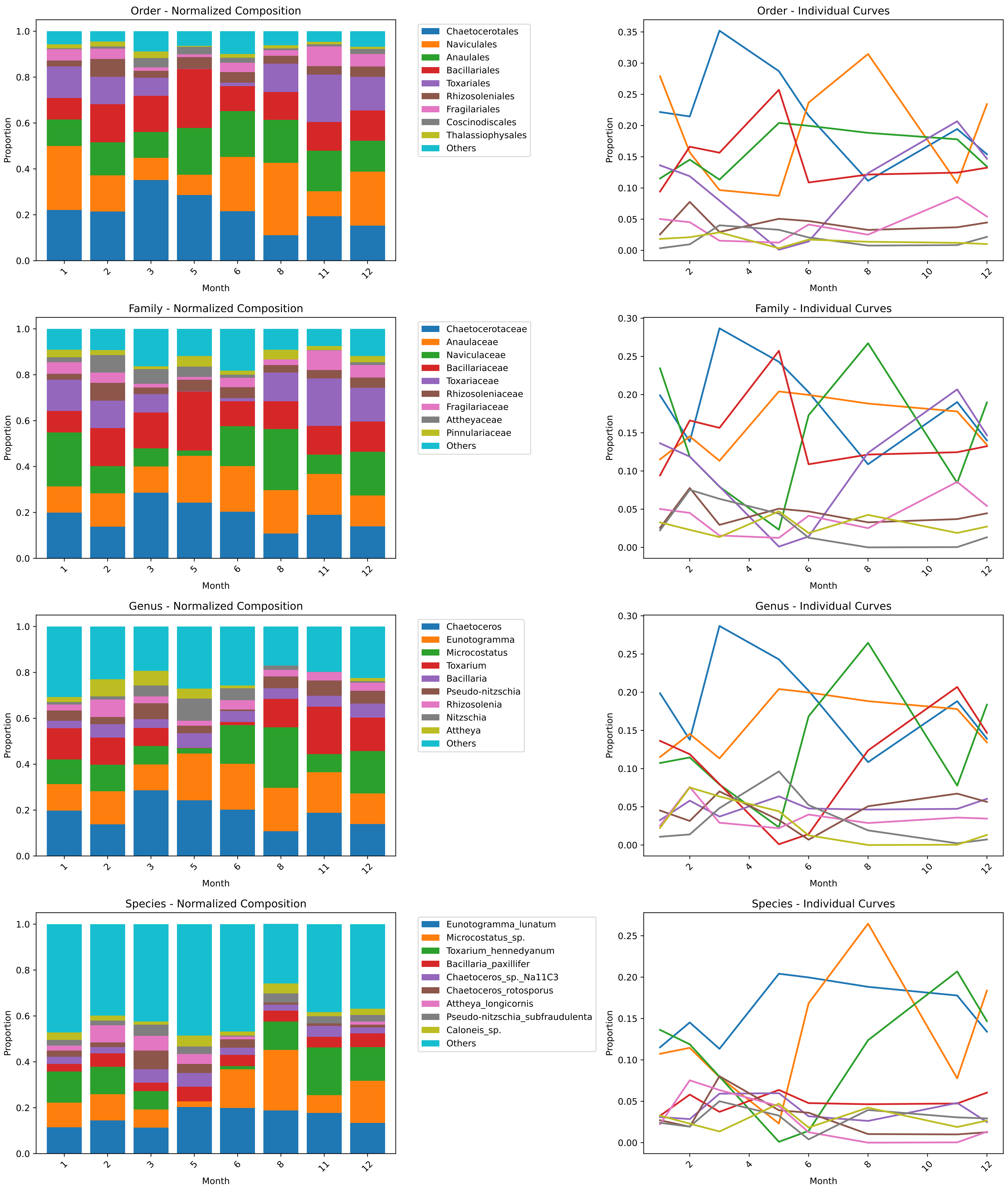

#### PNEC

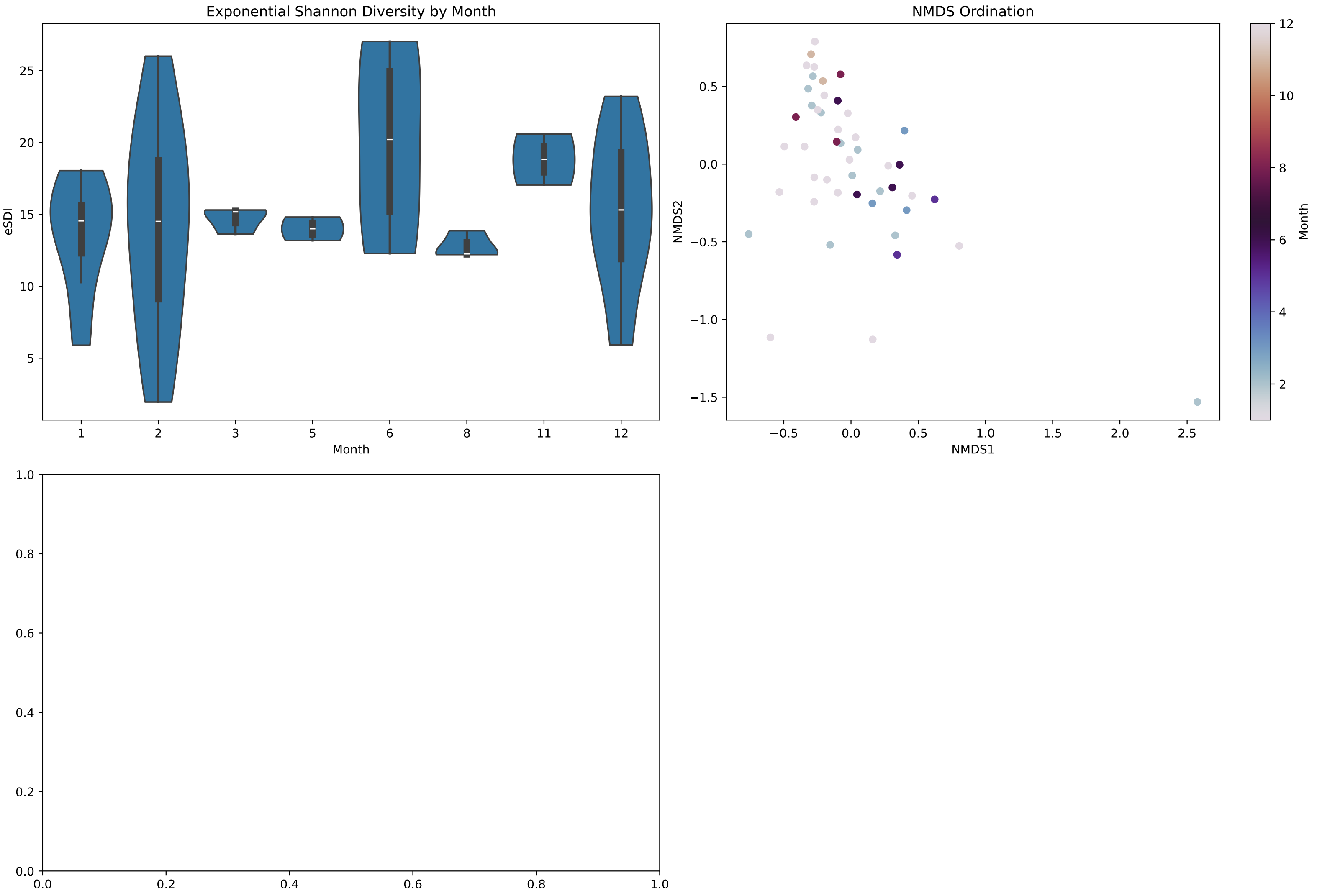
Exponential Shannon Diversity by Month
eSDI
Month
1 2 3 5 6 8 11 12
NMDS Ordination
NMDS1
NMDS2
Month

#### S7.3 Environmental drivers

Each double-page contains:

- VIF clustering based on dynamic complete linkage threshold (section S6),
- driver correlation matrix,
- RDA computed on hellinger-transformed abundances, informed by rescaled environmental drivers, Each sample is color-coded by month-of-year, and arrows are either green (significant, $p < 0.05$) or grey,
- variance partitioning on the top 4 expplanatory features from RDA,
- the top species responding to each driver,
- the previously-computed RDA k-means split into four clusters,
- for each of those clusters, its average environental characteristics, dominant species, and temporal sampling imbalance.

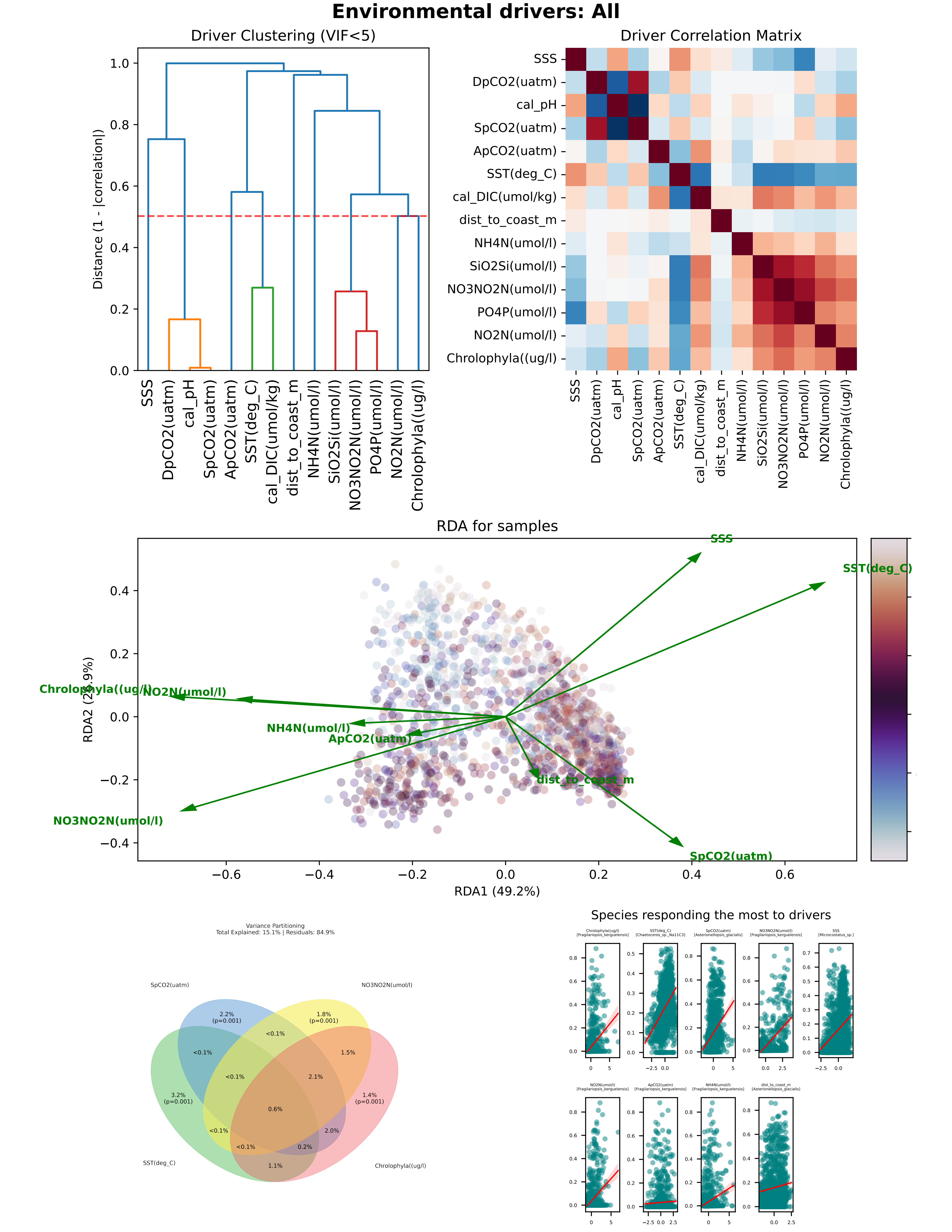
Environmental drivers: All
Driver Clustering (VIF<5)
Distance (1 - |correlation|)
SSS
DpCO2(uatm)
cal_pH
SpCO2(uatm)
ApCO2(uatm)
SST(deg_C)
cal_DIC(umol/kg)
dist_to_coast_m
NH4N(umol/l)
SiO2Si(umol/l)
NO3NO2N(umol/l)
PO4P(umol/l)
NO2N(umol/l)
Chrolophyla((ug/l)
Driver Correlation Matrix
RDA for samples
RDA1 (49.2%)
RDA2 (2?.9%)
Variance Partitioning
Total Explained: 15.1% | Residuals: 84.9%
2.2% (p=0.001)
1.8% (p=0.001)
3.2% (p=0.001)
1.4% (p=0.001)
<0.1%
1.5%
2.1%
0.6%
2.0%
0.2%
1.1%
Species responding the most to drivers

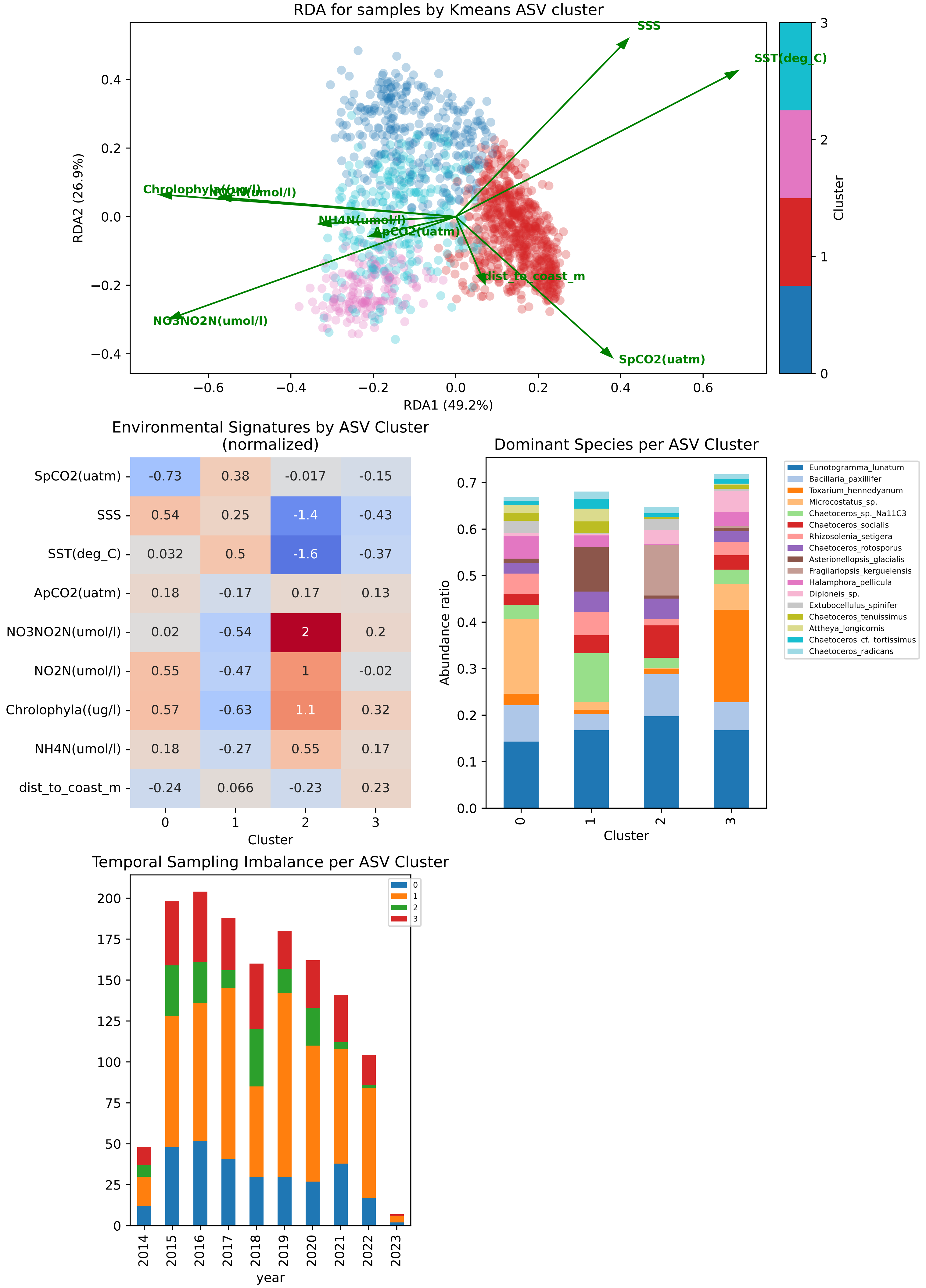

RDA for samples by Kmeans ASV cluster
SSS
SST(deg_C)
Chrolophyla((ug/l)
NO2N(umol/l)
NH4N(umol/l)
ApCO2(uatm)
dist_to_coast_m
NO3NO2N(umol/l)
SpCO2(uatm)
RDA1 (49.2%)
RDA2 (26.9%)
Cluster
Environmental Signatures by ASV Cluster
(normalized)
SpCO2(uatm) -0.73 0.38 -0.017 -0.15
SSS 0.54 0.25 -1.4 -0.43
SST(deg_C) 0.032 0.5 -1.6 -0.37
ApCO2(uatm) 0.18 -0.17 0.17 0.13
NO3NO2N(umol/l) 0.02 -0.54 2 0.2
NO2N(umol/l) 0.55 -0.47 1 -0.02
Chrolophyla((ug/l) 0.57 -0.63 1.1 0.32
NH4N(umol/l) 0.18 -0.27 0.55 0.17
dist_to_coast_m -0.24 0.066 -0.23 0.23
Cluster
Dominant Species per ASV Cluster
Abundance ratio
Cluster
Eunotogramma_lunatum
Bacillaria_paxillifer
Toxarium_hennedyanum
Microcostatus_sp.
Chaetoceros_sp._Na11C3
Chaetoceros_socialis
Rhizosolenia_setigera
Chaetoceros_rotosporus
Asterionellopsis_glacialis
Fragilariopsis_kerguelensis
Halamphora_pellicula
Diploneis_sp.
Extubocellulus_spinifer
Chaetoceros_tenuissimus
Attheya_longicornis
Chaetoceros_cf._tortissimus
Chaetoceros_radicans
Temporal Sampling Imbalance per ASV Cluster
year

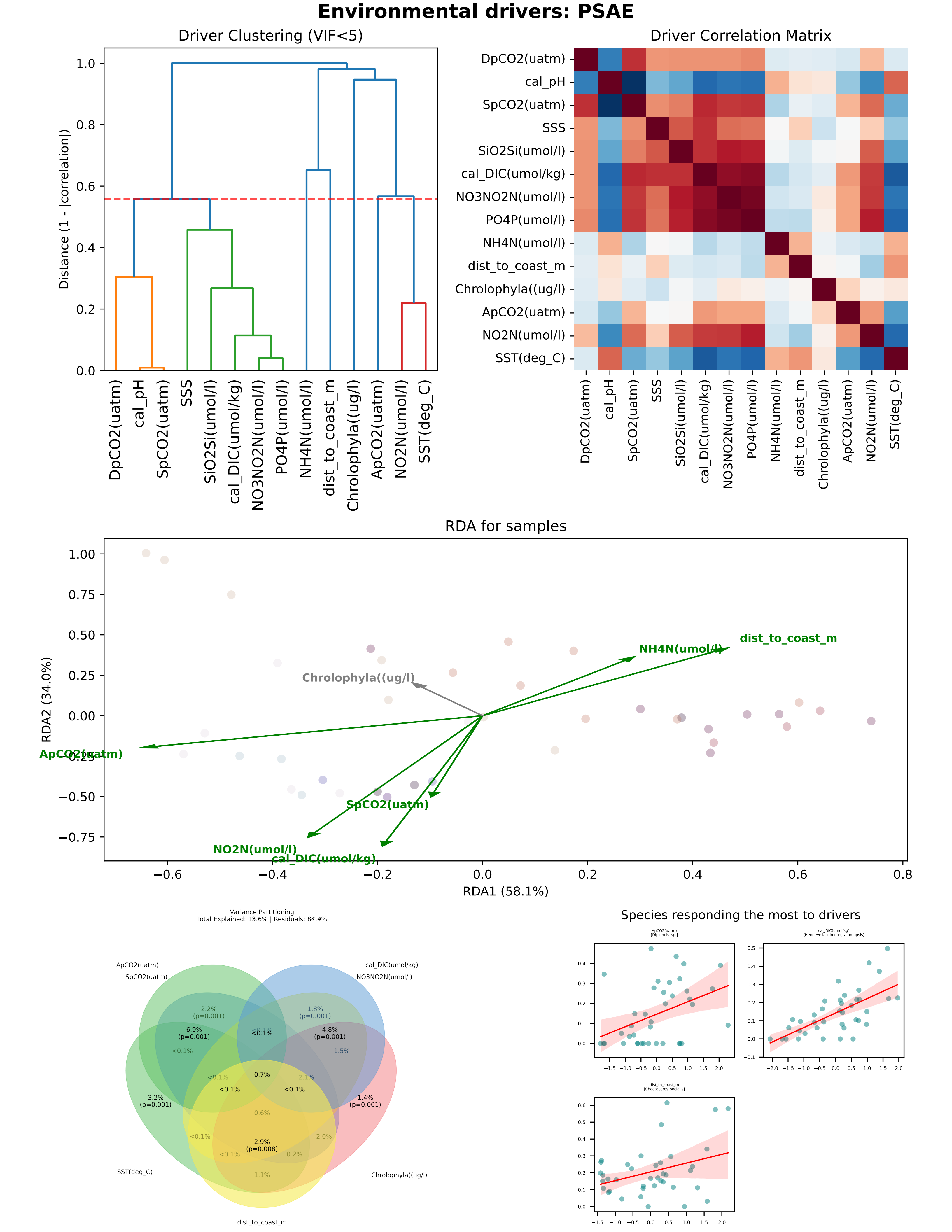

Environmental drivers: PSAE
Driver Clustering (VIF<5)
Distance (1 - |correlation|)
DpCO2(uatm)
cal_pH
SpCO2(uatm)
SSS
SiO2Si(umol/l)
cal_DIC(umol/kg)
NO3NO2N(umol/l)
PO4P(umol/l)
NH4N(umol/l)
dist_to_coast_m
Chrolophyla((ug/l)
ApCO2(uatm)
NO2N(umol/l)
SST(deg_C)
Driver Correlation Matrix
RDA for samples
RDA1 (58.1%)
RDA2 (34.0%)
Variance Partitioning
2.2% (p=0.001)
1.8% (p=0.001)
6.9% (p=0.001)
4.8% (p=0.001)
3.2% (p=0.001)
1.4% (p=0.001)
2.9% (p=0.008)
0.7%
Species responding the most to drivers
ApCO2(uatm) [Diploneis_sp.]
cal_DIC(umol/kg) [Hendeyella_dimeregrammopsis]
dist_to_coast_m [Chaetoceros_socialis]

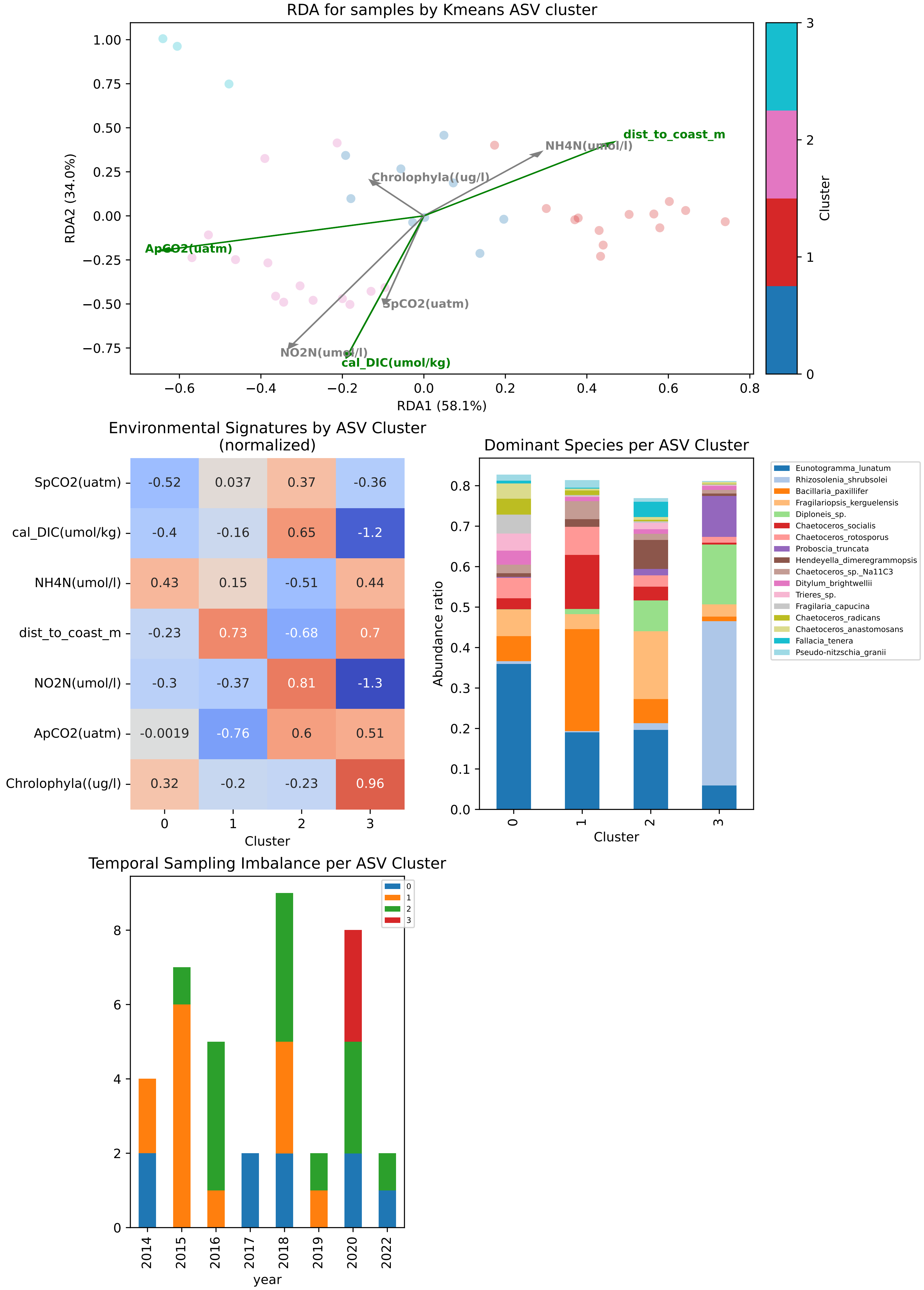
RDA for samples by Kmeans ASV cluster
RDA1 (58.1%)
RDA2 (34.0%)
Cluster
dist_to_coast_m
NH4N(umol/l)
Chrolophyla((ug/l)
ApCO2(uatm)
SpCO2(uatm)
NO2N(umol/l)
cal_DIC(umol/kg)
Environmental Signatures by ASV Cluster
(normalized)
SpCO2(uatm) -0.52 0.037 0.37 -0.36
cal_DIC(umol/kg) -0.4 -0.16 0.65 -1.2
NH4N(umol/l) 0.43 0.15 -0.51 0.44
dist_to_coast_m -0.23 0.73 -0.68 0.7
NO2N(umol/l) -0.3 -0.37 0.81 -1.3
ApCO2(uatm) -0.0019 -0.76 0.6 0.51
Chrolophyla((ug/l) 0.32 -0.2 -0.23 0.96
Cluster
Dominant Species per ASV Cluster
Abundance ratio
Cluster
Eunotogramma_lunatum
Rhizosolenia_shrubsolei
Bacillaria_paxillifer
Fragilariopsis_kerguelensis
Diploneis_sp.
Chaetoceros_socialis
Chaetoceros_rotosporus
Proboscia_truncata
Hendeyella_dimeregrammopsis
Chaetoceros_sp._Na11C3
Ditylum_brightwellii
Trieres_sp.
Fragilaria_capucina
Chaetoceros_radicans
Chaetoceros_anastomosans
Fallacia_tenera
Pseudo-nitzschia_granii
Temporal Sampling Imbalance per ASV Cluster
year
2014 2015 2016 2017 2018 2019 2020 2022

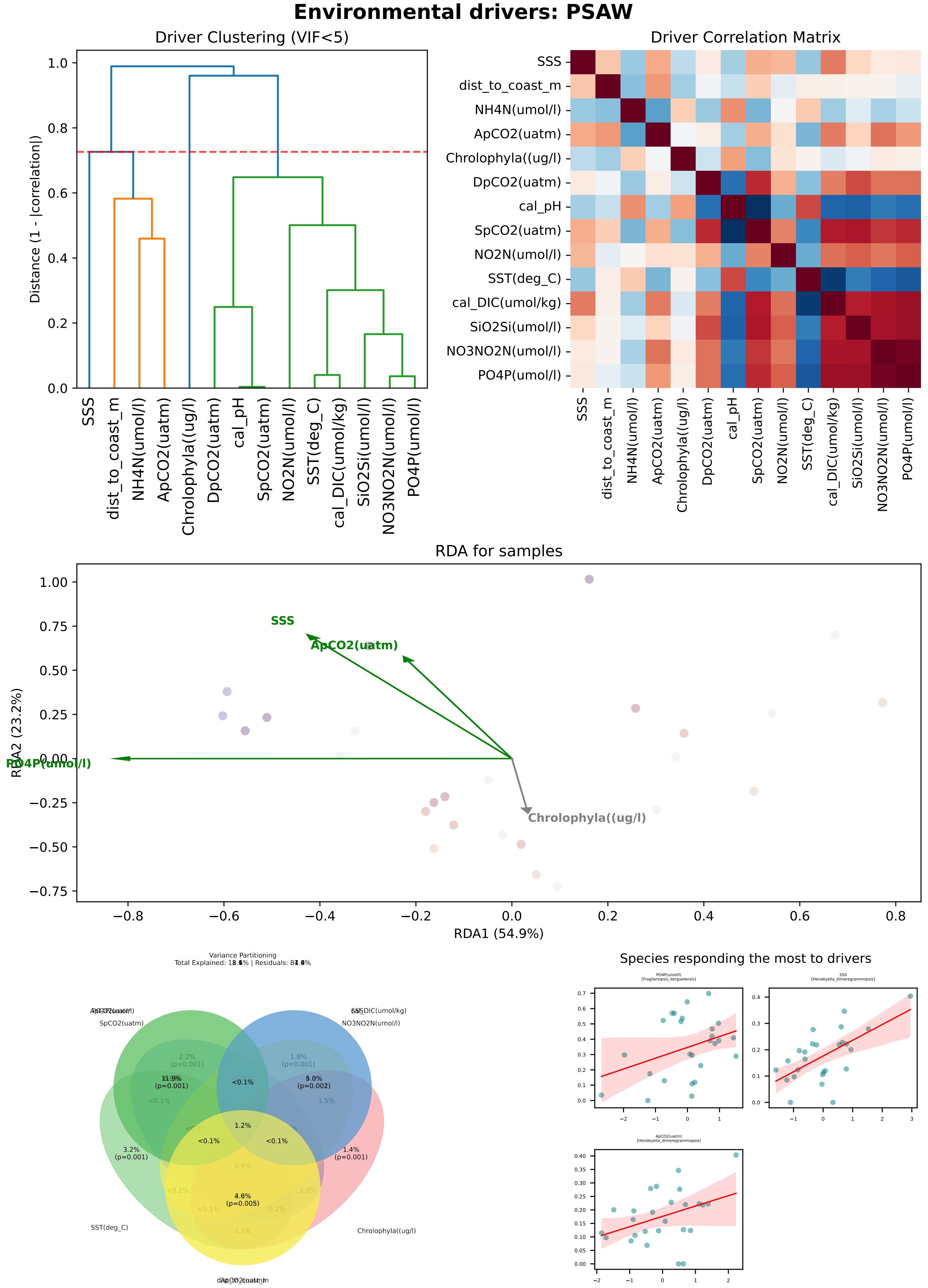

Environmental drivers: PSAW
Driver Clustering (VIF<5)
Distance (1 - |correlation|)
SSS
dist_to_coast_m
NH4N(umol/l)
ApCO2(uatm)
Chrolophyla((ug/l)
DpCO2(uatm)
cal_pH
SpCO2(uatm)
NO2N(umol/l)
SST(deg_C)
cal_DIC(umol/kg)
SiO2Si(umol/l)
NO3NO2N(umol/l)
PO4P(umol/l)
Driver Correlation Matrix
RDA for samples
RDA1 (54.9%)
RDA2 (23.2%)
Variance Partitioning
Species responding the most to drivers

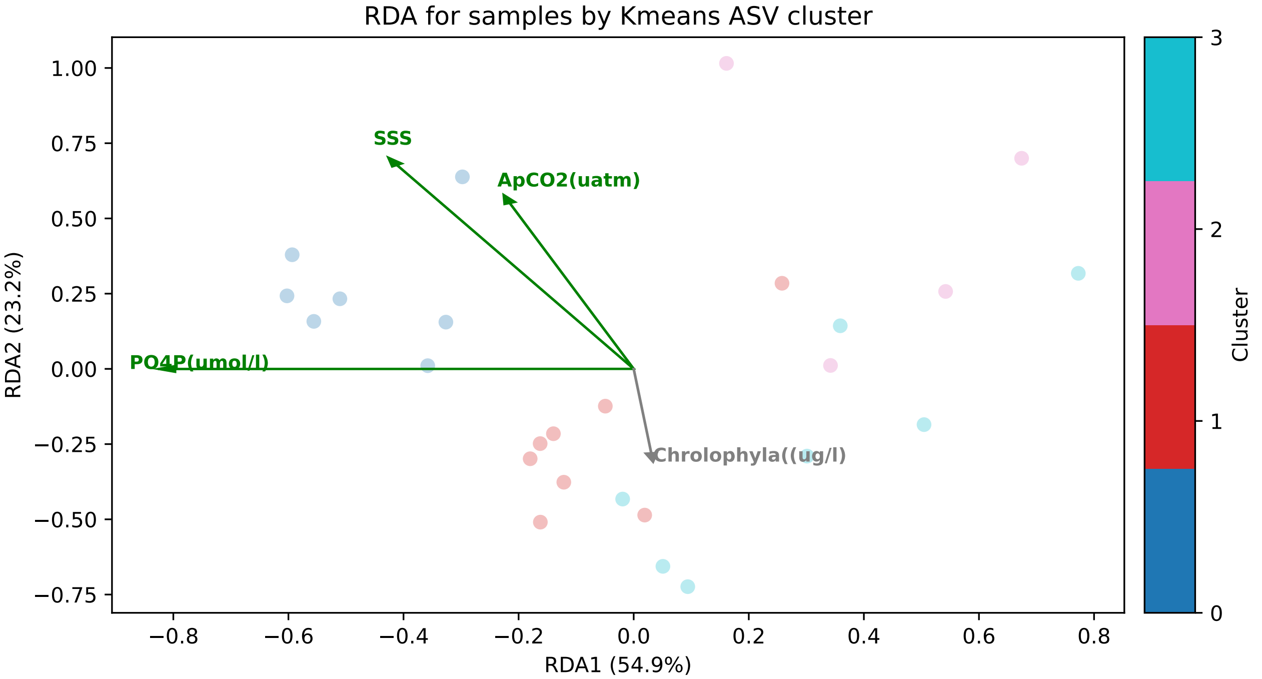

RDA for samples by Kmeans ASV cluster
SSS
ApCO2(uatm)
PO4P(umol/l)
Chrolophyla((ug/l)
RDA1 (54.9%)
RDA2 (23.2%)
Cluster


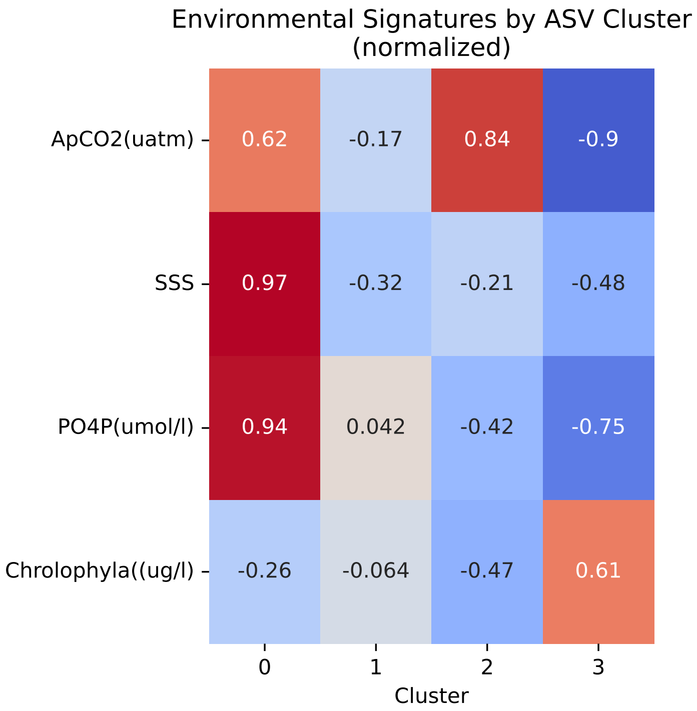

Environmental Signatures by ASV Cluster
(normalized)
ApCO2(uatm)
SSS
PO4P(umol/l)
Chrolophyla((ug/l)
0.62
-0.17
0.84
-0.9
0.97
-0.32
-0.21
-0.48
0.94
0.042
-0.42
-0.75
-0.26
-0.064
-0.47
0.61
Cluster


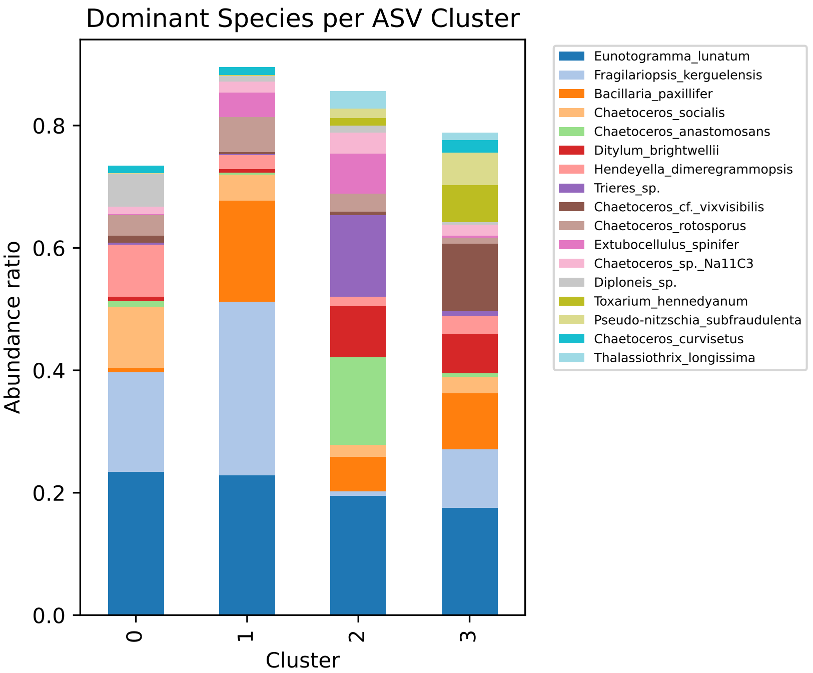

Dominant Species per ASV Cluster
Abundance ratio
Cluster
Eunotogramma_lunatum
Fragilariopsis_kerguelensis
Bacillaria_paxillifer
Chaetoceros_socialis
Chaetoceros_anastomosans
Ditylum_brightwellii
Hendeyella_dimeregrammopsis
Trieres_sp.
Chaetoceros_cf._vixvisibilis
Chaetoceros_rotosporus
Extubocellulus_spinifer
Chaetoceros_sp._Na11C3
Diploneis_sp.
Toxarium_hennedyanum
Pseudo-nitzschia_subfraudulenta
Chaetoceros_curvisetus
Thalassiothrix_longissima


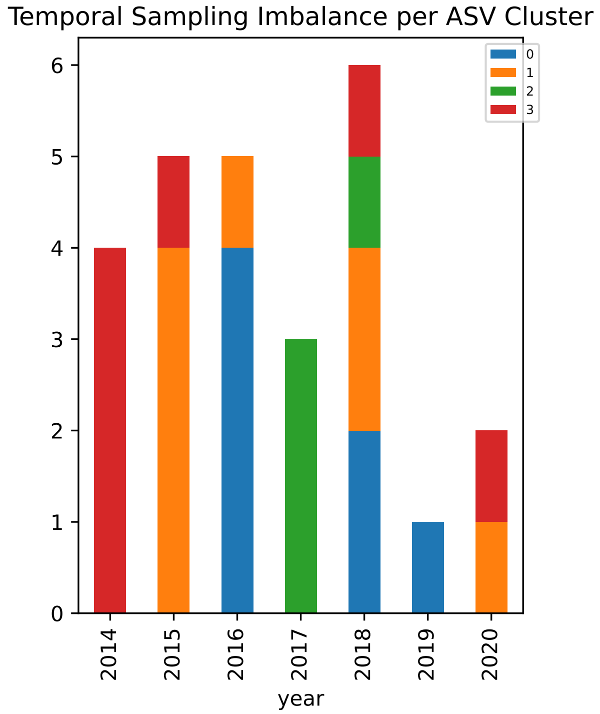

Temporal Sampling Imbalance per ASV Cluster
year

**Environmental drivers: KURO**

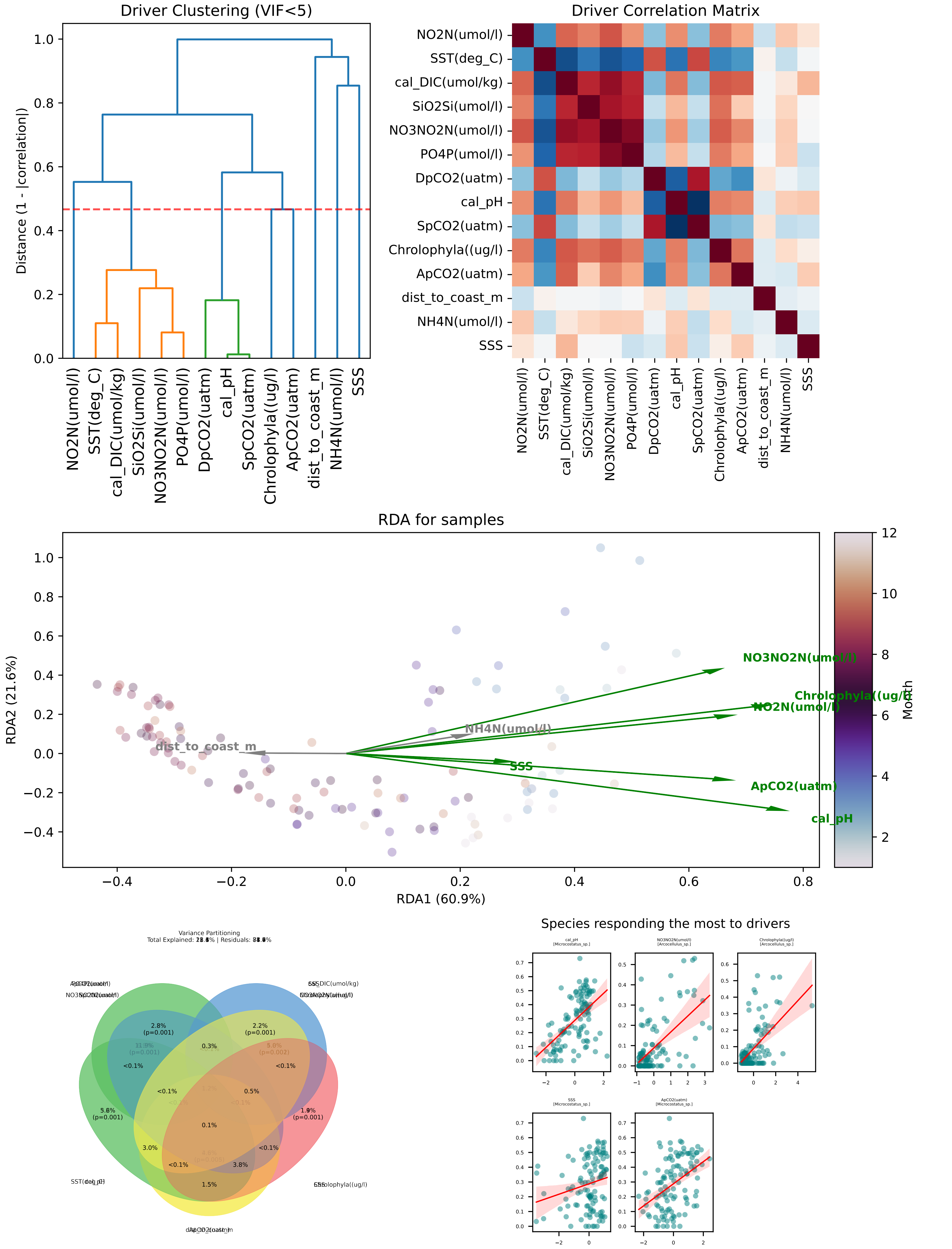

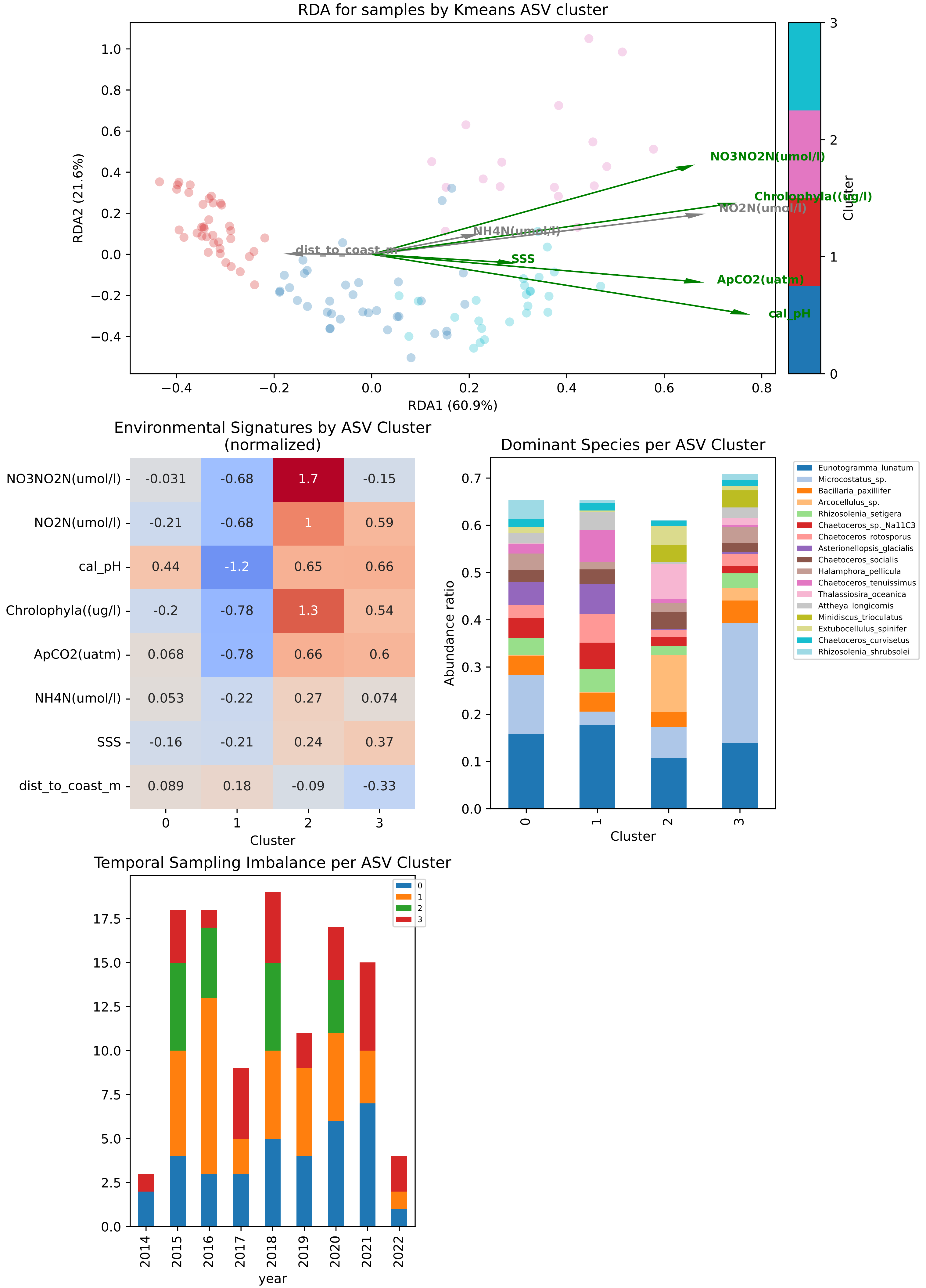
RDA for samples by Kmeans ASV cluster
RDA1 (60.9%)
RDA2 (21.6%)
Cluster
NO3NO2N(umol/l)
Chrolophyla((ug/l)
NO2N(umol/l)
NH4N(umol/l)
dist_to_coast_m
SSS
ApCO2(uatm)
cal_pH
Environmental Signatures by ASV Cluster
(normalized)
NO3NO2N(umol/l) -0.031 -0.68 1.7 -0.15
NO2N(umol/l) -0.21 -0.68 1 0.59
cal_pH 0.44 -1.2 0.65 0.66
Chrolophyla((ug/l) -0.2 -0.78 1.3 0.54
ApCO2(uatm) 0.068 -0.78 0.66 0.6
NH4N(umol/l) 0.053 -0.22 0.27 0.074
SSS -0.16 -0.21 0.24 0.37
dist_to_coast_m 0.089 0.18 -0.09 -0.33
Cluster
Dominant Species per ASV Cluster
Abundance ratio
Cluster
Eunotogramma_lunatum
Microcostatus_sp.
Bacillaria_paxillifer
Arcocellulus_sp.
Rhizosolenia_setigera
Chaetoceros_sp._Na11C3
Chaetoceros_rotosporus
Asterionellopsis_glacialis
Chaetoceros_socialis
Halamphora_pellicula
Chaetoceros_tenuissimus
Thalassiosira_oceanica
Attheya_longicornis
Minidiscus_trioculatus
Extubocellulus_spinifer
Chaetoceros_curvisetus
Rhizosolenia_shrubsolei
Temporal Sampling Imbalance per ASV Cluster
year

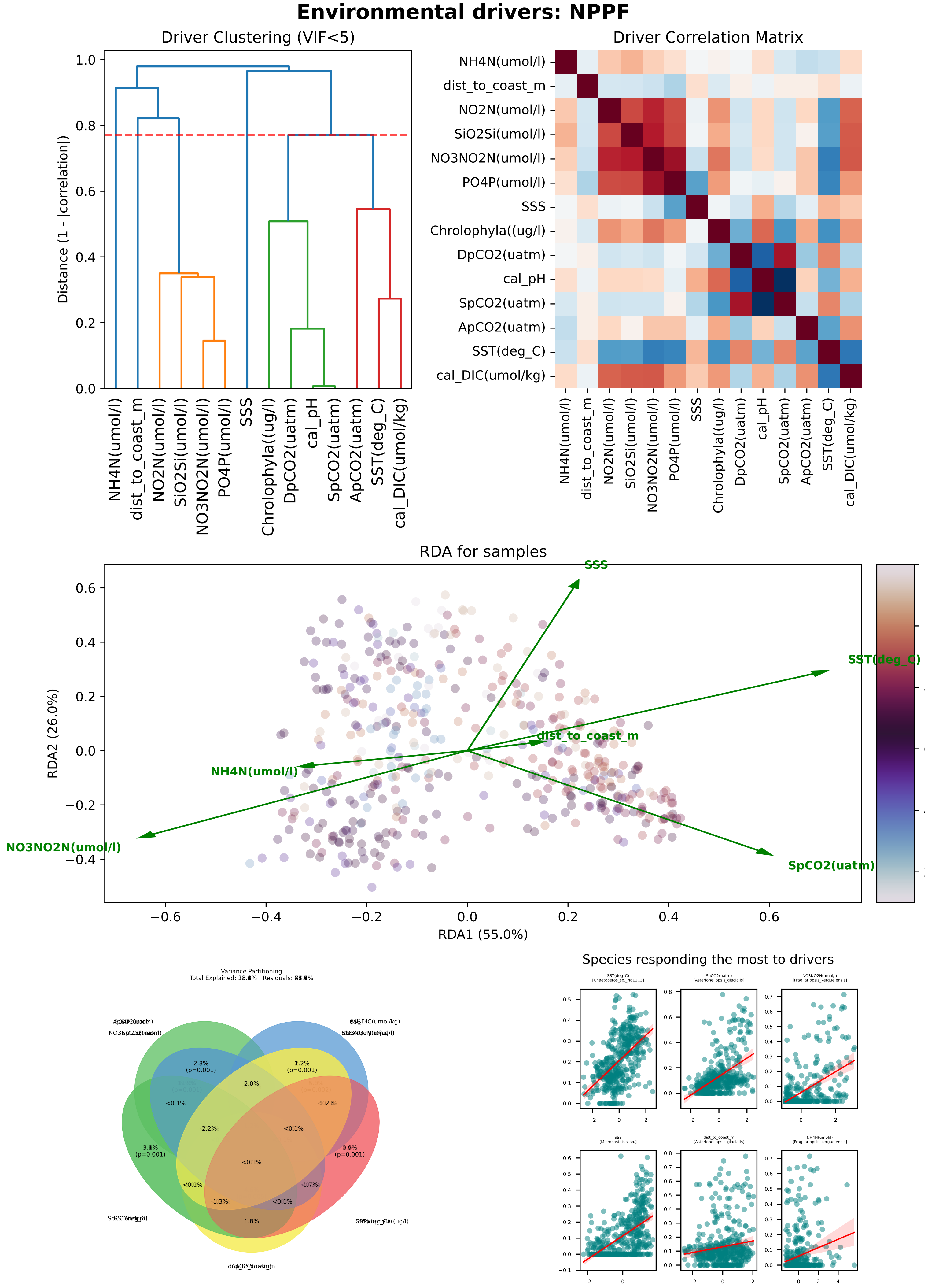
Environmental drivers: NPPF
Driver Clustering (VIF<5)
Distance (1 - |correlation|)
NH4N(umol/l)
dist_to_coast_m
NO2N(umol/l)
SiO2Si(umol/l)
NO3NO2N(umol/l)
PO4P(umol/l)
SSS
Chrolophyla((ug/l)
DpCO2(uatm)
cal_pH
SpCO2(uatm)
ApCO2(uatm)
SST(deg_C)
cal_DIC(umol/kg)
Driver Correlation Matrix
RDA for samples
RDA1 (55.0%)
RDA2 (26.0%)
Variance Partitioning
Species responding the most to drivers
SST(deg_C) [Chaetoceros_sp._Na11C3]
SpCO2(uatm) [Asterionellopsis_glacialis]
NO3NO2N(umol/l) [Fragilariopsis_kerguelensis]
SSS [Micrococtatus_sp.]
dist_to_coast_m [Asterionellopsis_glacialis]
NH4N(umol/l) [Fragilariopsis_kerguelensis]

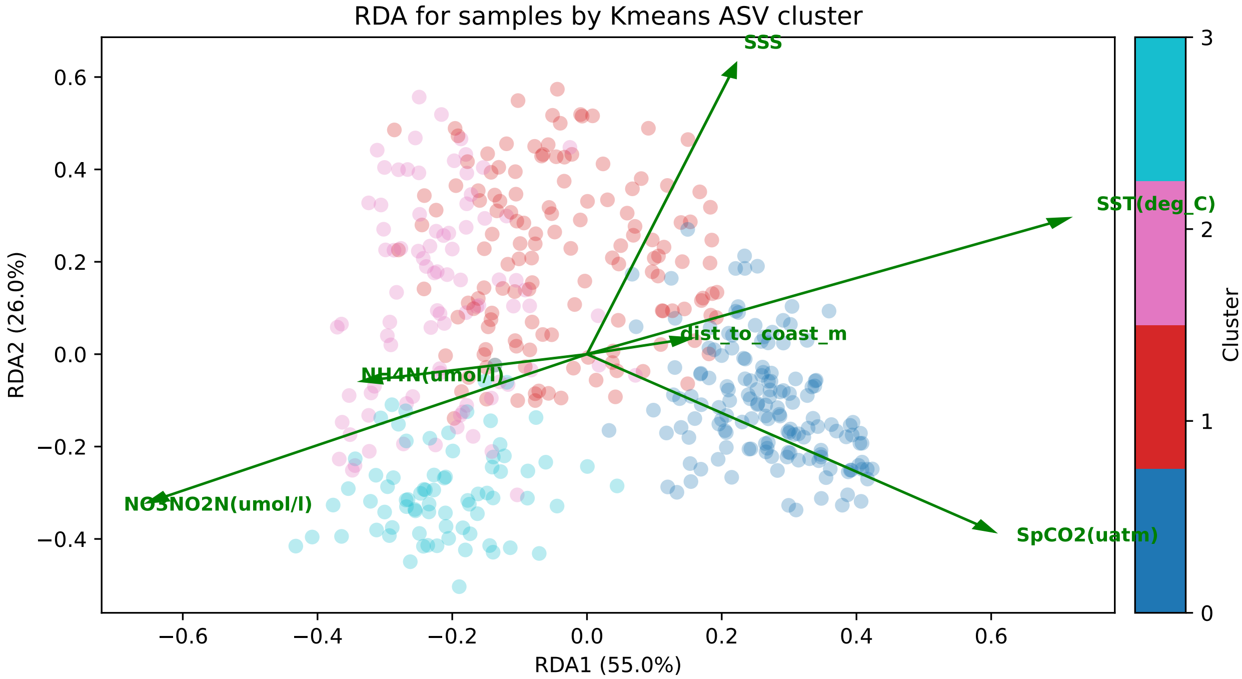
RDA for samples by Kmeans ASV cluster
SSS
SST(deg_C)
dist_to_coast_m
NH4N(umol/l)
NO3NO2N(umol/l)
SpCO2(uatm)
RDA2 (26.0%)
RDA1 (55.0%)
Cluster

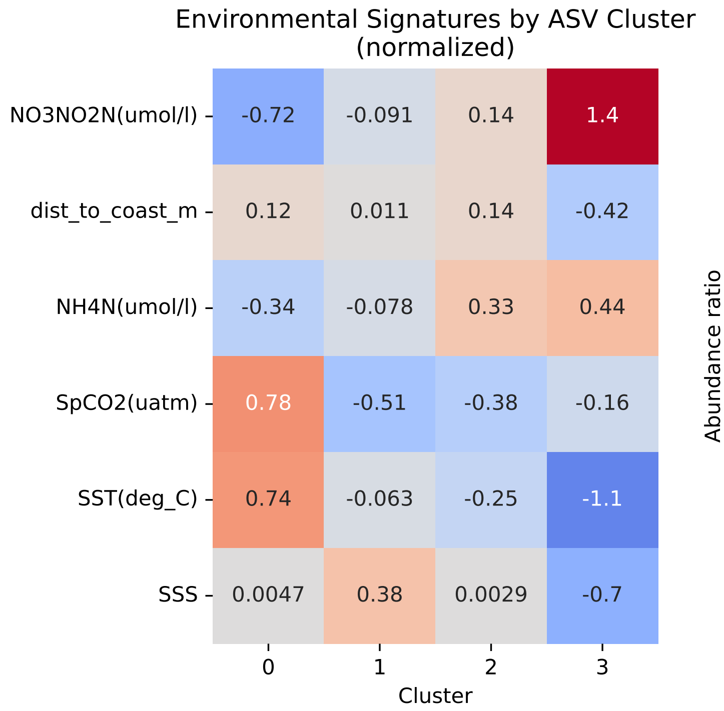
Environmental Signatures by ASV Cluster (normalized)
NO3NO2N(umol/l) -0.72 -0.091 0.14 1.4
dist_to_coast_m 0.12 0.011 0.14 -0.42
NH4N(umol/l) -0.34 -0.078 0.33 0.44
SpCO2(uatm) 0.78 -0.51 -0.38 -0.16
SST(deg_C) 0.74 -0.063 -0.25 -1.1
SSS 0.0047 0.38 0.0029 -0.7
Cluster

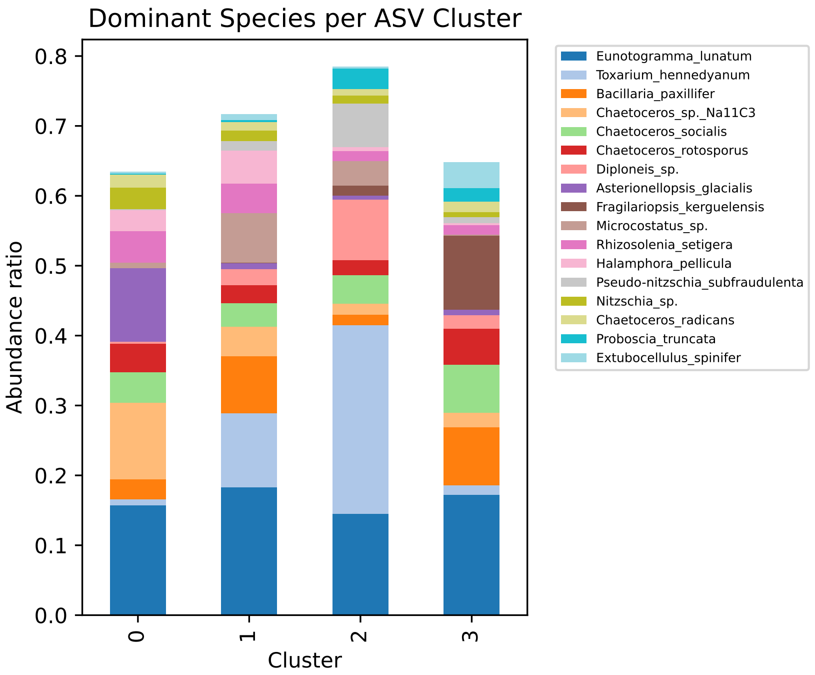
Dominant Species per ASV Cluster
Abundance ratio
Cluster
Eunotogramma_lunatum
Toxarium_hennedyanum
Bacillaria_paxillifer
Chaetoceros_sp._Na11C3
Chaetoceros_socialis
Chaetoceros_rotosporus
Diploneis_sp.
Asterionellopsis_glacialis
Fragilariopsis_kerguelensis
Microcostatus_sp.
Rhizosolenia_setigera
Halamphora_pellicula
Pseudo-nitzschia_subfraudulenta
Nitzschia_sp.
Chaetoceros_radicans
Proboscia_truncata
Extubocellulus_spinifer

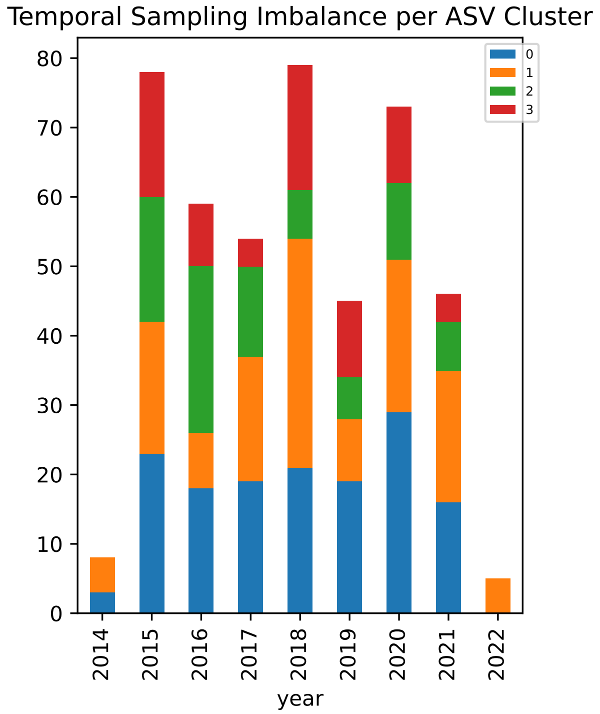
Temporal Sampling Imbalance per ASV Cluster
year

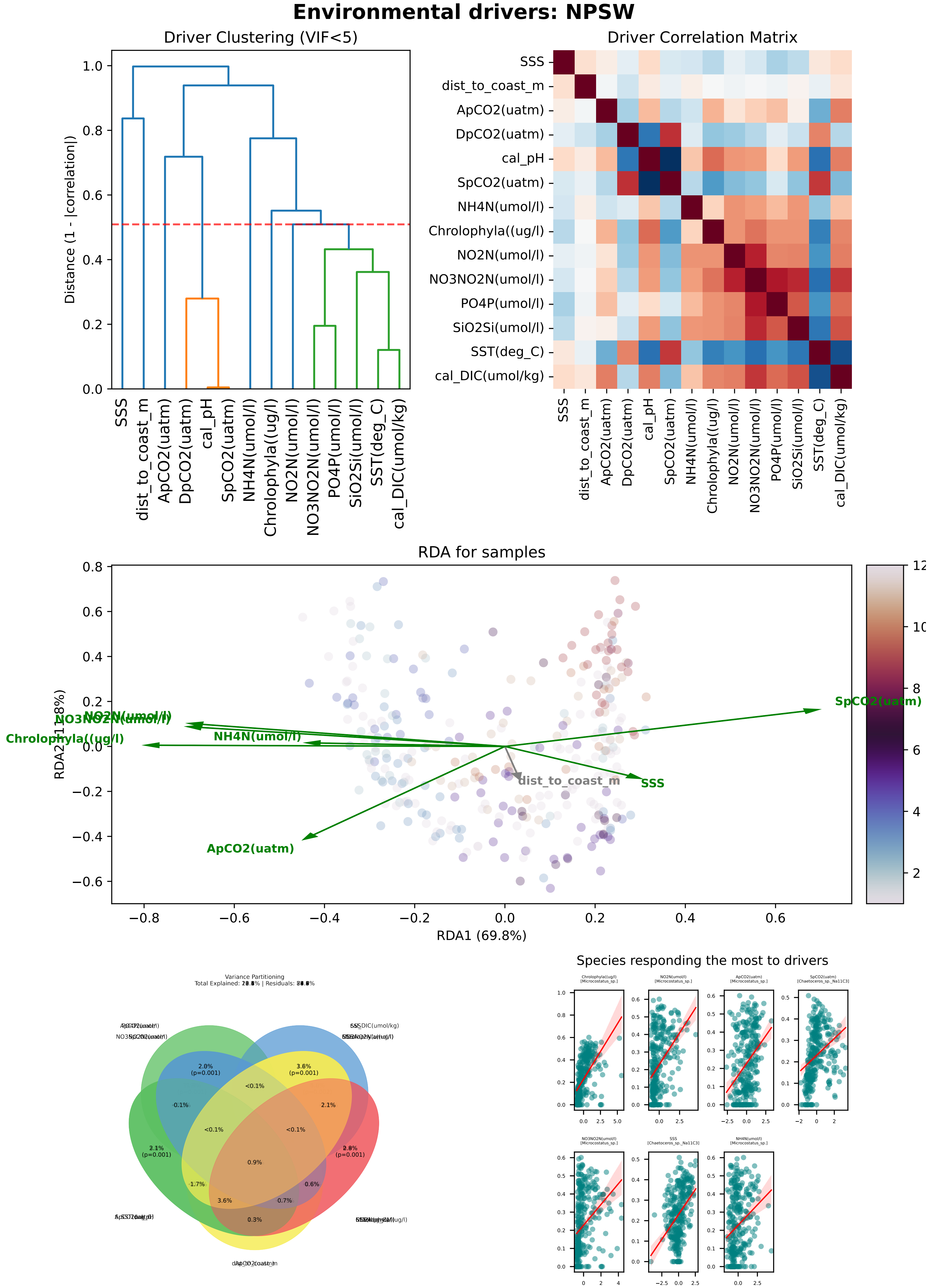
Environmental drivers: NPSW
Driver Clustering (VIF<5)
Distance (1 - |correlation|)
SSS
dist_to_coast_m
ApCO2(uatm)
DpCO2(uatm)
cal_pH
SpCO2(uatm)
NH4N(umol/l)
Chrolophyla((ug/l)
NO2N(umol/l)
NO3NO2N(umol/l)
PO4P(umol/l)
SiO2Si(umol/l)
SST(deg_C)
cal_DIC(umol/kg)
Driver Correlation Matrix
RDA for samples
RDA1 (69.8%)
SpCO2(uatm)
NO2N(umol/l)
NO3NO2N(umol/l)
Chrolophyla((ug/l)
NH4N(umol/l)
dist_to_coast_m
SSS
ApCO2(uatm)
Variance Partitioning
Species responding the most to drivers

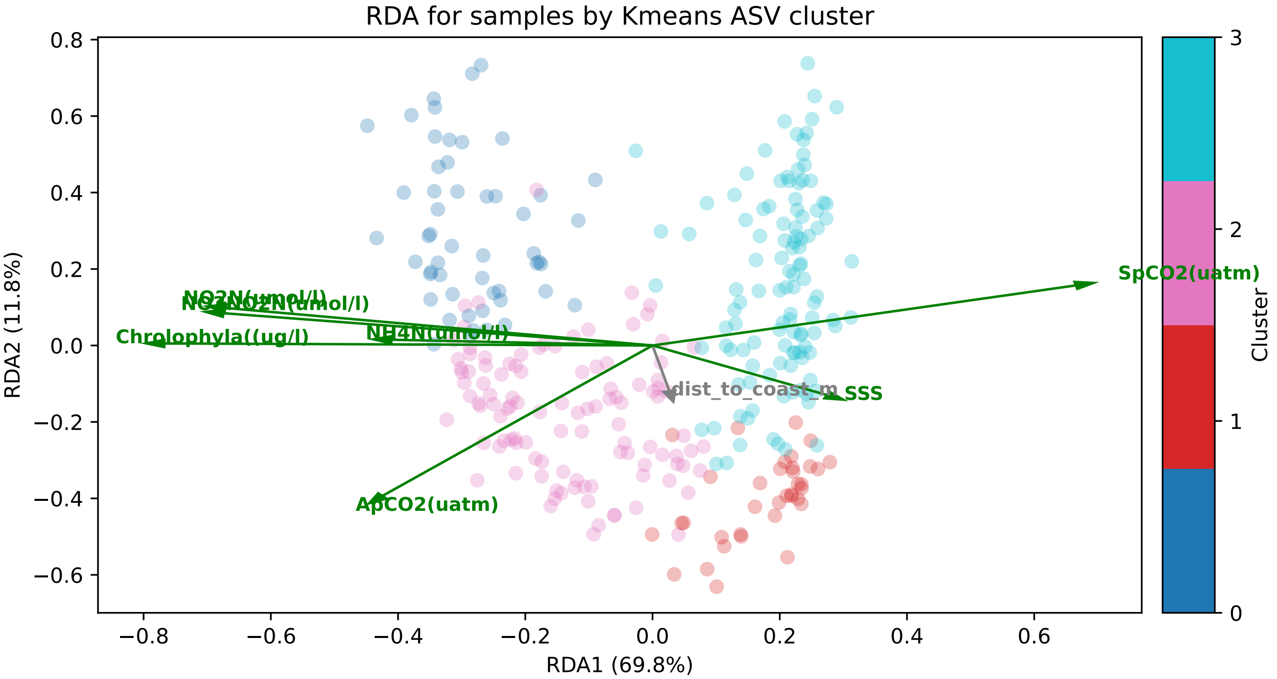
RDA for samples by Kmeans ASV cluster
SpCO2(uatm)
NO2N(umol/l)
NO3NO2N(umol/l)
Chrolophyla((ug/l)
NH4N(umol/l)
dist_to_coast_m
SSS
ApCO2(uatm)
RDA2 (11.8%)
RDA1 (69.8%)
Cluster

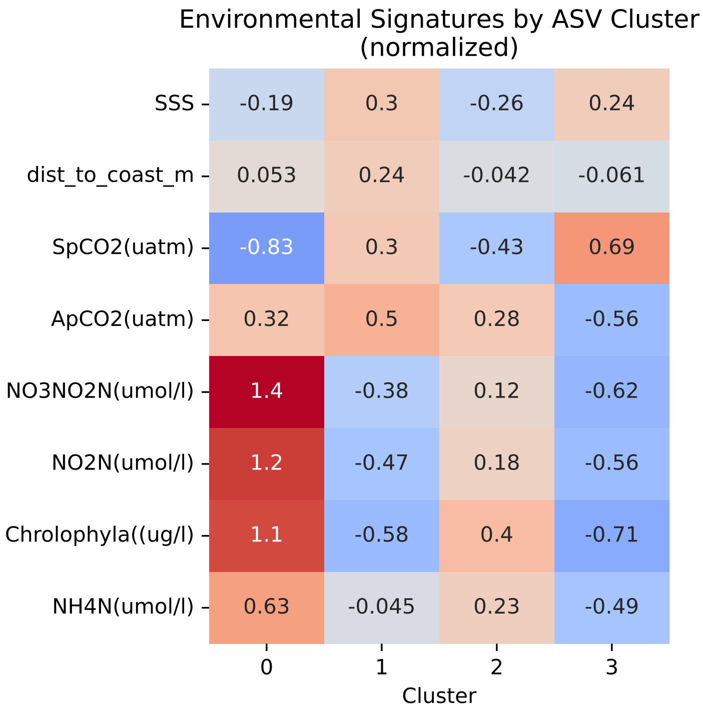
Environmental Signatures by ASV Cluster
(normalized)
SSS
dist_to_coast_m
SpCO2(uatm)
ApCO2(uatm)
NO3NO2N(umol/l)
NO2N(umol/l)
Chrolophyla((ug/l)
NH4N(umol/l)
-0.19 0.3 -0.26 0.24
0.053 0.24 -0.042 -0.061
-0.83 0.3 -0.43 0.69
0.32 0.5 0.28 -0.56
1.4 -0.38 0.12 -0.62
1.2 -0.47 0.18 -0.56
1.1 -0.58 0.4 -0.71
0.63 -0.045 0.23 -0.49
Cluster

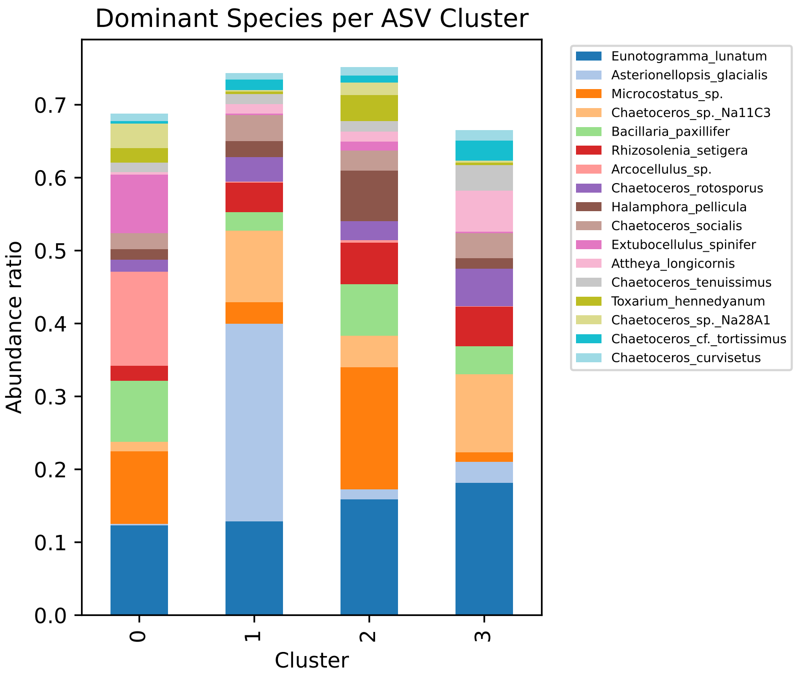
Dominant Species per ASV Cluster
Abundance ratio
Cluster
Eunotogramma_lunatum
Asterionellopsis_glacialis
Microcostatus_sp.
Chaetoceros_sp._Na11C3
Bacillaria_paxillifer
Rhizosolenia_setigera
Arcocellulus_sp.
Chaetoceros_rotosporus
Halamphora_pellicula
Chaetoceros_socialis
Extubocellulus_spinifer
Attheya_longicornis
Chaetoceros_tenuissimus
Toxarium_hennedyanum
Chaetoceros_sp._Na28A1
Chaetoceros_cf._tortissimus
Chaetoceros_curvisetus

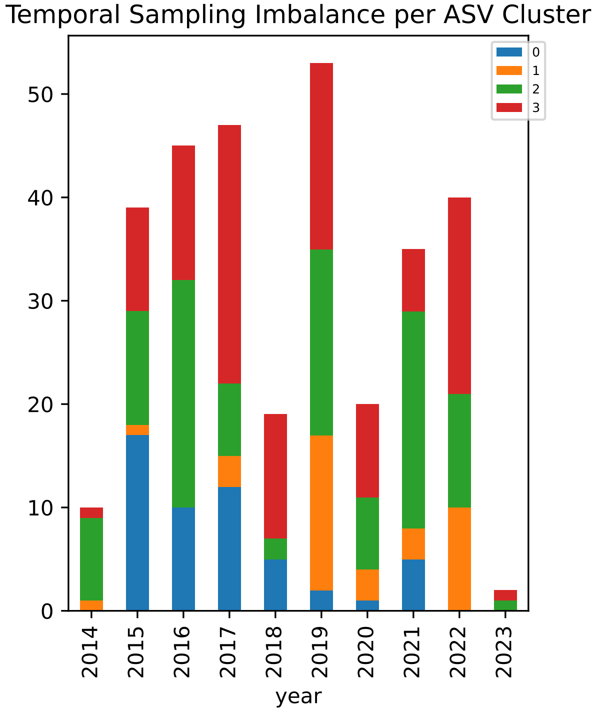
Temporal Sampling Imbalance per ASV Cluster
year

### Environmental drivers: NPTG

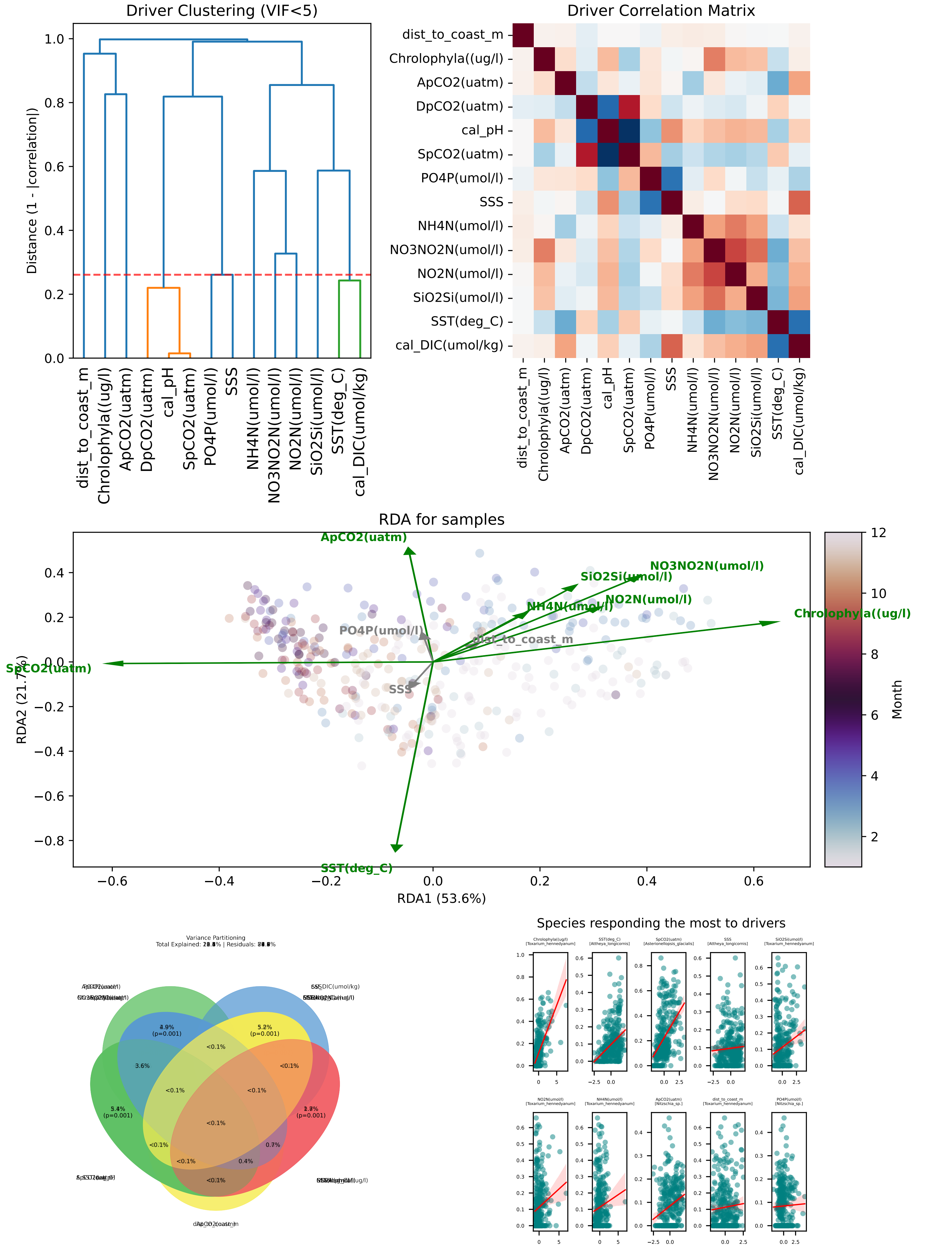

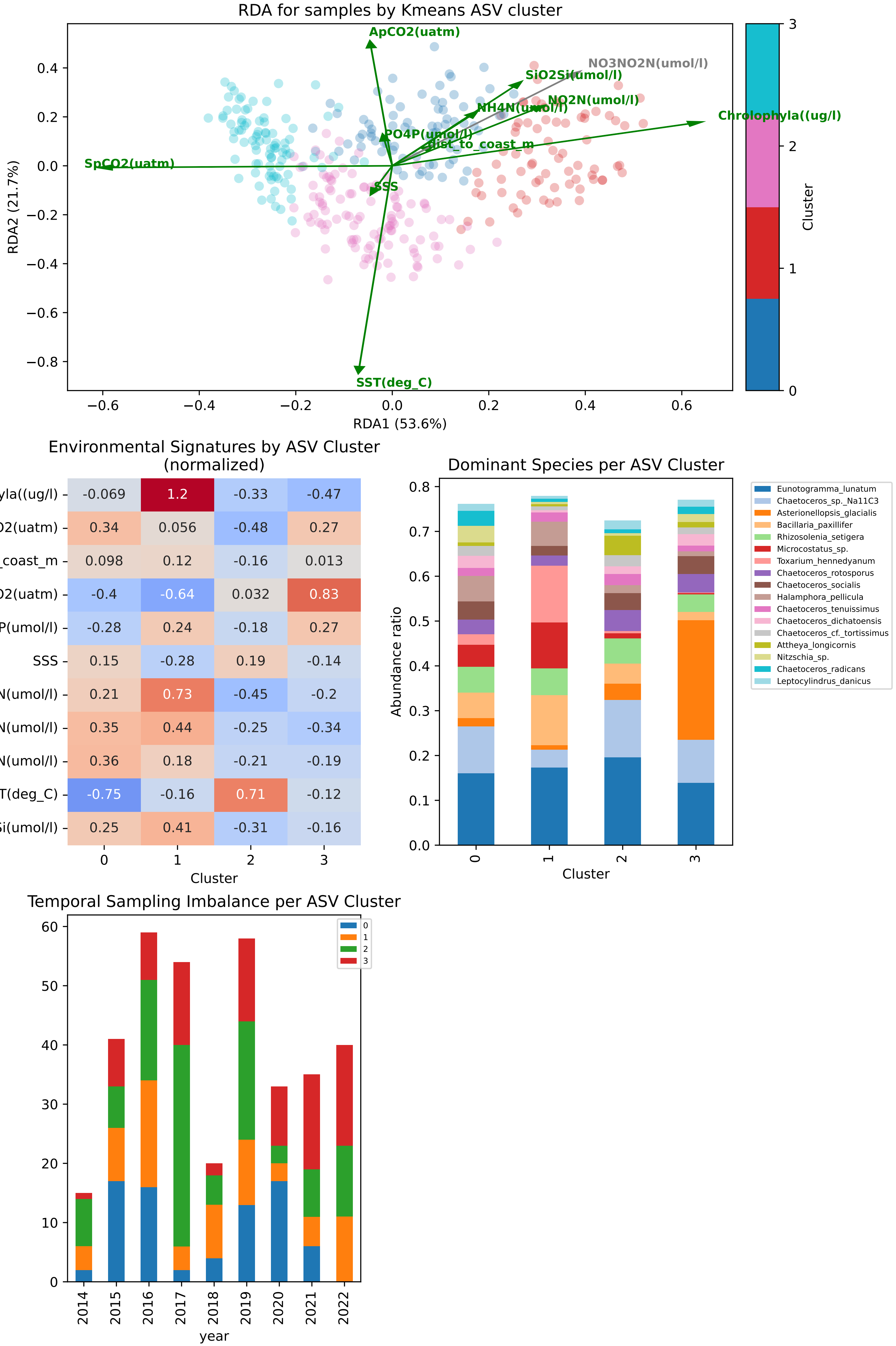

RDA for samples by Kmeans ASV cluster
ApCO2(uatm)
NO3NO2N(umol/l)
SiO2Si(umol/l)
NO2N(umol/l)
NH4N(umol/l)
Chrolophyla((ug/l)
PO4P(umol/l)
dist_to_coast_m
SpCO2(uatm)
SSS
SST(deg_C)
RDA2 (21.7%)
RDA1 (53.6%)
Cluster
Environmental Signatures by ASV Cluster (normalized)
Chrolophyla((ug/l) -0.069 1.2 -0.33 -0.47
ApCO2(uatm) 0.34 0.056 -0.48 0.27
dist_to_coast_m 0.098 0.12 -0.16 0.013
SpCO2(uatm) -0.4 -0.64 0.032 0.83
PO4P(umol/l) -0.28 0.24 -0.18 0.27
SSS 0.15 -0.28 0.19 -0.14
NO3NO2N(umol/l) 0.21 0.73 -0.45 -0.2
NO2N(umol/l) 0.35 0.44 -0.25 -0.34
NH4N(umol/l) 0.36 0.18 -0.21 -0.19
SST(deg_C) -0.75 -0.16 0.71 -0.12
SiO2Si(umol/l) 0.25 0.41 -0.31 -0.16
Cluster
Dominant Species per ASV Cluster
Abundance ratio
Cluster
Eunotogramma_lunatum
Chaetoceros_sp._Na11C3
Asterionellopsis_glacialis
Bacillaria_paxillifer
Rhizosolenia_setigera
Microcostatus_sp.
Toxarium_hennedyanum
Chaetoceros_rotosporus
Chaetoceros_socialis
Halamphora_pellicula
Chaetoceros_tenuissimus
Chaetoceros_dichatoensis
Chaetoceros_cf._tortissimus
Attheya_longicornis
Nitzschia_sp.
Chaetoceros_radicans
Leptocylindrus_danicus
Temporal Sampling Imbalance per ASV Cluster
year

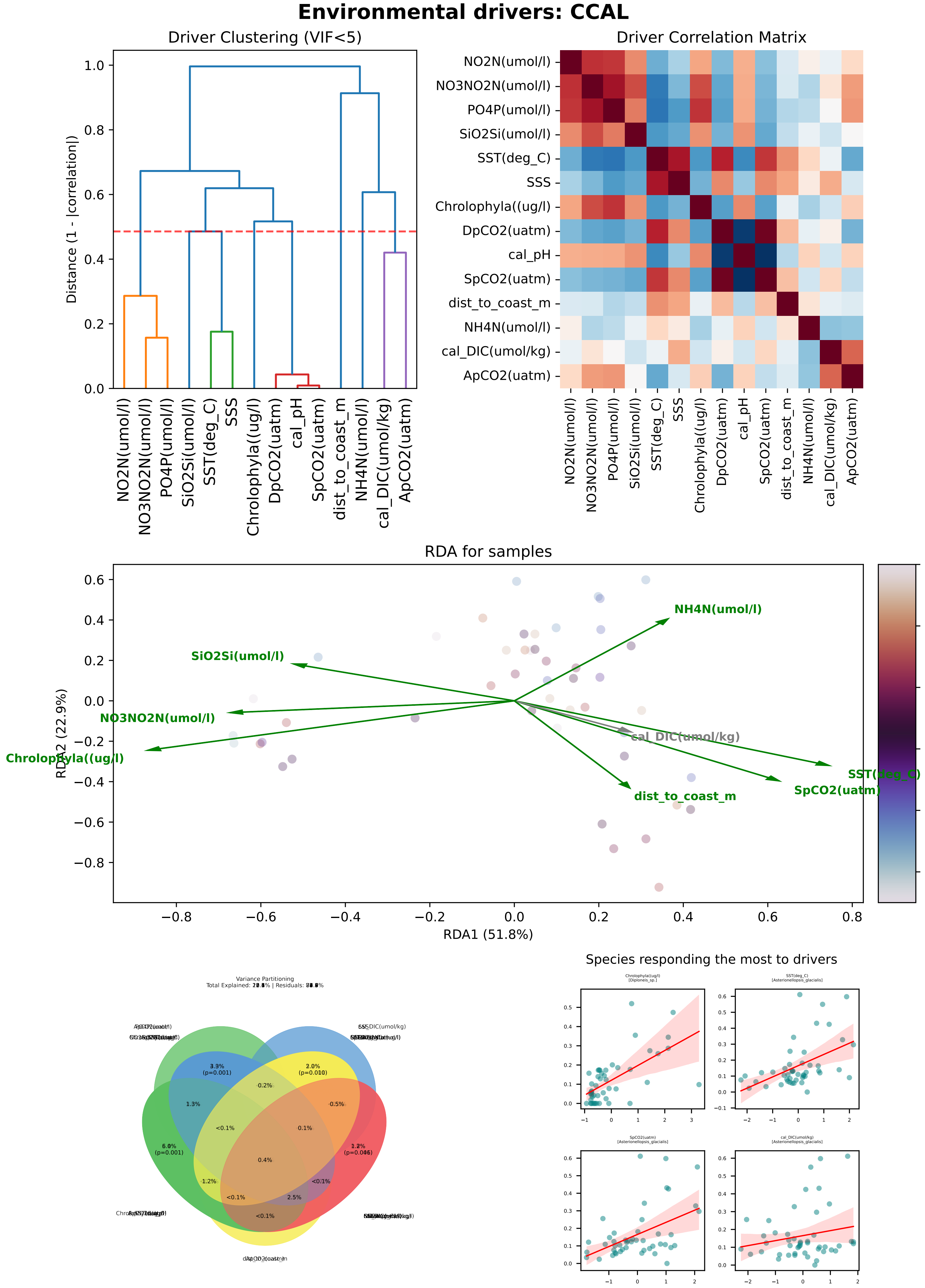
Environmental drivers: CCAL
Driver Clustering (VIF<5)
Distance (1 - |correlation|)
NO2N(umol/l)
NO3NO2N(umol/l)
PO4P(umol/l)
SiO2Si(umol/l)
SST(deg_C)
SSS
Chrolophyla((ug/l)
DpCO2(uatm)
cal_pH
SpCO2(uatm)
dist_to_coast_m
NH4N(umol/l)
cal_DIC(umol/kg)
ApCO2(uatm)
Driver Correlation Matrix
RDA for samples
RDA1 (51.8%)
RDA2 (22.9%)
Variance Partitioning
Species responding the most to drivers
Chrolophyla((ug/l) [Diploneis_sp.]
SST(deg_C) [Asterionellopsis_glacialis]
SpCO2(uatm) [Asterionellopsis_glacialis]
cal_DIC(umol/kg) [Asterionellopsis_glacialis]

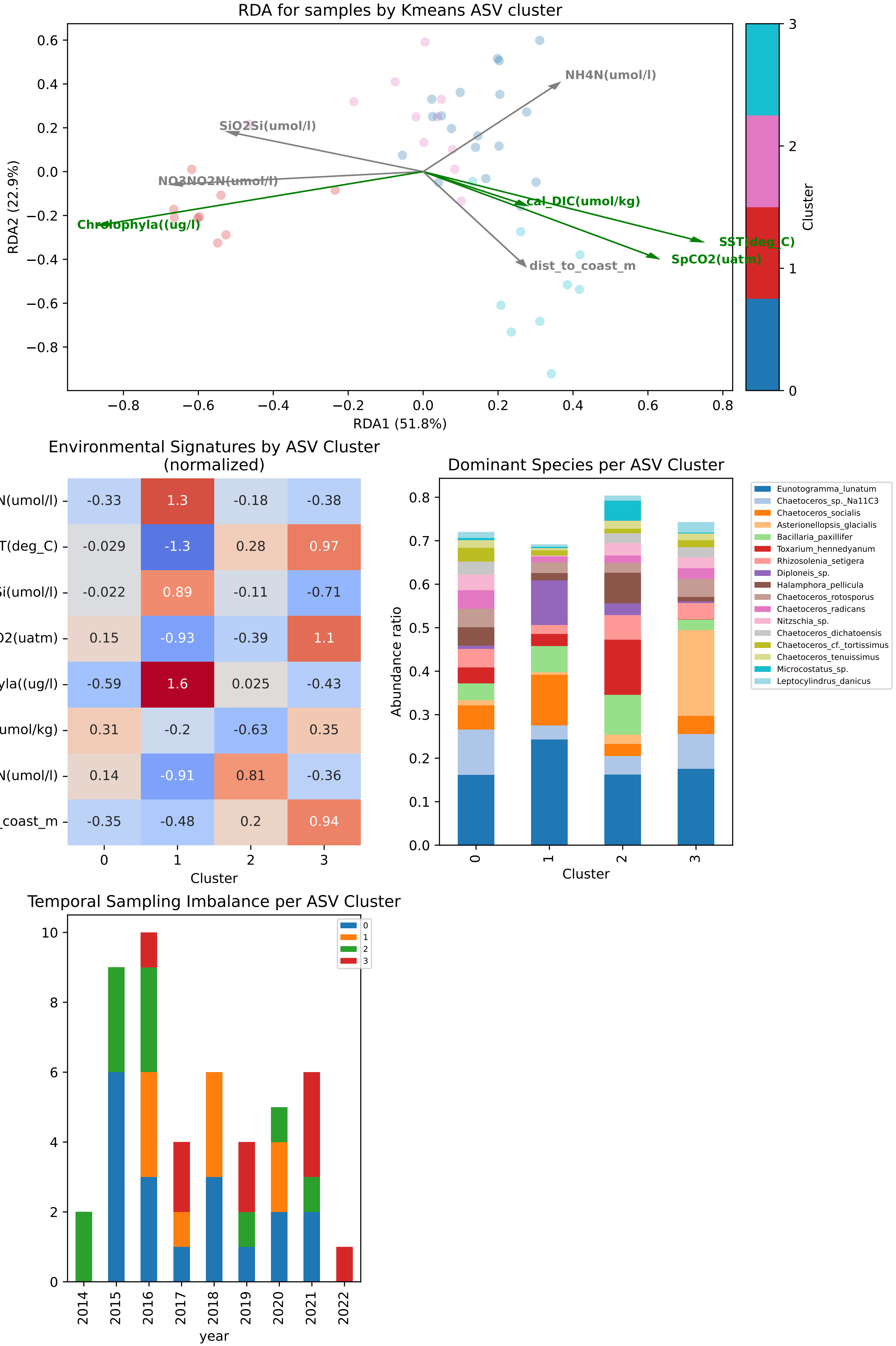

RDA for samples by Kmeans ASV cluster
RDA1 (51.8%)
RDA2 (22.9%)
NH4N(umol/l)
SiO2Si(umol/l)
NO3NO2N(umol/l)
Chrolophyla((ug/l)
cal_DIC(umol/kg)
SST(deg_C)
SpCO2(uatm)
dist_to_coast_m
Cluster
Environmental Signatures by ASV Cluster
(normalized)
NO3NO2N(umol/l) -0.33 1.3 -0.18 -0.38
SST(deg_C) -0.029 -1.3 0.28 0.97
SiO2Si(umol/l) -0.022 0.89 -0.11 -0.71
SpCO2(uatm) 0.15 -0.93 -0.39 1.1
Chrolophyla((ug/l) -0.59 1.6 0.025 -0.43
cal_DIC(umol/kg) 0.31 -0.2 -0.63 0.35
NH4N(umol/l) 0.14 -0.91 0.81 -0.36
dist_to_coast_m -0.35 -0.48 0.2 0.94
Cluster
Dominant Species per ASV Cluster
Abundance ratio
Cluster
Eunotogramma_lunatum
Chaetoceros_sp._Na11C3
Chaetoceros_socialis
Asterionellopsis_glacialis
Bacillaria_paxillifer
Toxarium_hennedyanum
Rhizosolenia_setigera
Diploneis_sp.
Halamphora_pellicula
Chaetoceros_rotosporus
Chaetoceros_radicans
Nitzschia_sp.
Chaetoceros_dichatoensis
Chaetoceros_cf._tortissimus
Chaetoceros_tenuissimus
Microcostatus_sp.
Leptocylindrus_danicus
Temporal Sampling Imbalance per ASV Cluster
year

### Environmental drivers: PNEC

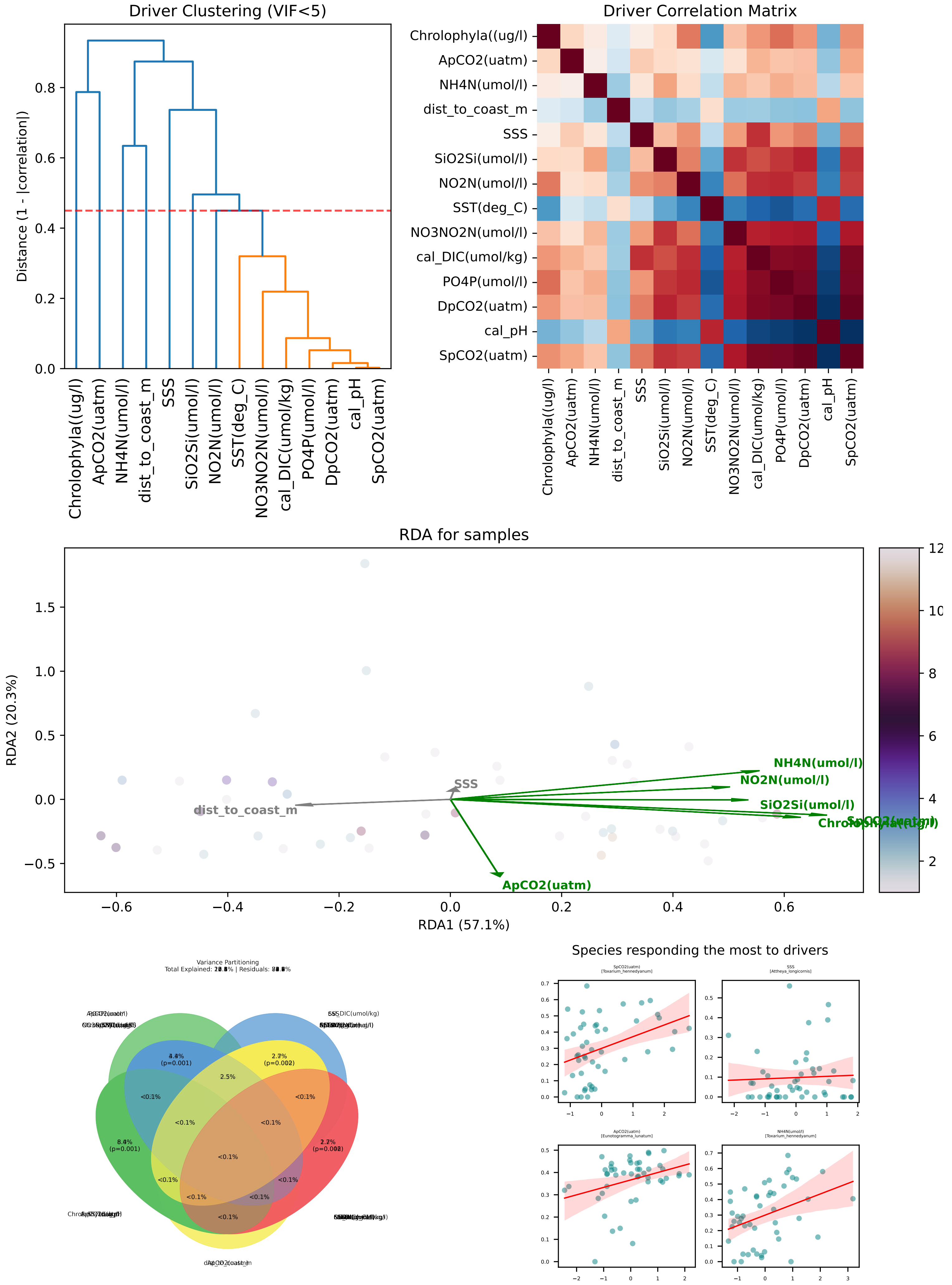

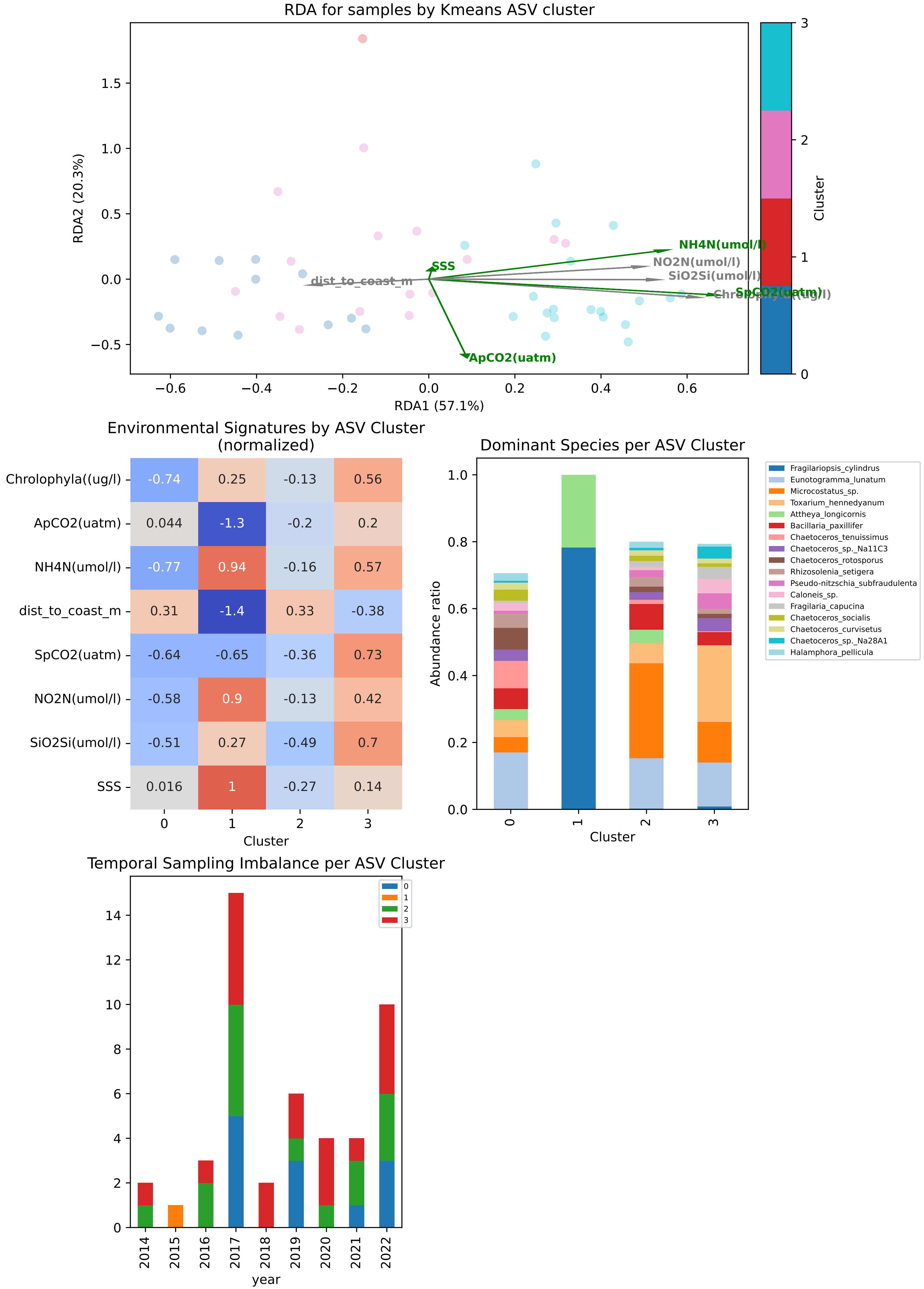
RDA for samples by Kmeans ASV cluster
RDA2 (20.3%)
RDA1 (57.1%)
Cluster
NH4N(umol/l)
NO2N(umol/l)
SiO2Si(umol/l)
SpCO2(uatm)
SSS
dist_to_coast_m
ApCO2(uatm)
Environmental Signatures by ASV Cluster (normalized)
Chrolophyla((ug/l) -0.74 0.25 -0.13 0.56
ApCO2(uatm) 0.044 -1.3 -0.2 0.2
NH4N(umol/l) -0.77 0.94 -0.16 0.57
dist_to_coast_m 0.31 -1.4 0.33 -0.38
SpCO2(uatm) -0.64 -0.65 -0.36 0.73
NO2N(umol/l) -0.58 0.9 -0.13 0.42
SiO2Si(umol/l) -0.51 0.27 -0.49 0.7
SSS 0.016 1 -0.27 0.14
Cluster
Dominant Species per ASV Cluster
Abundance ratio
Cluster
Fragilariopsis_cylindrus
Eunotogramma_lunatum
Microcostatus_sp.
Toxarium_hennedyanum
Attheya_longicornis
Bacillaria_paxillifer
Chaetoceros_tenuissimus
Chaetoceros_sp._Na11C3
Chaetoceros_rotosporus
Rhizosolenia_setigera
Pseudo-nitzschia_subfraudulenta
Caloneis_sp.
Fragilaria_capucina
Chaetoceros_socialis
Chaetoceros_curvisetus
Chaetoceros_sp._Na28A1
Halamphora_pellicula
Temporal Sampling Imbalance per ASV Cluster
year

### S8 ANCOM

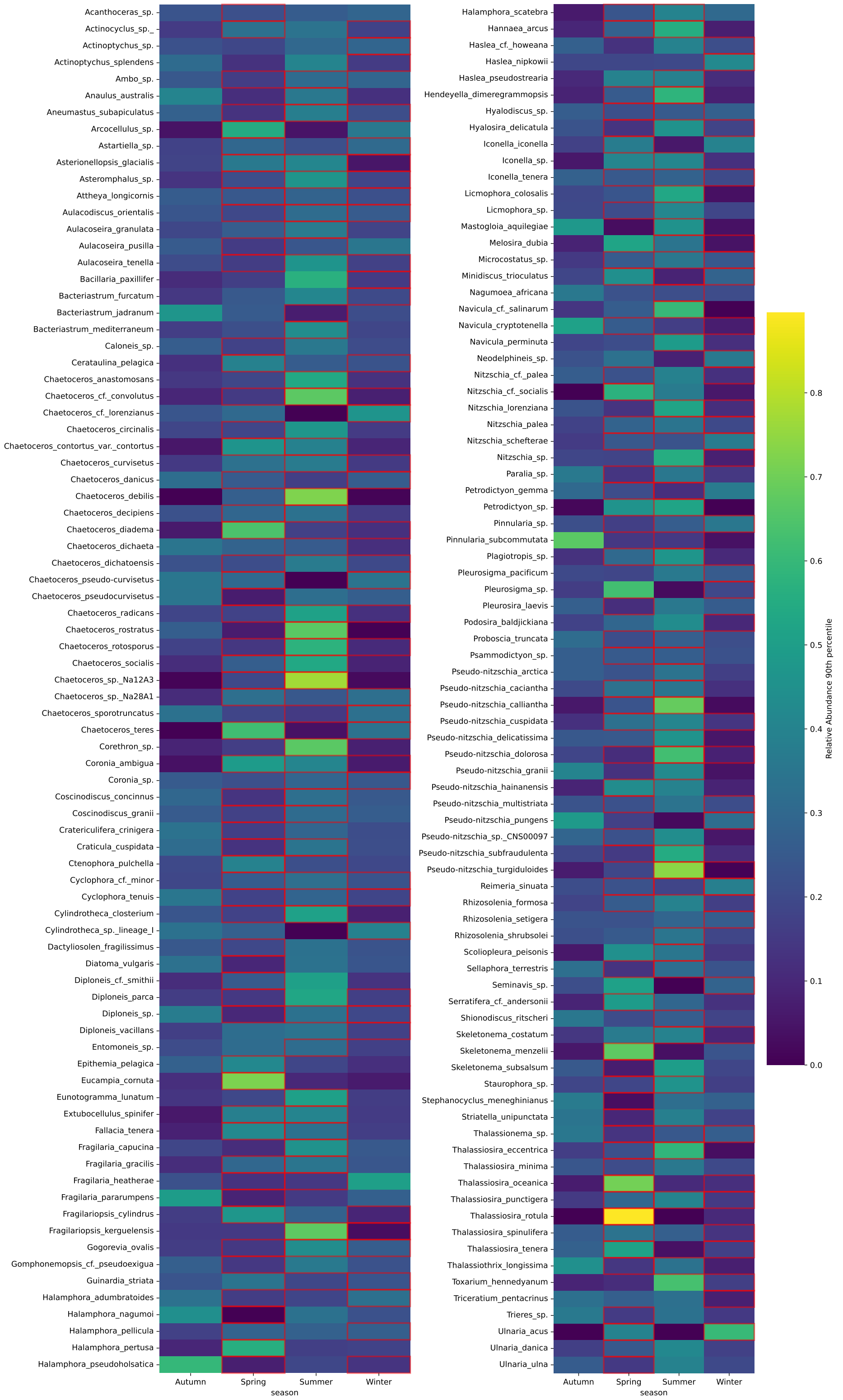


Figure 6: Seasonal distribution of differentially abundant taxa. Heatmap displaying taxa identified as significantly different across seasons using ANCOM-BC. Color intensity represents the species-normalized 90th percentile of nonzero relative abundance. Significance was determined using Benjamini-Hochberg adjusted p-values. Red outlines highlight the specific seasons for which each taxon was identified as significantly differentially abundant."
